# An intrinsic chromatin timer sets the tempo for development

**DOI:** 10.1101/2025.10.28.685222

**Authors:** Christopher Catalano, Michael S. Levine, Thomas W. Tullius

## Abstract

Embryonic patterning depends on both spatially localized developmental determinants and temporal competence of the regulatory genome to respond. Classical misexpression experiments revealed restricted time windows of regulatory competence, but the underlying molecular mechanisms remain unknown. Using single molecule whole genome methods, we identify hundreds of cis-regulatory modules (CRMs) that undergo a transition in chromatin accessibility during early *Drosophila* development. A special subset of these CRMs is transiently active only during gastrulation. Their detailed cis-regulatory architectures convert small changes in the balance between two pioneer factors, Zelda and GAF, into sharp temporal thresholds of chromatin competence. Remarkably, these “pulse” CRMs open and close on schedule without key regulatory determinants such as Dorsal (fly NF-kB) and Twist and in the absence of transcription. This intrinsic chromatin timer triggers global changes in transcriptional dynamics as cells exit pluripotency.

---

The specification of diverse cell types during development requires an invariant genome to make different regulatory decisions across time and space (*1*). Spatially restricted activators determine where developmental enhancers can act, but only after they acquire competence to respond. Previous misexpression experiments underscored this temporal requirement. The *hsp70* heat shock promoter and upstream regulatory sequences were used to misexpress a variety of different segmentation genes in early *Drosophila* embryos. Strong developmental defects appeared when they were induced during zygotic genome activation and gastrulation, but not after post-gastrular stages (*2, 3*). Temporal competence is a key property of the regulatory genome, and previous studies suggest that it is likely to depend on pioneer factors (*4–8*).

The pioneer factors Zelda and GAF/Trl are active during the 2–4 hour (h) interval of embryonic development when pluripotent nuclei enter distinct developmental programs at gastrulation (*9, 10*). Zelda preferentially controls the earliest wave of zygotic genome activation, whereas GAF progressively contributes as embryos enter gastrulation (*6, 10–15*). We used single-molecule footprinting to measure accessibility of regulatory chromatin, transcription-factor occupancy, and RNA polymerase II (Pol II) dynamics on millions of individual DNA molecules (*16–20*). Zelda- dominant cis-regulatory modules (CRMs) give way to GAF-dependent CRMs as development progresses. A special subset containing a mixture of Zelda and GAF motifs exhibits transient pulses of chromatin accessibility during gastrulation. These pulse CRMs open and close on schedule without transcription in mutant embryos lacking key regulatory determinants. We discuss the implications of this “pioneer clock” for the specification of lineage-restricted cell types from multipotent progenitors.

## Widespread potentiation of promoters and CRMs

We generated ultra-deep Fiber-seq datasets from staged *Drosophila* embryos (*16*), reaching nearly 2500-fold coverage with 19-kb median reads at both 2–4 and 6–8 h following fertilization. Individual fibers extend for tens of kilobases and frequently span multiple regulatory regions, promoters and transcription units. We used FiberHMM to identify protected footprints and intervening methyltransferase-sensitive patches (MSPs) on each molecule (*21*). Their shapes and genomic positions distinguish nucleosomes, transcription factors, preinitiation complexes (PICs), promoter-proximal paused Pol II, and elongating polymerases (Fig. 1A; fig. S1A–D). Each fiber connects the accessibility of regulatory chromatin with poised or active promoters and transcriptional output. We validated these operational states with public embryonic chromatin-binding and nascent-transcription datasets, including published early PRO-seq (*6, 10–12, 22*) (fig. S1E–H).

**Fig. 1.**
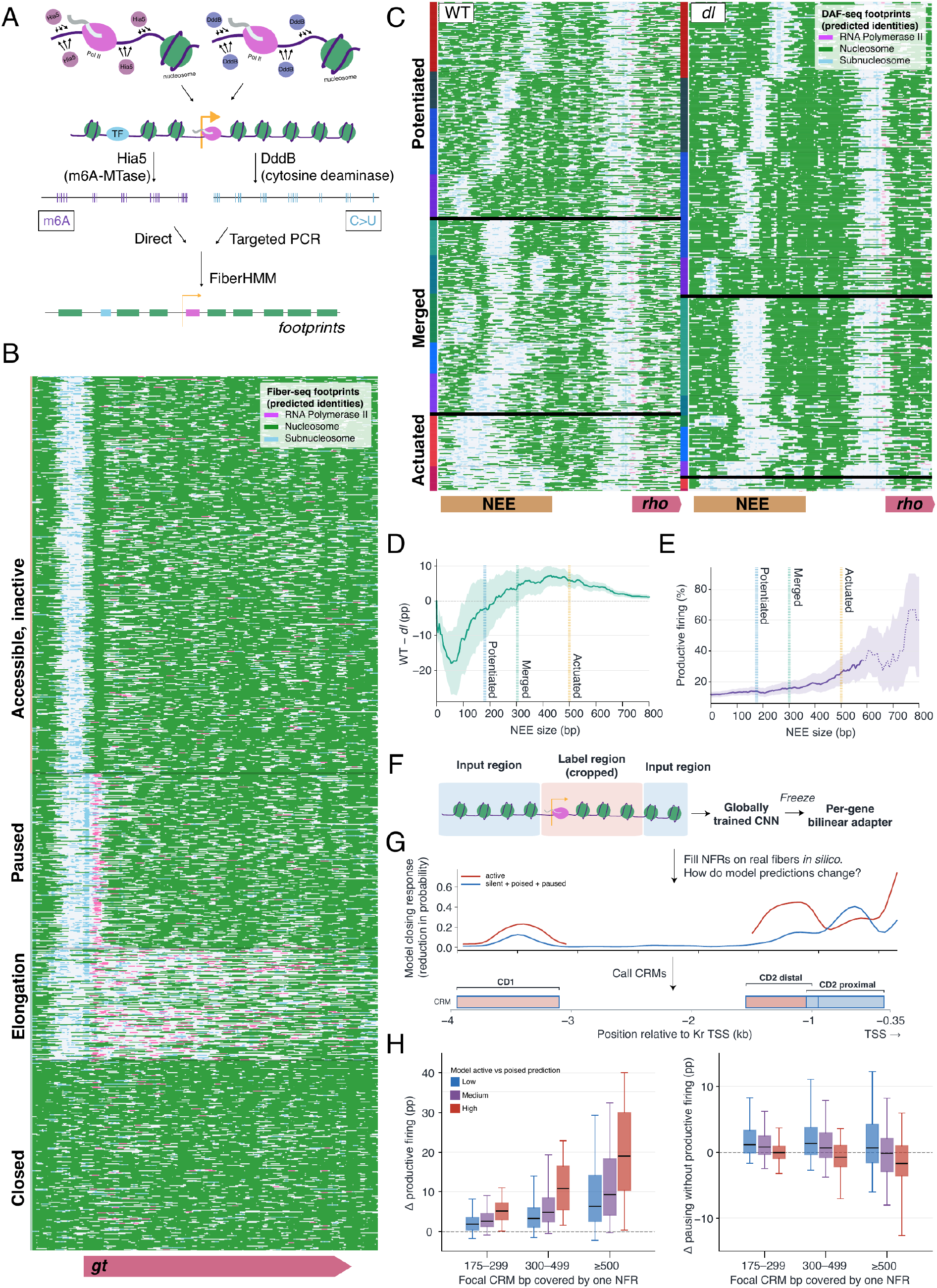
Single molecules distinguish potentiation from actuation and resolve CRMs. (**A**) Fiber-seq and Deaminase-Assisted Fiber-seq (DAF-seq) chemistry and FiberHMM interpretation. Hia5 methylates exposed adenines and DddB deaminates exposed cytosines; FiberHMM segments protected intervals and nucleosome-free regions (NFRs), with identities assigned from geometry and genomic context. (**B**) Representative WT 2–4-hour PacBio Fiber-seq molecules at *gt*, arranged by promoter/gene-body state. (**C**) WT and maternal *dorsal* null (*dl*) pooled 2–4-hour DAF-seq molecules across the *rhomboid* (*rho*) neuroectodermal enhancer (NEE), arranged by NEE opening and co-clustered by footprint pattern; colored blocks denote shared clusters. (**D**) WT − *dl*− NEE opening after genotype-matched, exact-width non-CRM background subtraction. Shading is a 95% interval combining molecule sampling with variation across six matched background intervals per genotype. (**E**) Productive-firing frequency versus NEE NFR size in WT PacBio molecules; dotted segments contain <50 fibers, and shading is a 95% confidence interval. (**F**) FiberCNN architecture: a shared local encoder is frozen and interpreted by a gene-specific adapter. (**G**) *Kr* computational-closing map; peaks in firing loss define candidate CRMs. (**H**) Changes in productive firing and pausing without productive firing relative to closed fibers as focal CRM opening broadens, stratified by low, medium, or high model active-versus-poised prediction. Boxes show medians/IQRs; whiskers, 5th–95th percentiles.

Promoter accessibility and engagement by PICs and paused Pol II are widespread, far more so than would be expected given the restricted limits of expression of most patterning genes (*23, 24*). The limited spatial profiles are instead revealed by the small fraction of fibers exhibiting productive Pol II elongation (Fig. 1B; fig. S1D,E). The levels of elongation on active fibers are highly variable, with the most active fibers often carrying closely spaced convoys of elongating polymerases and extensive nucleosome eviction that appears to persist during refractory periods of transcription (figs. S2A–D and S3A–G). Regulatory regions are similarly heterogeneous, spanning short, single-nucleosome openings, expanded evictions of ∼300 bp, and rarer nucleosome-free regions (NFRs) of 500 bp or more.

To test the association between NFR size and transcriptional output, we highlight the well-defined, Dorsal-dependent *rhomboid* (*rho*) neuroectodermal enhancer (NEE) (*25*), which displays the full range of potentiated and actuated states (Fig. 1C). Targeted Deaminase-Assisted Fiber-seq assays (DAF-seq) (*17, 26*) were used to compare wild-type embryos with those derived from *dl*/*dl* mothers. Mutant embryos lack both Dorsal and Twist, the principal activators of the *rho* NEE as well as many other developmental enhancers (*25*). Openings spanning approximately 175–300 bp are enriched in the mutant, whereas openings above ∼300 bp become progressively depleted (Fig. 1C,D; fig. S6A,B). Productive elongation across rho is also strongly reduced, as expected (Fig. 1C). In wild-type, the extent of NEE opening predicts transcriptional output: fibers carrying NFRs of at least 500 bp exhibit increased productive firing (Fig. 1E). The same ∼300-bp crossover point of potentiated and actuated CRMs is also observed for other Dorsal target genes such as *snail* (fig. S6C,D).

The persistence of small NFRs without Dorsal, coupled to the loss of broad opening and productive elongation, provides a clear distinction between widespread potentiation and rarer actuation. We used nested lower bounds of approximately 175 bp and 300 bp for potentiated CRMs, and 500 bp for actuated CRMs. By these criteria, most accessible regulatory DNA in the early embryo is potentiated but inactive, mirroring the many open or engaged promoters that are devoid of productive Pol II elongation.

### A single-molecule atlas for developmental enhancers and constituent CRMs

To gain a broader understanding of regulatory potentiation, we trained FiberCNN, a convolutional network model, to predict promoter engagement, pausing, elongation, and nucleosome eviction across individual fibers after masking their promoters and transcription units (Fig. 1F,G). For test set fibers, the model was able to distinguish different states of transcription. Because transcription is stochastic, the model predicts the probability of each state rather than the exact outcome on every molecule (fig. S4B–F). Consistent with the earlier analysis of *rho* regulation, the model assigned its largest impact on transcriptional output to broad NFRs (> 500 bp) associated with actuation (Fig. 1H). To create a comprehensive atlas of CRMs from the model, we closed NFRs computationally and measured the predicted consequence on transcription (Fig. 1G). This procedure mapped 12,121 candidate CRMs linked to 1,276 active genes, most within 10 kb of a transcription start site, and assigned functional consequences of these CRMs that are consistent with empirical observations on single fibers (Fig. 1F,G; figs. S4A–F and S5A–D; archived CRM catalog (*49*)). Annotated developmental enhancers frequently partition into multiple CRMs. For example, nearly two-thirds of all extended (2 kb and larger) REDfly enhancers (*27*) contain at least two CRMs (fig. S8A,E).

We consider the regulation of *snail* expression in greater detail due to its importance in epithelial-mesenchymal transitions (EMT) in a variety of developmental and disease processes in both flies and vertebrates (*28*). In flies, it is essential for the coordinated invagination of the mesoderm at the onset of gastrulation (*29*). The *snail* proximal enhancer resolves into three separate modules: P1, P2, and P3. Approximately 60% of the fibers containing actuated P3 exhibit productive Pol II elongation, compared with only ∼15% for those containing actuated P1 modules. The distal (or shadow) enhancer likewise separates into high-output D1 and low-output D2, which appears to reduce productive elongation when actuated (Fig. 2A–C; figs. S6C,D and S13A,B). These assignments align with previous genetic studies (*30–34*). P2 closely matches a recently identified *snail* autorepression element, whereas D2 coincides with binding of both Snail and unidentified spatial silencers (*31, 34*).

**Fig. 2.**
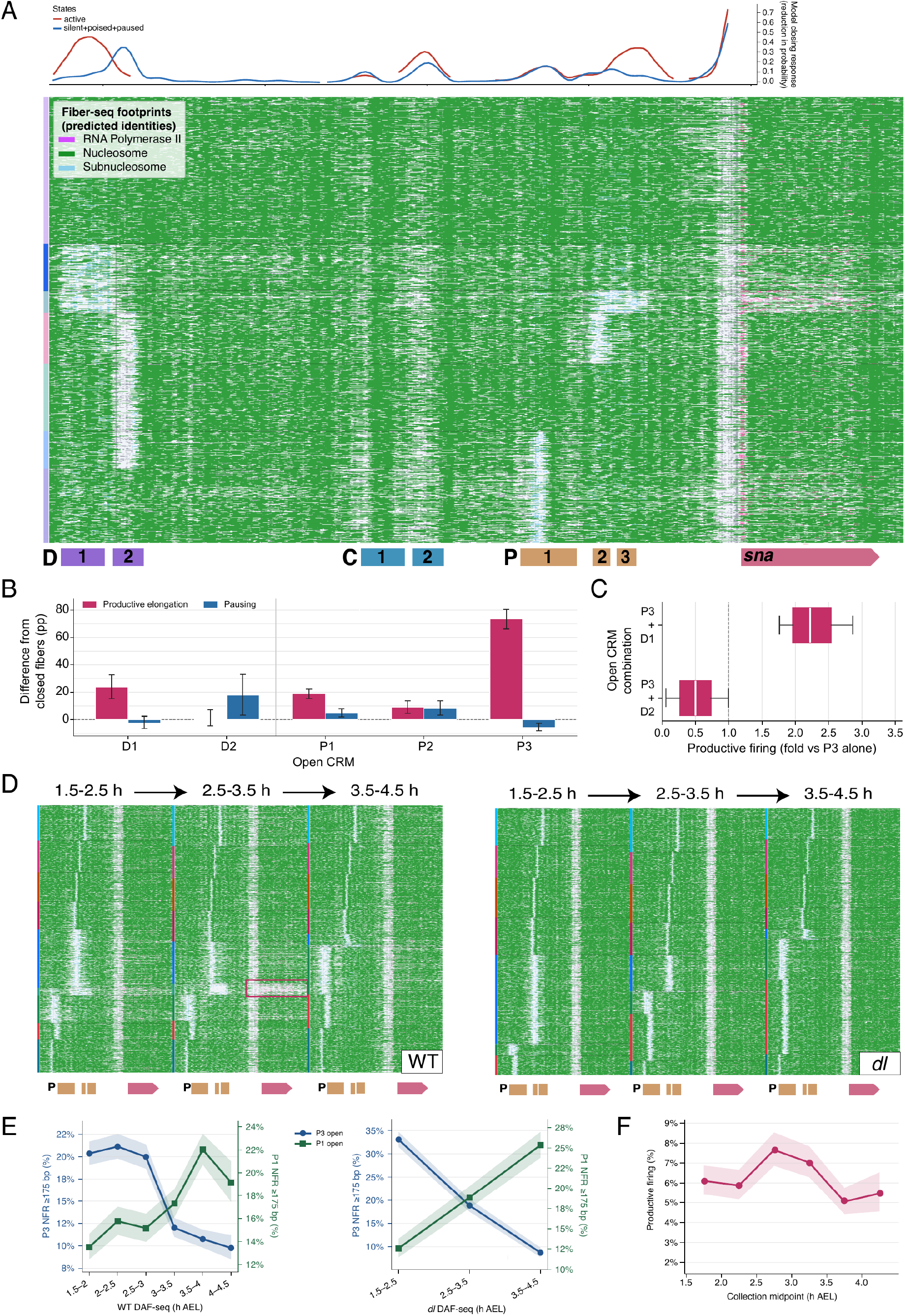
A developmental timer within a single enhancer. (**A**) *snail* (*sna*) computational-closing profiles and representative WT 2–4-hour PacBio Fiber-seq molecules. Red denotes model support for productive transcription and blue pooled silent, poised, and paused states; distal D1/D2, central C1/C2, and proximal P1/P2/P3 CRMs are indicated. (**B**) Excess productive elongation and pausing relative to closed fibers for molecules carrying an isolated ≥500-bp opening centered on each CRM. Error bars are library-stratified molecule-bootstrap 95% confidence intervals. (**C**) Productive-firing rate on fibers with P3 co-actuated with D1 or D2, relative to P3 alone. (**D**) WT and maternal *dl* targeted DAF-seq molecules across the *sna* proximal enhancer and transcription unit in three 1-hour windows from 1.5–4.5 hours after egg laying (AEL). WT 1-hour views pool adjacent 30-min collections. Molecules are co-clustered across samples by footprint pattern; colored blocks denote shared clusters. The WT 2.5–3.5-hour high-activity cluster is boxed. (**E**) P3 and P1 ≥175-bp NFR frequencies across six 30-min WT collections and three 1-hour *dl* collections; ribbons are 95% Wilson intervals. (**F**) Productive- firing frequency across the complete *sna* transcription unit in the WT DAF-seq time course; ribbon, 95% Wilson interval.

Such repressor elements are hard to identify with activation-based reporter assays but readily emerge from their same-fiber relationship to transcription. A more common mode of repression is seen for the *zen* ventral repression element (VRE) (*35*), whereby Dorsal restricts merging and actuation of an otherwise highly productive CRM (fig. S7A–D).Globally, independently actuated CRM pairs are usually additive rather than sub- or super-additive. A fraction nevertheless shows synergistic interactions, including 96 repressor-like pairs where one CRM overrides the output of the other, like D1 and D2 (fig. S9B–M). While clearly synergistic interactions appear rare and require high sequencing depth to call confidently, a large subset of CRM pairs often actuates coordinately, providing a more widespread mechanism for regulatory cooperativity (fig. S10).

### A developmental timer within a single enhancer

Past studies documented intense levels of *snail* transcription during early periods of zygotic genome activation, but as embryos approach gastrulation there is a marked reduction in the level of expression due in part to autorepression (*34, 36*). Interestingly, we found that in the 2–4 h dataset P1 and P3 had strikingly different relationships to *snail* transcription. After subtraction of a matched closed background, actuated P1 NFRs were found to be associated with less productive elongation and more pausing than comparable actuated P3 NFRs (Fig. 2B). These stronger P3 and weaker P1 configurations largely occupy different fibers. Because the 2–4 h collection spans zygotic genome activation through gastrulation, the temporal sequence of P3 and P1 activity was uncertain.

We therefore used targeted DAF-seq to follow the *snail* proximal enhancer and associated transcription at half-hour resolution (Fig. 2D,E; fig. S13B). Productive firing across *snail* rises and then declines as embryos enter gastrulation (Fig. 2F). The shifting potentiation of P1 and P3 over time revealed a corresponding exchange. The fraction of fibers carrying potentiated openings anywhere within the P1–P3 interval remains broadly stable, with a small peak at 2.5–3 h, but the distribution of those openings shifts sharply over time (fig. S13B). P3 displays a two-fold reduction in chromatin opening, whereas P1 rises and becomes the dominant proximal CRM at the onset of gastrulation at 3 h (Fig. 2E; fig. S13B). The 2–4 h dataset therefore combines an early, highly productive P3 program with a later, less productive P1 program. CRM switching is associated with a downshift in transcriptional dynamics (see below).

To determine whether this switch from P3 to P1 opening depends on transcriptional activation, we examined embryos derived from *dl*/*dl* mothers. As expected, no bona fide elongating Pol II is detected across the *snail* transcription unit in these dorsalized embryos. As at *rho* (Fig. 1D), we observe a concomitant loss of actuated P1 and P3 NFRs and an increase in potentiated 175–300 bp P1 and P3 NFRs (Fig. 2D,E; figs. S6C and S13C). Nevertheless, P3 opening declines and P1 opening rises on schedule (Fig. 2E; fig. S13C). These observations suggest that the P3 to P1 switch is independent of transcription and spatial activators.

### Pulses of chromatin potentiation during gastrulation

P3 and P1 contain distinct pioneer-factor motifs. P3 carries several high-affinity Zelda motifs, whereas P1 contains lower-affinity Zelda and GAF motifs. We therefore asked whether the P3 to P1 switch was part of the broader Zelda-to-GAF transition during early development. The 2–4 h and 6–8 h Fiber-seq datasets recover the expected global shift from Zelda- to GAF-associated CRMs (*10*). P1, however, declines by 6–8 h, indicating that its rise during the DAF-seq time course is transient.

We generated additional Fiber-seq datasets from 1.5–3 h and 3–4.5 h wild-type embryos to identify related trajectories genome-wide. Combining these measurements with the 2–4 h to 6–8 h transition allowed us to identify 104 early, 245 pulse, and 240 late CRMs (Fig. 3A,B; fig. S14A,H,I; archived temporal-class source data (*49*)). These three classes highlight dynamic changes in CRM opening across a continuous temporal landscape. Early CRMs lose potentiation after 1.5–3 h, late CRMs rise across the entirety of the time course, and pulse CRMs open transiently during gastrulation before declining by 6–8 h. The temporal signal is strongest at the 175-bp threshold, placing it within widespread potentiated chromatin rather than the rarer actuated state associated with productive transcription (fig. S14F). Thus, the *snail* P3-to-P1 switch is part of a pulse within the broader Zelda-to-GAF transition (*4, 10*).

**Fig. 3.**
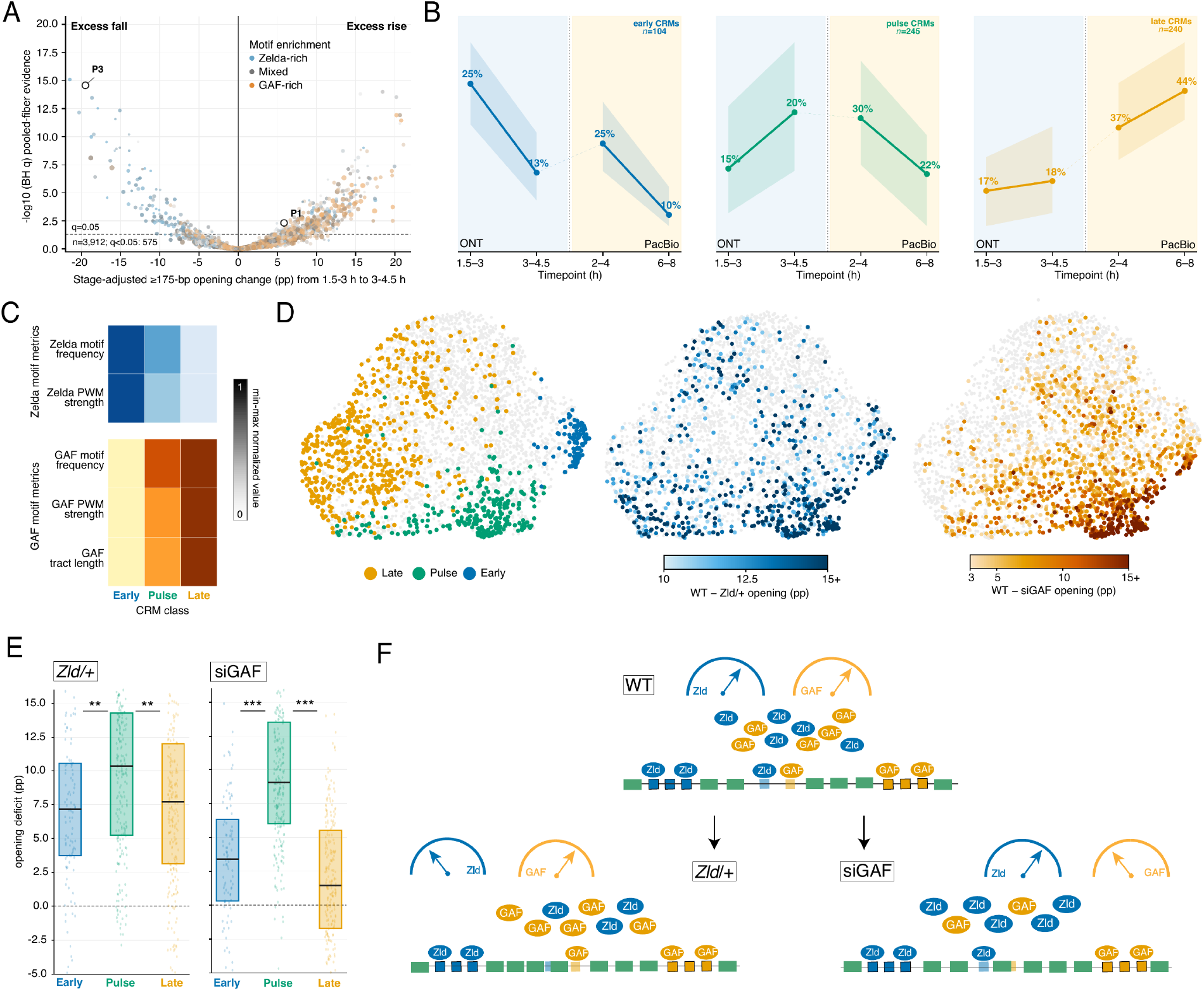
A gastrulation pulse from pioneer-factor balance. (**A**) Stage-adjusted change in ≥175- bp opening from WT 1.5–3 to 3–4.5 hours. Color denotes GAF-versus-Zelda motif balance. Of 3,912 CRMs with a widest boundary ≥300 bp and ≥30 callable fibers at each endpoint, 575 pass pooled-fiber Benjamini–Hochberg *q* < 0.05; *sna* P1 and P3 are labeled. (**B**) Median WT opening trajectories with interquartile bands for 104 early, 245 pulse, and 240 late CRMs. The 1.5–3 h and 3–4.5 h measurements are ONT Fiber-seq; 2–4 h and 6–8 h measurements are PacBio Fiber-seq. The dotted segment marks the assay handoff. (**C**) Row-normalized Zelda and GAF motif frequency/strength and GAF tract length by temporal class. Motifs were scanned on both strands within empirical CRM boundaries using matched JASPAR vfl/Trl matrices at MOODS *P* ≤ 10^−3^. (**D**) Fixed embedding of temporal opening and sequence-predicted accessibility for CRMs in (**A**). Left, temporal classes; middle and right, WT −− *zld*/+ and WT −− siGAF opening deficits in matched 3–4.5-hour ONT collections, all projected without refitting. (**E**) Opening deficits after reduced Zld dosage or GAF depletion. Boxes show medians and interquartile ranges; points denote CRMs. Pairwise *P* values use chromosome-stratified, gene-unioned 1-Mb block bootstraps; **, *P* < 0.01; ***, *P* < 0.001. (**F**) Schematic of changing pioneer-factor balance in *zld*/+ and siGAF embryos relative to WT.

Pulse CRMs occupy an intermediate temporal profile and pioneer binding regime, with mixed Zelda and GAF motif content and sequence composition signatures that distinguish them from early and late CRMs (Fig. 3B,C; fig. S14D–G; fig. S15). GAF depletion during the pulse window reduces their potentiation more strongly than that seen for early (Zelda-dominant) CRMs or late (GAF-dominant) CRMs. Reduced maternal Zelda dosage also weakly affects pulse CRMs (Fig. 3D,E; fig. S14B,C). Their mixed architecture and sensitivity to both perturbations are consistent with a model in which changing Zelda and GAF availability defines a transient window of competence (*37, 38*) (Fig. 3F).Fine-resolution DAF-seq at additional loci places the early-to-pulse switch near the onset of gastrulation, at approximately 3 h (Figs. 2D–F and 4D–G; fig. S18).

### Temporal identity shapes transcriptional output

At *snail*, the P3-to-P1 switch coincides with a shift to lower levels of elongating polymerases (Fig. 2B). Across CRM–gene associations, actuation of pulse CRMs is associated with less productive elongation than actuation of early CRMs (Fig. 4A). Considering only those genes containing both classes, we found that nearly 80% show a difference in the levels of transcription between early-only and pulse-only states. Of these, ∼80% exhibit lower productive elongation in the pulse-only state, whereas ∼20% show an increase (Fig. 4B; figs. S16–S18). Temporal shifts in CRM identity therefore influence transcriptional dynamics, usually downward but sometimes upward.

**Fig. 4.**
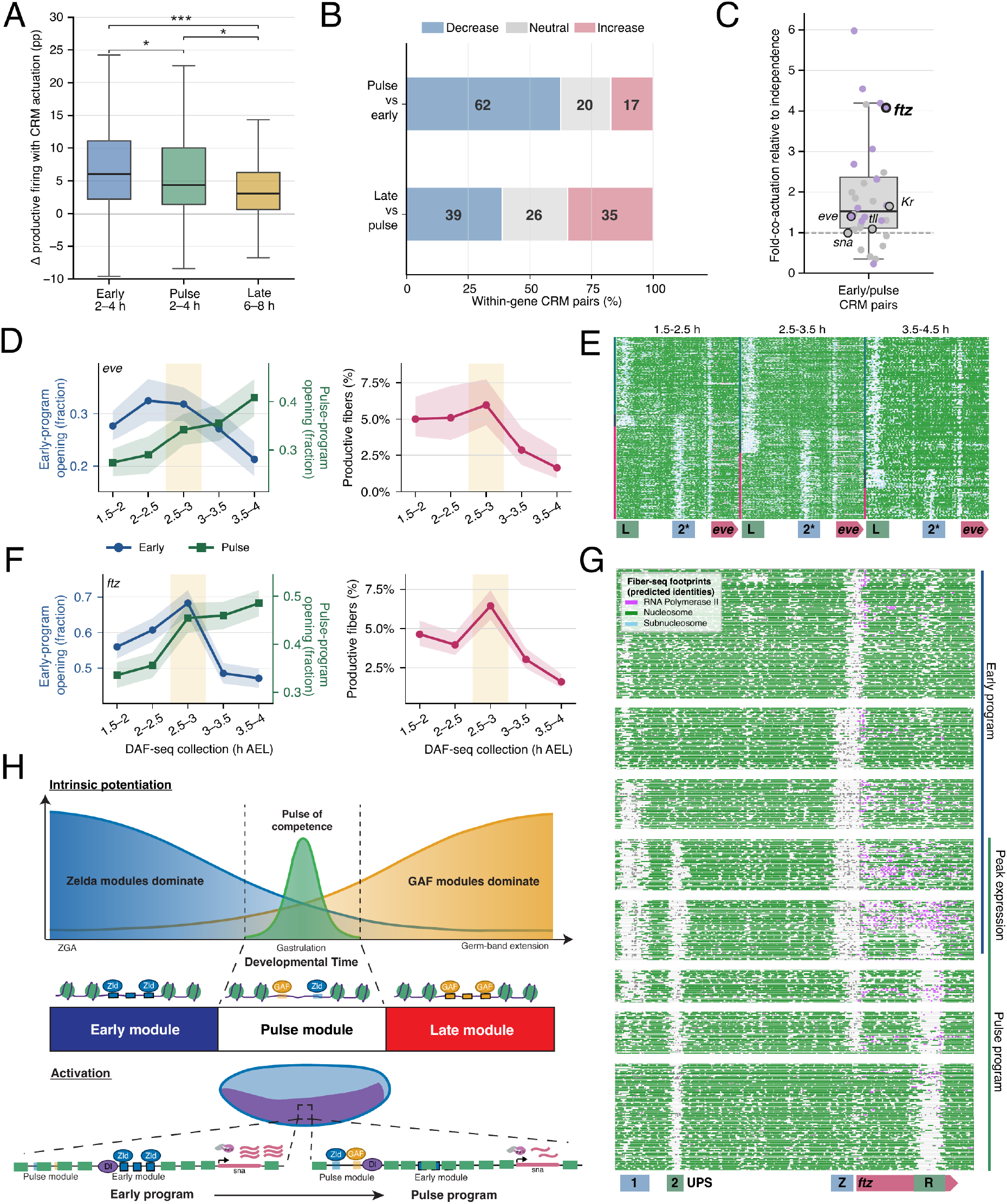
Temporal identity shapes transcriptional output. (**A**) Change in productive-firing frequency associated with CRM actuation in WT PacBio Fiber-seq, comparing fibers carrying a peak-containing ≥500-bp CRM NFR with each linked gene’s overall firing rate (early/pulse/late: 118/277/292 CRM–gene contexts). Boxes show median/IQR; whiskers, 1.5× IQR; brackets, Holm-adjusted two-sided Wald tests with gene-clustered errors. (**B**) Firing differences between mutually exclusive CRM states linked to the same gene: pulse versus early at 2–4 hours (64 pairs) and late versus pulse at 6–8 hours (72 pairs). Differences <−2.5or >+2.5 percentage points are decreases or increases; others are neutral. (**C**) Same-fiber early–pulse co-actuation relative to independence in WT 2–4-hour Fiber-seq; points are gene-level median observed/expected ratios (32 genes, 63 pairs). (**D**) WT targeted DAF-seq *eve* early- and pulse-program opening and productive-firing trajectories. (**E**) Co-clustered WT *eve* molecules across three 1-hour windows from 1.5–4.5 hours AEL. (**F**) Corresponding *ftz* trajectories for early (Z, UPS1) and pulse (R, UPS2) programs. Program opening in (D,F) is the fraction of jointly callable fibers with ≥1 constituent CRM carrying a ≥175-bp NFR; ribbons are 95% Wilson intervals and shading marks2.5–3 hours. (**G**) *ftz* molecules grouped into early, joint early/pulse (“peak expression”), and pulse states. (**H**) Model of a transient competence pulse generated by declining Zelda-and rising GAF-dominated modules.

Specific pairs of early and pulse CRMs show elevated co-actuation, suggesting a cooperative function between the two programs (Fig. 4C; figs. S10 and S11). We therefore generated 30-min DAF-seq time courses for *eve, ftz, tll* and *Kr*. At each locus, early and pulse potentiation overlap maximally at 2.5–3 h, coinciding with a local maximum in productive elongation, although the trajectories differ among genes (Fig. 4D–G; fig. S18). The maximum is sharpest for *ftz* (*39*), where the two programs also show coordinated co-actuation (Fig. 4F,G). Timed CRM succession often changes the transcriptional dynamics of key developmental control genes, and their temporal overlap can produce a transient output maximum that is amplified by coordination at particular genes such as *ftz*.

## Discussion

Our study identifies a developmental timer that produces sequential changes in potentiated chromatin in response to subtle changes in the relative levels of two pioneer factors, Zelda and GAF (Fig. 4H). Previous studies suggest a broad Zelda-to-GAF transition during zygotic genome activation (*10, 14, 37*). The discovery of pulse CRMs reveals a previously unappreciated response within this broader transition. We propose that transient potentiation of pulse CRMs is driven by an intrinsic developmental timer, which primes regulatory chromatin for timely activation. A remarkable property of this timer is that it is independent of transcription and highly tunable. We identify over 200 pulse CRMs that are transiently accessible during just a 1–2 h period during gastrulation. They are found in the regulatory regions of some of the most critical patterning genes in *Drosophila*, including a number of segmentation genes such as *eve, ftz*, and *Krüppel* as well as dorsal-ventral determinants like *snail, twist*, and *rho* (archived temporal-class source data (*49*)). In most cases, the switch from early Zelda-dominant CRMs to GAF-associated pulse CRMs results in a downshift in the levels of elongating polymerases as embryos enter gastrulation. There is evidence that attenuated expression of *snail* facilitates coordinated invagination of the mesoderm (*34*).

Pulse CRMs contain a series of suboptimized binding motifs that exhibit dosage-sensitive chromatin potentiation upon reductions in the levels of either Zelda or GAF. This mechanism resembles the interpretation of morphogen gradients in space (*40, 41*). Sequence-encoded thresholds convert graded spatial inputs into discrete expression domains, while pioneer factor motif architecture converts temporal changes in the levels of pioneer activity into bounded time-windows of developmental competence. At *snail*, the P3-to-P1 switch persists in embryos lacking maternal Dorsal despite the loss of enhancer actuation and productive transcription. An embryo that never makes mesoderm nonetheless runs the regulatory schedule for making it. Developmental time is therefore written into potentiated chromatin before it is read out in space.

Sequential pioneer-factor environments recur across development. During differentiation of human pluripotent stem cells toward definitive endoderm, GATA6 binding precedes and is required for FOXA2 recruitment and chromatin opening at shared regulatory elements (*42*). In mouse preimplantation embryos, OBOX promotes zygotic genome activation and induces *Nr5a2*, which later cooperates with KLF5 and GATA6 to regulate chromatin and transcription (*43–45*). Our findings predict that mixed-affinity CRMs can be used to form transient windows of regulatory competence wherever such regimes overlap. We suggest that even subtle changes in the relative levels of these pioneer factors could control the timing of key developmental transitions as seen for the pioneer clock in *Drosophila*.

## Acknowledgments

The authors would like to thank Jie Hu for purifying DddB enzyme; Andrew Stergachis for providing Hia5 enzyme; Bomyi Li and Christine Rushlow for providing fly lines; Bernard Kim and his research group for use of their PromethION sequencers; the University of Washington PacBio sequencing core for PacBio sequencing services; Xiaoliang Sunney Xie for providing the plasmid for transgenic expression of DddB; Cara Weisman for use of her MinION sequencer and reagents for ONT sequencing; Elizabeth Gavis and her research group for providing *yw* flies and miscellaneous materials; Alana Bernys and Zoë Feder for many helpful discussions; and all members of the Levine lab for countless discussions and insights throughout this project.

## Funding

National Institutes of Health grant R35 GM118147 (MSL)

## Author contributions

Conceptualization: CC, MSL, TWT

Methodology: CC, MSL, TWT

Investigation: CC, TWT

Visualization: CC, TWT

Funding acquisition: MSL

Project administration: MSL, TWT

Supervision: MSL, TWT

Writing – original draft: CC, MSL, TWT

Writing – review & editing: CC, MSL, TWT

## Competing interests

Authors declare that they have no competing interests.

## Data, code, and materials availability

Processed sequencing data have been deposited in the Gene Expression Omnibus, and raw sequencing data in the Sequence Read Archive; accession numbers will be provided upon publication. FiberHMM is publicly available at https://github.com/fiberseq/FiberHMM, with the manuscript-facing release archived in Zenodo (*47*). Archived manuscript releases of FiberCNN (*46*), FiberBrowser (*48*), and the analysis code and source data (*49*) are also deposited in Zenodo. The associated live repositories are linked from each Zenodo record. The archived analysis release includes the CRM catalog, processed source-data tables, data dictionaries, and figure-source manifests (*49*). The GEO, SRA, and Zenodo records will be made public upon publication. Materials are available from the corresponding authors upon reasonable request.

## Supplementary Materials

Materials and Methods

Supplementary Text

Figs. S1 to S18

Tables S1 to S9

References (50–59; cited only in the supplementary materials)

## Notes

### Competing Interest Statement

TWT is an inventor on a provisional patent application related to Fiber-seq applied to human sperm chromatin. This application is unrelated to the current study.

### Summary of Updates

This version substantially revises and expands the manuscript, with a new title and central biological framework. We added extensive new developmental time-course, perturbation, and single-molecule analyses that identify distinct temporal classes of cis-regulatory elements and support a pioneer-factor-driven chromatin timer controlling developmental windows of regulatory competence. The manuscript has been reorganized around these findings, with expanded genome-wide analyses and targeted validation experiments. Results that formed the main focus of the previous version have been retained in the Supplementary Information where relevant.

## References

1. E. Z. Kvon, T. Kazmar, G. Stampfel, J. O. Yáñez-Cuna, M. Pagani, K. Schernhuber, B. J. Dickson, A. Stark, Genome-scale functional characterization of Drosophila developmental enhancers in vivo. Nature 512, 91–95 (2014).

2. G. Struhl, Near-reciprocal phenotypes caused by inactivation or indiscriminate expression of the Drosophila segmentation gene ftz. Nature 318, 677–680 (1985).

3. S. J. Poole, T. B. Kornberg, Modifying expression of the engrailed gene of Drosophila melanogaster. Development 104, 85–93 (1988).

4. S. A. Blythe, E. F. Wieschaus, Establishment and maintenance of heritable chromatin structure during early Drosophila embryogenesis. eLife 5, e20148 (2016).

5. J. Dufourt, A. Trullo, J. Hunter, C. Fernandez, J. Lazaro, M. Dejean, L. Morales, S. Nait-Amer, K. N. Schulz, M. M. Harrison, C. Favard, O. Radulescu, M. Lagha, Temporal control of gene expression by the pioneer factor Zelda through transient interactions in hubs. Nat. Commun. 9, 5194 (2018).

6. K. J. Brennan, M. Weilert, S. Krueger, A. Pampari, H. Liu, A. W. H. Yang, J. A. Morrison, T. R. Hughes, C. A. Rushlow, A. Kundaje, J. Zeitlinger, Chromatin accessibility in the Drosophila embryo is determined by transcription factor pioneering and enhancer activation. Dev. Cell 58, 1898–1916.e9 (2023). doi:10.1016/j.devcel.2023.07.007.

7. I. V. Soluri, L. M. Zumerling, O. A. Payan Parra, E. G. Clark, S. A. Blythe, Zygotic pioneer factor activity of Odd-paired/Zic is necessary for late function of the Drosophila segmentation network. eLife 9, e53916 (2020).

8. T. Koromila, F. Gao, Y. Iwasaki, P. He, L. Pachter, J. P. Gergen, A. Stathopoulos, Odd-paired is a pioneer-like factor that coordinates with Zelda to control gene expression in embryos. eLife 9, e59610 (2020).

9. H.-L. Liang, C.-Y. Nien, H.-Y. Liu, M. M. Metzstein, N. Kirov, C. Rushlow, The zinc-finger protein Zelda is a key activator of the early zygotic genome in Drosophila. Nature 456, 400–403 (2008).

10. M. M. Gaskill, T. J. Gibson, E. D. Larson, M. M. Harrison, GAF is essential for zygotic genome activation and chromatin accessibility in the early Drosophila embryo. eLife 10, e66668 (2021). doi:10.7554/eLife.66668.

11. M. M. Harrison, X.-Y. Li, T. Kaplan, M. R. Botchan, M. B. Eisen, Zelda Binding in the Early Drosophila melanogaster Embryo Marks Regions Subsequently Activated at the Maternal-to-Zygotic Transition. PLoS Genet. 7, e1002266 (2011). doi:10.1371/journal.pgen.1002266.

12. X.-Y. Li, M. M. Harrison, J. E. Villalta, T. Kaplan, M. B. Eisen, Establishment of regions of genomic activity during the Drosophila maternal to zygotic transition. eLife 3, e03737 (2014). doi:10.7554/eLife.03737.

13. Y. Sun, C.-Y. Nien, K. Chen, H.-Y. Liu, J. Johnston, J. Zeitlinger, C. Rushlow, Zelda overcomes the high intrinsic nucleosome barrier at enhancers during Drosophila zygotic genome activation. Genome Res. 25, 1703–1714 (2015).

14. K. N. Schulz, E. R. Bondra, A. Moshe, J. E. Villalta, J. D. Lieb, T. Kaplan, D. J. McKay, M. M. Harrison, Zelda is differentially required for chromatin accessibility, transcription factor binding, and gene expression in the early Drosophila embryo. Genome Res. 25, 1715–1726 (2015).

15. S. L. McDaniel, T. J. Gibson, K. N. Schulz, M. F. Garcia, M. Nevil, S. U. Jain, P. W. Lewis, K. S. Zaret, M. M. Harrison, Continued Activity of the Pioneer Factor Zelda Is Required to Drive Zygotic Genome Activation. Mol. Cell 74, 185–195.e4 (2019).

16. A. B. Stergachis, B. M. Debo, E. Haugen, L. S. Churchman, J. A. Stamatoyannopoulos, Single-molecule regulatory architectures captured by chromatin fiber sequencing. Science 368, 1449–1454 (2020).

17. E. G. Swanson, Y. Mao, B. J. Mallory, M. R. Vollger, S. C. Bohaczuk, C. B. Oliveira, D. B. Lyon, J. Ranchalis, N. L. Parmalee, B. A. Cohen, J. T. Bennett, A. B. Stergachis, Mapping single-cell diploid chromatin fiber architectures using DAF-seq. Nat. Biotechnol., doi: 10.1038/s41587-025-02914-3 (2025).

18. B. R. Doughty, M. M. Hinks, J. M. Schaepe, G. K. Marinov, A. R. Thurm, C. Rios-Martinez, B. E. Parks, Y. Tan, E. Marklund, D. Dubocanin, L. Bintu, W. J. Greenleaf, Single-molecule states link transcription factor binding to gene expression. Nature 636, 745–754 (2024).

19. V. Baderna, G. Barzaghi, R. Kleinendorst, K. Chatsirisupachai, L. Moniot-Perron, C. Doyle, M. Schopp, T. Hochepied, C. Libert, D. T. Odom, J. B. Zaugg, A. R. Krebs, Cumulative transcription factor binding and p300-mediated histone acetylation drive enhancer activation frequency. Nat. Genet., doi: 10.1038/s41588-026-02703-x (2026).

20. N. J. Abdulhay, C. P. McNally, L. J. Hsieh, S. Kasinathan, A. Keith, L. S. Estes, M. Karimzadeh, J. G. Underwood, H. Goodarzi, G. J. Narlikar, V. Ramani, Massively multiplex single-molecule oligonucleosome footprinting. eLife 9, e59404 (2020).

21. T. W. Tullius, R. S. Isaac, D. Dubocanin, J. Ranchalis, L. S. Churchman, A. B. Stergachis, RNA polymerases reshape chromatin architecture and couple transcription on individual fibers. Mol. Cell 84, 3209–3222.e5 (2024). doi:10.1016/j.molcel.2024.08.013.

22. G. Hunt, R. Vaid, S. Pirogov, A. Pfab, C. Ziegenhain, R. Sandberg, J. Reimegård, M. Mannervik, Tissue-specific RNA Polymerase II promoter-proximal pause release and burst kinetics in a Drosophila embryonic patterning network. Genome Biol. 25, 2 (2024). doi:10.1186/s13059-023-03135-0.

23. F. Cardamone, A. Piva, E. Löser, B. Eichenberger, M. C. Romero-Mulero, F. Zenk, E. J. Shields, N. Cabezas-Wallscheid, R. Bonasio, G. Tiana, Y. Zhan, N. Iovino, Chromatin landscape at cis-regulatory elements orchestrates cell fate decisions in early embryogenesis. Nat. Commun. 16, 3007 (2025).

24. K. Chen, J. Johnston, W. Shao, S. Meier, C. Staber, J. Zeitlinger, A global change in RNA polymerase II pausing during the Drosophila midblastula transition. eLife 2, e00861 (2013).

25. Y. T. Ip, R. E. Park, D. Kosman, E. Bier, M. Levine, The dorsal gradient morphogen regulates stripes of rhomboid expression in the presumptive neuroectoderm of the Drosophila embryo. Genes Dev. 6, 1728–1739 (1992).

26. R. He, W. Dong, Z. Wang, C. Xie, L. Gao, W. Ma, K. Shen, D. Li, Y. Pang, F. Jian, J. Zhang, Y. Yuan, X. Wang, Z. Zhang, Y. Zheng, S. Liu, C. Luo, X. Chai, J. Ren, Z. Zhu, X. S. Xie, Genome-wide single-cell and single-molecule footprinting of transcription factors with deaminase. Proc. Natl. Acad. Sci. 121, e2423270121 (2024).

27. S. V. E. Keränen, A. Villahoz-Baleta, A. E. Bruno, M. S. Halfon, REDfly: An Integrated Knowledgebase for Insect Regulatory Genomics. Insects 13, 618 (2022). doi:10.3390/insects13070618.

28. A. Cano, M. A. Pérez-Moreno, I. Rodrigo, A. Locascio, M. J. Blanco, M. G. del Barrio, F. Portillo, M. A. Nieto, The transcription factor Snail controls epithelial–mesenchymal transitions by repressing E-cadherin expression. Nat. Cell Biol. 2, 76–83 (2000).

29. M. Leptin, twist and snail as positive and negative regulators during Drosophila mesoderm development. Genes Dev. 5, 1568–1576 (1991).

30. M. W. Perry, A. N. Boettiger, J. P. Bothma, M. Levine, Shadow Enhancers Foster Robustness of Drosophila Gastrulation. Curr. Biol. 20, 1562–1567 (2010).

31. L. Dunipace, A. Ozdemir, A. Stathopoulos, Complex interactions between cis-regulatory modules in native conformation are critical for Drosophila snail expression. Development 138, 4075–4084 (2011).

32. A. Ozdemir, L. Ma, K. P. White, A. Stathopoulos, Su(H)-Mediated Repression Positions Gene Boundaries along the Dorsal-Ventral Axis of Drosophila Embryos. Dev. Cell 31, 100–113 (2014).

33. J. P. Bothma, H. G. Garcia, S. Ng, M. W. Perry, T. Gregor, M. Levine, Enhancer additivity and non-additivity are determined by enhancer strength in the Drosophila embryo. eLife 4, e07956 (2015).

34. L. Dunipace, J. M. McGehee, J. Irizarry, A. Stathopoulos, The proximal enhancer of the snail gene mediates negative autoregulatory feedback in Drosophila melanogaster. GENETICS 230, iyaf058 (2025).

35. H. J. Doyle, R. Kraut, M. Levine, Spatial regulation of zerknüllt: a dorsal-ventral patterning gene in Drosophila. Genes Dev. 3, 1518–1533 (1989).

36. V. L. Pimmett, M. Douaihy, L. Maillard, A. Trullo, P. Garcia Idieder, M. Costes, J. Dufourt, H. Lenden-Hasse, O. Radulescu, M. Lagha, Coordinated active repression operates via transcription factor cooperativity and multiple inactive promoter states in a developing organism. Nat. Commun. 16, 8157 (2025).

37. S. S. Dima, G. T. Reeves, Bulk-level maps of pioneer factor binding dynamics during the Drosophila maternal-to-zygotic transition. Development 152, dev204460 (2025).

38. N. Maziak, Y. Zhang, F. Groll, H. E. Brown, A. Madich, Y. Kaur, M. M. Harrison, J. Zhou, J. M. Vaquerizas, Three-dimensional genome reorganization foreshadows zygotic genome activation in Drosophila. Nat. Genet. 58, 607–617 (2026).

39. A. Birnie, A. Plat, C. Korkmaz, J. P. Bothma, Precisely timed regulation of enhancer activity defines the binary expression pattern of Fushi tarazu in the Drosophila embryo. Curr. Biol. 33, 2839–2850.e7 (2023).

40. L. Wolpert, Positional information and the spatial pattern of cellular differentiation. J. Theor. Biol. 25, 1–47 (1969).

41. S. Naqvi, S. Kim, S. Tabatabaee, A. Pampari, A. Kundaje, J. K. Pritchard, J. Wysocka, Transfer learning reveals sequence determinants of the quantitative response to transcription factor dosage. Cell Genomics 5 (2025).

42. J. A. Heslop, B. Pournasr, J.-T. Liu, S. A. Duncan, GATA6 defines endoderm fate by controlling chromatin accessibility during differentiation of human-induced pluripotent stem cells. Cell Rep. 35, 109145 (2021).

43. S. Ji, F. Chen, P. Stein, J. Wang, Z. Zhou, L. Wang, Q. Zhao, Z. Lin, B. Liu, K. Xu, F. Lai, Z. Xiong, X. Hu, T. Kong, F. Kong, B. Huang, Q. Wang, Q. Xu, Q. Fan, L. Liu, C. J. Williams, R. M. Schultz, W. Xie, OBOX regulates mouse zygotic genome activation and early development. Nature 620, 1047–1053 (2023).

44. F. Lai, L. Li, X. Hu, B. Liu, Z. Zhu, L. Liu, Q. Fan, H. Tian, K. Xu, X. Lu, Q. Li, K. Feng, L. Wang, Z. Lin, H. Deng, J. Li, W. Xie, NR5A2 connects zygotic genome activation to the first lineage segregation in totipotent embryos. Cell Res. 33, 952–966 (2023).

45. W. Kobayashi, S. Ruangroengkulrith, E. N. Arslantas, A. Mohanan, K. Tachibana, Feed-forward loops by NR5A2 ensure robust gene activation during pre-implantation development. Development 153, dev205059 (2026).

46. T. W. Tullius, FiberCNN training and promoter-specific adapters, version 1.0.0. Zenodo(2026). doi:10.5281/zenodo.22062387.

47. T. W. Tullius, FiberHMM manuscript-facing source snapshot, version 2026.08.22-manuscript-snapshot. Zenodo(2026). doi:10.5281/zenodo.22062389.

48. T. W. Tullius, FiberBrowser manuscript-facing source snapshot, version 2026.08.22-manuscript-snapshot. Zenodo(2026). doi:10.5281/zenodo.22062391.

49. T. W. Tullius, Analysis code and source data for the Drosophila developmental-timer study, version 1.0.1. Zenodo(2026). doi:10.5281/zenodo.22063860.

50. I. E. Schor, J. F. Degner, D. Harnett, E. Cannavò, F. P. Casale, H. Shim, D. A. Garfield, E. Birney, M. Stephens, O. Stegle, E. E. M. Furlong, Promoter shape varies across populations and affects promoter evolution and expression noise. Nat. Genet. 49, 550–558 (2017). doi:10.1038/ng.3791.

51. N. Karaiskos, P. Wahle, J. Alles, A. Boltengagen, S. Ayoub, C. Kipar, C. Kocks, N. Rajewsky, R. P. Zinzen, The Drosophila embryo at single-cell transcriptome resolution. Science 358, 194–199 (2017). doi:10.1126/science.aan3235.

52. P. J. Batut, X. Y. Bing, Z. Sisco, J. Raimundo, M. Levo, M. S. Levine, Genome organization controls transcriptional dynamics during development. Science 375, 566–570 (2022).

53. N. Koenecke, J. Johnston, B. Gaertner, M. Natarajan, J. Zeitlinger, Genome-wide identification of Drosophila dorso-ventral enhancers by differential histone acetylation analysis. Genome Biol. 17, 196 (2016). doi:10.1186/s13059-016-1057-2.

54. A. Siepel, G. Bejerano, J. S. Pedersen, A. S. Hinrichs, M. Hou, K. Rosenbloom, H. Clawson, J. Spieth, L. W. Hillier, S. Richards, G. M. Weinstock, R. K. Wilson, R. A. Gibbs, W. J. Kent, W. Miller, D. Haussler, Evolutionarily conserved elements in vertebrate, insect, worm, and yeast genomes. Genome Res. 15, 1034–1050 (2005). doi:10.1101/gr.3715005.

55. D. Calderon, R. Blecher-Gonen, X. Huang, S. Secchia, J. Kentro, R. M. Daza, B. Martin, A. Dulja, C. Schaub, C. Trapnell, E. Larschan, K. M. O’Connor-Giles, E. E. M. Furlong, J. Shendure, The continuum of Drosophila embryonic development at single-cell resolution. Science 377, eabn5800 (2022). doi:10.1126/science.abn5800.

56. J. A. Castro-Mondragon, R. Riudavets-Puig, I. Rauluseviciute, R. Berhanu Lemma, L. Turchi, R. Blanc-Mathieu, J. Lucas, P. Boddie, A. Khan, N. Manosalva Pérez, O. Fornes, T. Y. Leung, A. Aguirre, F. Hammal, D. Schmelter, D. Baranasic, B. Ballester, A. Sandelin, B. Lenhard, K. Vandepoele, W. W. Wasserman, F. Parcy, A. Mathelier, JASPAR 2022: the 9th release of the open-access database of transcription factor binding profiles. Nucleic Acids Res. 50, D165–D173 (2022). doi:10.1093/nar/gkab1113.

57. M. Nevil, E. R. Bondra, K. N. Schulz, T. Kaplan, M. M. Harrison, Stable Binding of the Conserved Transcription Factor Grainy Head to its Target Genes Throughout Drosophila melanogaster Development. Genetics 205, 605–620 (2017). doi:10.1534/genetics.116.195685.

58. A. A. Evdokimova, A. A. Tarakanova, M. Erokhin, D. Chetverina, M. Sun, N. E. Vorobyeva, SRP/dGATAb modulates the ecdysone response bidirectionally via locus-specific regulatory landscape. Epigenetics Chromatin 19, 3 (2026). doi:10.1186/s13072-025-00652-z.

59. M. M. Kudron, A. Victorsen, L. Gevirtzman, L. W. Hillier, W. W. Fisher, D. Vafeados, M. Kirkey, A. S. Hammonds, J. Gersch, H. Ammouri, M. L. Wall, J. Moran, D. Steffen, M. Szynkarek, S. Seabrook-Sturgis, N. Jameel, M. Kadaba, J. Patton, R. Terrell, M. Corson, T. J. Durham, S. Park, S. Samanta, M. Han, J. Xu, K.-K. Yan, S. E. Celniker, K. P. White, L. Ma, M. Gerstein, V. Reinke, R. H. Waterston, The ModERN Resource: Genome-Wide Binding Profiles for Hundreds of Drosophila and Caenorhabditis elegans Transcription Factors. Genetics 208, 937–949 (2018). doi:10.1534/genetics.117.300657.

