## Supplemental text and figures for "An intrinsic chromatin timer sets the tempo for development"

**This file includes:**

Materials and Methods
Supplementary Text
Figs. S1 to S18
Tables S1 to S9
References (*6*, *10–12*, *16*, *17*, *21*, *22*, *27*, *31*, *34*, *46–59*)

**Materials and Methods**

Study design

The study tested how cis-regulatory chromatin and transcription change on the same DNA molecules during early embryogenesis and whether regulatory opening encodes developmental time before spatial activation. Ultra-deep PacBio Fiber-seq (*16*) at 2–4 and 6–8 hours provided the principal genome-wide maps; ONT Fiber-seq at 1.5–3 and 3–4.5 hours resolved the transition into gastrulation; matched 3–4.5-hour GAF-depleted and maternal *zld*-heterozygous ONT libraries tested pioneer-factor dependence; and targeted DAF-seq provided finer time courses and *dorsal*-null comparisons. Coverage and sample provenance are reported in Table S1.

Analyses were performed at the molecule level. Genome-wide temporal summaries used nonredundant CRMs integrated across stages; cross-gene summaries used genes, CRMs, or CRM pairs as specified. Each condition or time point represents one pooled population library assembled from independently collected embryo/reaction batches, and technical sequencing movies resequenced the same pool. Inference therefore uses the stated molecule-, gene-, or genomic-block-sampling unit within each pooled population rather than treating source collections or movies as biological replicates. Randomization and blinding were not applicable to pooled molecular collections; embryos were assigned by genotype and developmental collection window, and molecular processing and analysis used prespecified, condition-invariant rules.

Temporal CRM classes were defined from WT development before perturbation data were examined; matched 3–4.5-hour GAF-depletion and maternal *zld*-heterozygote deficits were then evaluated within those fixed classes (fig. S14).

Operational terminology

Nucleosome-free-region (NFR) width was treated continuously; the nested operational states were pioneered (≥175 bp), merged (≥300 bp), and actuated (≥500 bp). An opening register denotes one of two alternative NFR positions across overlapping regulatory sequence; one NFR spanning both positions was not counted as two independently open CRMs.

These state names describe physical opening depth, not inferred factor identity or function.

FiberHMM resolves footprint geometry. Assignment of a footprint or CRM to a particular transcription factor required independent sequence, binding, or perturbation evidence. The terms pioneer-associated, Zld-associated, and GAF-associated therefore refer to such independent evidence and not to footprint size alone.

Fly stocks and genetics

Wild-type controls were collected from the laboratory *yellow white* (*yw*) background. Maternal *dorsal*-null embryos were collected from homozygous $dl^1$ mothers derived from BDSC stock 3236 ($dl^1$/CyO). Maternal *zld*-heterozygous embryos were collected from mothers carrying $zld^{294}$, *P*{neoFRT}19A/FM7. GAF depletion used the maternal triple driver *P*{w[+mC]=otu-GAL4::VP16.R}1, w[*]; *P*{w[+mC]=GAL4-nanos.NGT}40; *P*{w[+mC]=GAL4::VP16-nanos.UTR}CG6325[MVD1] (MTD-GAL4, BDSC 31777): MTD-GAL4 virgin females were crossed to the UAS hairpin line y1 v1; *P*{y+t7.7 v+t1.8=TRiP.GL00699}attP2 directed against *Trl*/GAF (BDSC 41582), and embryos were collected from female progeny carrying both driver and hairpin. The two pioneer perturbations use different genetic designs and were therefore analyzed as separate within-condition contrasts rather than as dose-matched perturbations. Temporal classes were defined in wild type before either perturbation was examined.

Biological collection and reaction-pool structure

Each PacBio stage library pooled approximately 30 independently collected and Hia5-treated embryo batches; each ONT library pooled approximately six, and each DAF time point or perturbation pooled approximately ten independently collected and reacted batches. Component reactions were combined before library construction and were not separately indexed after pooling; exact component counts are therefore approximate. Collection and reaction operators and technical sequencing sources are summarized in Table S1.

Embryo collection, staging, and perturbation windows

Embryos were staged by time after egg laying. Before nuclei isolation, pooled embryos were dechorionated for 1 min 45 s in 50% bleach and rinsed. The developmental windows used for each library are reported in Table S1; the targeted DAF-seq series used adjacent half-hour or one-hour collection windows.

Whole-genome collections used nominal 1.5–3-, 2–4-, 3–4.5-, and 6–8-hour windows after egg laying. WT DAF-seq comprised adjacent half-hour bins from 1.5 to 4.5 hours; maternal *dorsal*-null DAF-seq comprised three one-hour bins. The pooled 2–4-hour WT and *dorsal*-null samples supplied the *rho* NEE comparison.

Displayed temporal-class comparisons use the matched 3–4.5-hour WT, GAF-depleted, and maternal *zld*-heterozygous ONT libraries.

PacBio Fiber-seq

Dechorionated embryos were homogenized in 250 mM sucrose, 10 mM Tris–Cl (pH 8.0), 25 mM KCl, 5 mM $MgCl_2$, and 0.1% Triton X-100 by 40 strokes with a tight Dounce pestle. Nuclei were passed through a 40-µm filter, collected at 350 × *g* for 10 min at 4°C, and resuspended in 30 µl of 15 mM Tris–Cl (pH 8.0), 15 mM NaCl, 60 mM KCl, 1 mM EDTA, 0.5 mM EGTA, and 0.5 mM spermidine. Hia5 was purified and activity-calibrated as described (*16*) and provided by the Stergachis laboratory. One unit was the amount required to fully methylate 1 µg DNA in 1 hour at 37°C under the calibration conditions. S-adenosylmethionine and Hia5 were added to 0.8 mM and 200 U, respectively; reactions proceeded for 10 min at 25°C and were quenched with SDS to 1% while mixing with a wide-bore tip. Each embryo collection was reacted separately and high-molecular-weight DNA was purified with the Promega Wizard HMW DNA Extraction Kit (A2920), and pooled 2–4- and 6–8-hour libraries were barcoded and sequenced on the PacBio Revio platform.

The analyzed 2–4- and 6–8-hour datasets each comprised one pooled stage library sequenced across 11 technical movies. FiberHMM 2.16.7 tracks contained 16,383,316 and 16,587,728 nuclear records spanning 326.74 and 332.06 Gb, respectively (2,375.5× and 2,414.1× analytical coverage). Median mapped spans were 19,908 and 19,887 bp. Complete track provenance is reported in Table S1 and the source data.

ONT Fiber-seq

ONT Fiber-seq used the embryo nuclei-isolation and Hia5-stenciling workflow above, followed by Promega Wizard HMW DNA purification and PromethION sequencing. MM/ML adenine-modification probabilities were retained for FiberHMM; perturbation BAM headers record Dorado 2.0.1 with the dna_r10.4.1_e8.2_400bps_sup@v5.2.0 base model and 6mA v1 model.

WT 1.5–3- and 3–4.5-hour tracks contained 6,067,058 and 6,444,175 nuclear records (491.7× and 462.5× coverage; median mapped spans, 11,272 and 9,155 bp). Matched 3–4.5-hour siGAF and maternal *zld*-heterozygote tracks contained 3,107,984 and 2,234,721 records (195.2× and 124.4×).

DAF-seq

Targeted DAF-seq combined amplification-compatible DddB footprinting (*17*) with the embryo nuclei-isolation workflow used here. Each 50-µL DddB footprinting reaction used 2 µL DddB in 2× DRB buffer [20 mM Tris-HCl (pH 7.5), 20 mM NaCl, and 2 mM DTT] and was incubated for 30 min at 25°C; all other processing was identical to the Fiber-seq workflow. Approximately ten independently collected embryo batches were reacted per time point or perturbation and pooled before locus-specific PCR. The nine amplicon products used in displayed figures and their dm6 intervals are listed in Table S1; all nine corresponding PCR primer pairs are listed in Table S2. ONT sequencing retained cytidine-conversion evidence in MD tags for the DddB/Nanopore FiberHMM model; platform and base-calling provenance are in Table S1.

DddB expression and purification

His-tagged DddB was coexpressed with its immunity protein in *Escherichia coli* BL21. A 1-liter LB culture inoculated at 1:100 was grown to an $OD_{600}$ of approximately 0.6, induced with 0.5 mM IPTG, and incubated with shaking for 16 hours at 18°C. Cells were collected by centrifugation at 4,000 × *g* for 30 min and resuspended in 80 ml of lysis buffer [50 mM Tris-HCl (pH 8.0), 500 mM NaCl, 10 mM imidazole, 10% glycerol, and protease-inhibitor cocktail]. Cells were lysed by five 10-s sonication pulses. Debris was removed by two successive centrifugations at 16,000 × *g* for 30 min. While the cleared lysate was stirred, 2.1 ml of neutralized 10% polyethyleneimine (PEI; Sigma, P3143) was added dropwise, and the resulting precipitate was removed by centrifugation at 16,000 × *g* for 30 min. Inclusion of the PEI precipitation step reduced carryover of *E. coli* DNA into sequencing libraries.

The His-tagged DddB–immunity-protein complex was captured on a nickel-affinity column and eluted in 50 mM Tris-HCl (pH 7.5), 300 mM NaCl, 300 mM imidazole, and 1 mM DTT. The eluate was mixed with 50 ml of denaturing buffer [8 M urea, 50 mM Tris-HCl (pH 7.5), 300 mM NaCl, and 1 mM DTT] and incubated for 16 hours at 4°C. The denatured proteins were reloaded onto the nickel-affinity column, which was washed with 50 ml of the 8 M urea buffer to remove the immunity protein. The column was then washed sequentially with denaturing buffer containing 6, 4, 2, 1, and 0 M urea. The pooled DddB eluate was dialyzed against two 1-liter changes of dialysis buffer [20 mM Tris-HCl (pH 7.5), 200 mM NaCl, 1 mM DTT, and 10% (w/v) glycerol]. Protein was concentrated at 4°C in Amicon Ultra 10-kDa centrifugal filters (Millipore) at 3,000 × *g* for 20- to 30-min intervals with intermittent mixing until the required working concentration was reached. Protein concentration was measured by Bradford assay (Bio-Rad Protein Assay), purity was assessed by Coomassie-stained SDS–polyacrylamide gel electrophoresis with BSA as a control, and aliquots were stored at −80°C.

Reference genome, gene models, and coordinate system

All analyses used the dm6 Drosophila melanogaster reference coordinate system. Gene and transcript models were taken from the standard FlyBase annotation distributed in Ensembl release 100 for the BDGP6.28 assembly (Drosophila_melanogaster.BDGP6.28.100.gtf). Coordinates were oriented by the annotated gene strand for per-gene model inputs and locus plots, while exported CRM identifiers retained genomic chromosome coordinates. Gene-linked TSS coordinates used for model inputs were refined against the strand-specific nuclear-cycle-14 CAGE data of Schor et al. (*50*; E-MTAB-4787). For each gene, annotated transcript starts were searched within ±100 bp, and the transcript and local CAGE peak with the greatest signal were used when the summed signal was at least 1; the annotated TSS was retained otherwise.

Alternative promoters were represented by separate TSS identifiers and were not pooled with primary-promoter molecules during model fitting or same-fiber tests.

Literature enhancer coordinates were obtained from REDfly (*27*) and from locus-specific primary reports. REDfly identifiers, database release, coordinate conversion if any, and the exact comparison universe are reported in the archived CRM catalog (*49*) and Table S5.

Alignment and molecule-level quality control

PacBio Hia5 reads were aligned to dm6 with pbmm2 1.8.0 using the CCS preset and coordinate sorting (pbmm2 align --preset CCS --sort). ONT Hia5 reads were aligned with minimap2 2.30-r1287 using the Nanopore preset (minimap2 -ax map-ont -y) and coordinate sorted with samtools 1.21. DddB DAF-seq reads were aligned with minimap2 2.28-r1209 or 2.30-r1287, according to the source library, using -a -x map-ont --MD (with -Y and -y where recorded), and coordinate sorted with samtools 1.19.2 or 1.21. MD tags were retained for FiberHMM to locate DddB-associated C-to-T and G-to-A conversions. Exact aligner and sorting commands are retained in BAM program headers where available.

Before quantitative molecule counting and feature-track extraction, targeted DAF-seq BAMs were deduplicated with FiberHMM's deamination-fingerprint algorithm. Coordinate duplicate marking is uninformative for these UMI-free, primer-defined amplicons because distinct molecules share alignment endpoints. Each primary read with at least 10 called conversions was represented by the set of converted reference positions detected from R/Y sequence codes, MM/ML dU calls, or MD-tag mismatches. Candidate matches were proposed by 32-component MinHash signatures in eight locality-sensitive-hashing bands and verified by exact Jaccard similarity within chromosome and alignment strand. Reads connected at Jaccard similarity of at least 0.95 were assigned to one transitive cluster, which was collapsed to the primary representative with the highest mapping quality, longest aligned span, and lowest edit distance. Reads with fewer than 10 conversions were retained without clustering. Alignment strand was used only to interpret the strand-specific conversion pattern; neither DAF-seq nor Fiber-seq read orientation was assigned biological transcriptional-strand meaning.

Unless otherwise specified, molecule-level analyses required a primary, nonduplicate, non-QC-fail, nonsupplementary alignment with mapping quality of at least 20. DAF-seq molecules additionally required valid per-read accessibility calls. ONT perturbation analyses requiring complete model input retained reads with at least five called footprints. Analyses that tested a CRM and promoter or gene body on the same molecule required continuous alignment coverage of both queried regions and the specified outer flanks. Molecules that did not span the complete threshold-specific window were treated as missing, not closed.

Technical movies were combined within each pooled PacBio stage library; condition/time-point pools and their technical source files are identified in the source-data records.

FiberHMM footprint segmentation and biological annotation

PacBio Fiber-seq, ONT Fiber-seq, and DAF-seq molecules were converted to a common footprint representation with FiberHMM 2.16.7 (tag v2.16.7; commit d4607e47e33fddc7933a5103560f34517caabe01). Assay-specific enzyme and sequencing models were selected explicitly: Hia5/PacBio for PacBio Fiber-seq, Hia5/Nanopore for ONT Fiber-seq, and DddB/Nanopore DAF mode for targeted DAF-seq. Every call set analyzed here underwent nucleosome recall and factor-sized-footprint recall. ONT Hia5 used the topology-constrained nucleosome-recall policy. The original FiberHMM transcription-state framework is described in (*21*). Assay-specific call sets were evaluated with the TSS-centered interval-geometry and external transcription-ordering checks in fig. S1B,E–H; the same fixed configuration was applied across loci, stages, and perturbations within each assay.

FiberHMM first applies an assay-specific two-state hidden Markov model to segment protected and accessible sequence. Its nucleosome-recall pass then scores the exact per-base m6A or deamination pattern against the same context-specific protected and accessible emission models. Supported accessible runs split overmerged protected intervals; a protected-evidence pass trims each surviving nucleosome to conservative boundaries bracketed by informative modification sites and records unresolved edge ambiguity. Long overmerged intervals could also be split at sample-specific nucleosome-repeat-length positions, but only when local accessibility evidence supported the candidate linker. For ONT Fiber-seq data, a topology constraint accepted only cuts that left nucleosome-sized pieces and preserved protection where sparse evidence did not resolve an edge. Intervening methyltransferase-sensitive patches (MSPs) were rederived from the refined nucleosome boundaries.

The factor-sized-footprint recall pass scanned the refined accessible intervals and short protected intervals with a local log-likelihood ratio of protected versus accessible emission. Calls required at least three informative target bases and cumulative support of at least 5 natural-log units for Hia5 or 4 units for DddB. Boundaries were placed at the nearest informative hits and misses, yielding generally smaller, cleaner intervals than the initial HMM segment. Recalled intervals of nucleosome scale were returned to the nucleosome class, overlapping short initial nucleosome candidates were removed, and the resulting nonoverlapping nucleosome/MSP tiling was regenerated. Biological labels were assigned only afterward from interval size and position: nucleosome-scale (> 89bp) intervals represented chromatin occupancy, and shorter intervals represented factor-sized protection without identifying the bound factor. At promoters and transcription units, appropriately positioned factor-sized protections were annotated operationally as preinitiation complexes, promoter-proximal paused RNA polymerase II (Pol II), or elongating Pol II.

Reconstruction of centered NFRs and opening width

For each molecule, NFRs were reconstructed directly as gaps between consecutive nucleosomes of at least 90 bp in the recall-refined FiberHMM nucleosome track. The recall output is a nonoverlapping nucleosome/MSP tiling, so the measured interval is the physical nucleosome-bounded gap rather than a separately merged call class. Factor-sized protections lying within a gap were retained but did not split the NFR. This definition measures the nucleosome-scale accessible run while preserving shorter bound complexes within it.

For a point anchor, the centered NFR was the inverse-nucleosome interval containing that anchor. For a CRM with more than one prespecified peak, the analysis either used the largest qualifying centered NFR at any prespecified anchor or used the CRM's integrated physical boundary, as specified for each analysis below. Exact continuous width required both NFR edges to be observed. Threshold analyses allowed a read-edge-censored interval only when the observed lower bound already exceeded the tested threshold and the molecule satisfied the outer-span requirement.

The primary nested thresholds were 175, 300, and 500 bp, corresponding to pioneered, merged and actuated states.

Transcription-state measurements

Transcription was read from footprint-derived Pol II states on the same molecules used to measure regulatory opening. Coordinates were oriented from the relevant TSS. Promoter initiation signal was measured in the immediate TSS region, promoter-proximal pausing in the downstream pause window, and elongating Pol II in the gene body. The model label windows were approximately −50 to +10 bp for the initiation complex, +20 to +70 bp for paused Pol II, and +100 bp through at most +1,500 bp of the gene body for elongation, with the gene-body endpoint shortened for compact genes.

Embryo analyses used sensitivity-oriented 15–70-bp Pol-II-associated protections, broader than the high-specificity settings in (*21*), because pausing is detected more readily than sparse elongation on individual molecules. These are operational transcription-associated annotations; quantitative conclusions use within-gene or matched-state contrasts, background subtraction, null comparisons, or external rank ordering rather than interpreting raw call fractions as absolute biochemical occupancy.

The principal binary productive-firing outcome for genome-wide same-fiber analyses was elongation density ≥2.857 eligible Pol-II-associated footprints per kilobase of the CRM-masked gene-body window. Productive-firing frequency denotes the fraction of eligible molecules that passed this threshold, and conditional elongating-Pol-II load denotes elongation density among productively firing molecules. High-output, gene-body eviction, promoter opening, initiation-complex occupancy, pausing without productive elongation, and productive-elongation occupancy were analyzed as distinct outcomes.

CRM analyses used transcription-state annotations in which the focal CRM was removed from the gene-body measurement region before state assignment. This prevented the CRM's own factor-sized footprints from being counted as Pol II.

<u>Global single-fiber chromatin and transcription summaries</u>

Promoter-state and gene-body summaries were computed among molecules spanning the complete promoter and transcription window (fig. S1). The mutually exclusive descriptive states were promoter-open only, initiation-complex without productive elongation, pausing without productive elongation, and closed. Elongation was allowed to co-occur with other states. Independent comparison with nuclear-cycle-14 single-cell RNA detection and matched-window PRO-seq tested population-level ordering rather than one-to-one calibration between assays.

The fixed transcription-state analysis set contained 1,301 genes selected from the 2–4-hour multigene cache. Genes entered the set when at least 500 spanning fibers were available and more than 4% carried at least two eligible elongating-Pol-II-associated calls; 20 curated developmental-patterning genes that met the depth requirement were added despite falling below the activity threshold. For the TSS-aligned geometry maps, QC-passing strand-oriented records from these 1,301 genes were sampled deterministically to 100,000 records per stage. Protected intervals of 10–90 bp and 100–300 bp were summarized separately within ±200 bp of the TSS with a shared color range within protection class. State composition was calculated within gene and averaged with equal gene weight.

External validation used the nuclear-cycle-14 single-cell RNA matrix of Karaiskos et al. (*51*) and early PRO-seq from Hunt et al. (*22*; GSM6454906), summarized over the matched strand-oriented gene-body window. Genes were ranked into scRNA-seq deciles or PRO-seq quartiles, and Fiber-seq state fractions were averaged within bins with gene-bootstrap 95% confidence intervals. The developmental-patterning analysis used 35 genes with scRNA-seq measurements and callable promoter-accessibility and transcription states in the same 2–4-hour Fiber-seq data; associations were evaluated by Spearman rank correlation. The PRO-seq concordance matrix was normalized within quartile (fig. S1).

A Pol II convoy was a maximal chain of at least two such calls in which every adjacent center-to-center gap was at most 80 bp. The 80-bp boundary was chosen from the first sustained zero crossing after the approximately 49-bp short-gap mode in the observed-minus-constrained-null spacing distribution. The null preserved gene and gene-body identity, call count and sizes, marginal positional propensity, and non-overlap. Each fiber contributed total weight one before genes were averaged. These geometric analyses quantify Pol-II-call spacing and convoy membership (fig. S2).

Gene-body nucleosome occupancy was measured across the continuous Pol II-call axis after centering within gene. To distinguish chromatin loss from simple replacement of nucleosome calls by polymerase-sized calls, each Pol II interval plus 25-bp flanks was excluded; unexcluded and 50-bp-flank analyses were included as sensitivities. Residual analyzed spans were at least 400 bp. A matched analysis compared convoy and nonconvoy fibers within gene and exact Pol II-call count. Opening-width distributions at regulatory regions were analyzed separately among molecules with bounded centered NFRs (figs. S2, S3 and S6).

High-output and chromatin-depletion summaries

High-output analyses used eligible Pol-II-sized calls from the CRM-masked 2–4-hour state table. Gene-body nucleosome occupancy was recalculated after excluding each eligible call plus 25-bp flanks and required at least 400 residual analyzed base pairs. Chromatin-depleted fibers had buffered nucleosome coverage below 50%; output was divided into fewer than four versus at least four eligible calls. The equal-gene PIC comparison required at least 10 fibers in each output group and retained 953 paired genes. The *slp1* analysis divided 2,156 fibers into 20 equal-count eviction bins and used Wilson molecule-sampling intervals. The ten-gene output-only summary used the same four-call threshold. Genome-wide distributions used fixed 8-percentage-point eviction bins from 0 to 96%; gene-bin estimates required at least five productive fibers for conditional call load and at least 10 fibers for productive-firing frequency (fig. S3).

rho NEE analysis

The *rho* neuroectodermal enhancer (NEE) was represented by two prespecified anchors at approximately −1.4 kb and −1.05 kb relative to the *rho* TSS. Each molecule was assigned the larger centered opening at either anchor. The pooled 2–4-hour DAF comparison used deduplicated recall-refined BAMs. Each retained WT or *dorsal*-null input library contributed equal weight before molecule fractions were averaged.

Opening was summarized at 175-, 300-, and 500-bp thresholds. Molecules used for the transcription comparison spanned both the NEE and the *rho* gene body. The direct transcription outcome was derived from 30–70-bp Pol II footprints in the gene body. Molecule-bootstrap intervals resampled within input libraries and recombined equal-library-weighted estimates. Opening-width survival and pausing without productive elongation measurements are shown in fig. S6A,B; the matched-background opening and elongation-positive analyses are shown in Fig. 1D,E.

The background-adjusted curve compared the NEE with exact-width non-CRM regions in the same DAF data. For each NFR-width threshold, the plotted contrast was [NEE minus matched background]WT minus [NEE minus matched background]*dorsal*-null. The primary interval was the 815-bp 2–4-hour CRM union; the two individual anchors and their outer union were included as boundary sensitivities. The curve crossed zero near 260–265 bp: *dorsal*-null molecules were relatively enriched for shorter openings, whereas WT became enriched above approximately 300 bp and showed a 7.5-percentage-point advantage at 500 bp. The same depth-resolved comparison was performed at D1, D2, C1, C2, P1, P2 and P3 at *sna* (fig. S6C,D).

Opening depth and additional shallow states

Opening-dependent transcriptional readouts in Fig. 1H were evaluated in 2–4-hour fibers using regulatory CRMs outside annotated insulators and chromatin boundaries whose widest stage-specific called boundary was at least 300 bp. CRMs first required at least 200 callable fibers; the common-support cohort then required at least 20 fibers in each of four mutually exclusive opening bands (<175, 175–299, 300–499, and ≥500 bp), yielding 1,491 CRMs. For each CRM, the productive-firing and pausing-without-productive-firing fractions in the three deeper bands were differenced from the <175-bp fraction. CRM-level differences were reduced to one median per linked gene within each model-effect class. The class score was (activating − repressive)/(activating + repressive): negative values were repressive, values from 0 to <1/3 intermediate, and values ≥1/3 activating. The three classes contained 699, 573, and 132 CRMs contributing 447, 384, and 120 genes; 87 CRMs without a resolved same-gene 2–4-hour score were excluded. Boxes show the median and interquartile range across genes, and whiskers the 5th to 95th percentiles.

Contribution of additional smaller openings at a fixed actuated CRM state

The contribution of additional small CRM-linked openings was tested among fibers spanning a fixed strand-oriented 9-kb field upstream of the selected TSS plus 500-bp support at both edges. Each annotated CRM was queried at its peak, and the same physical inter-nucleosome gap was counted once when it contained multiple peaks. Retained fibers contained exactly one CRM-linked NFR at least 500 bp wide. The identity of this actuated state was the exact set of CRM peaks contained by the NFR.

Two additional-opening definitions were evaluated. The first summed distinct 175–299-bp CRM-linked NFRs and excluded every fiber containing a CRM-linked 300–499-bp NFR. The second summed all distinct 175–499-bp CRM-linked NFRs. Opening was divided by the common 9-kb denominator. Within each gene and actuated-CRM identity, productive firing was regressed on the 50–100-versus-zero-bp/kb contrast while adjusting continuously for the width of the actuated state. Each comparison required at least ten fibers in each group, and estimates were averaged within gene before gene-bootstrap inference.

The pioneered-only contrast was unresolved at both stages, whereas the broader pioneered-or-merged dose showed a small positive association. Estimates, confidence intervals, and gene counts are reported in fig. S9A and its source data. All comparisons hold the identity and width of the observed actuated state fixed within the same 9-kb field.

Global convolutional model

The architecture factorized a shared, position-agnostic local sequence/chromatin encoder from promoter-specific regulatory adapters. This design prevented a gene/location shortcut while allowing each promoter to integrate its distinct CRM number, spacing, orientation, and distance. The resulting model is a shared local representation with gene-specific regulatory readout.

The FiberCNN transcription backbone was a stage-specific, low-receptive-field model with 15 one-base-resolution input channels. Three channels encoded definite factor-sized protection, ambiguous factor-sized protection, and nucleosome occupancy. Two learnable channels encoded NFR width from nucleosome-bounded gaps of at least 150 bp; the complementary soft transition was initialized at 500 bp with a sigmoid temperature of 50 bp and was optimized during training. Eight sequence channels encoded A/C/G/T in two chromatin contexts: beneath definite or ambiguous factor-sized protections and throughout regulatory NFRs of at least 150 bp, with overlap allowed at protected positions. Sequence outside those contexts was set to zero. The remaining channels were a valid-data mask and a nucleosome-boundary-ambiguity channel.

Neither backbone nor adapter received genomic position, relative position, distance to the TSS, or a positional embedding.

The encoder projected the 15 input channels to 128 features with a 21-bp convolution, followed by eight residual blocks. Each block contained two 21-bp dilated convolutions with batch normalization, ReLU activation, and dropout of 0.1. Dilations were 1, 2, 2, 4, 4, 4, 4, and 4, giving a 1,021-bp receptive field including the input projection while retaining one-base resolution. Features were averaged into nonoverlapping 200-bp tokens. The shared backbone heads combined masked mean and masked maximum pooling across valid tokens and used 64-unit multilayer-perceptron readouts with dropout of 0.2. The model contained 5,618,379 trainable parameters, of which 5,551,872 were in the convolutional backbone.

The label-bearing interval from TSS −200 bp through 200 bp beyond the gene end was physically excised before input, preventing direct label readout without leaving a mask edge. The 2–4-hour backbone used 1,617,028 fibers from 793 training genes and a deterministic 100,000-fiber validation subset from 236 nonoverlapping genes; complete span distributions are in the source data.

The backbone jointly learned four-class promoter state, an elongation-active gate at density 2.857, continuous elongation density, gene-body nucleosome-free fraction, and an auxiliary local occupancy target. AdamW optimization, loss weights, batch budgets, early stopping, seed, selected checkpoints, hashes, and the otherwise identical stage-specific 6–8-hour training are specified in Table S6.

<u>Per-gene adapters</u>

For each promoter and stage, the backbone was frozen and its 200-bp token representation supplied a separate adapter. Upstream input extended from TSS −9 kb to −200 bp after excision; downstream input retained as much as 6 kb beyond the gene end. Fibers spanned the complete retained window. Canonical and alternative TSSs had separate adapters but inherited the parent-gene split, yielding eight primary/alternative × upstream/downstream × stage combinations.

Each adapter used one symmetric bilinear-attention layer over 128-feature tokens, concatenated mean attended and untransformed vectors, and six 128-unit output heads. The adapter contained 222,918 trainable parameters while the 5.55-million-parameter backbone remained frozen; no position channel, distance channel, attention penalty, or auxiliary importance gate was used (Table S6).

Targets were initiation-complex occupancy, paused Pol II, promoter accessibility, gene-body nucleosome-free fraction, overall elongation, and high elongation. Fibers were assigned deterministically to 80% training, 10% validation, and 10% held-out test bins. Optimizer settings, class weighting, early stopping, and continuous-target definitions are given in Table S6.

In the primary-upstream 2–4-hour arm, 1,273 promoters had at least two evaluable outputs and a median held-out mean AUC of 0.682; *sna*, *eve*, and *ftz* scored 0.686, 0.700, and 0.785. Across all eight arms, 5,243 promoter models met this evaluability rule (Table S6).

Because transcriptional states are stochastic snapshots, AUC was interpreted as discrimination among propensities rather than deterministic prediction. Isotonic transforms fit only on validation fibers reduced mean held-out expected calibration error from 0.260 to 0.006 and Brier score from 0.209 to 0.118 without changing rank-based AUC (fig. S4).

Held-out comparisons tested each matched adapter against foreign-promoter adapters, the frozen shared backbone, and molecule-level context shuffles. Across 5,313 promoter/output comparisons, matched adapters improved AUC by 0.0320, 0.0090, and 0.1774, respectively; bootstrap intervals are shown in fig. S4 and its source data.

Counterfactual closing scans and CRM calling

Functional maps were generated by closing local observed NFRs on real fibers *in silico*, re-encoding the altered molecule, and measuring the change in each transcriptional prediction. Closing replaced the accessible interval with a locally plausible nucleosome-occupied representation while leaving the remainder of the molecule unchanged. Positive necessity values indicated that the observed opening supported an output; negative values indicated that closing the interval increased that output. These model counterfactuals were used to localize candidate functional intervals and do not by themselves establish causal sequence necessity.

Scans were performed in both upstream and downstream regulatory domains at 2–4 and 6–8 hours and for canonical and alternative TSSs. Coarse scans defined supported neighborhoods, and fine 50-bp scans localized channel-specific peaks. Fine positions below the absolute response floor were retained when supported by the coarse scan. Contiguous above-floor support was split at resolvable internal valleys so adjacent elements were not merged simply because their functional envelopes overlapped.

An initial call combined one model output with one interval of positional support. Activating and pausing- or autorepression-associated outputs were allowed to overlap. Calls were normalized across prediction output, stage, and linked TSS while preserving distinct promoter links and overlapping subelements. The cross-stage catalog contains 12,121 regulatory units across 1,276 of the 1,301 genes in the fixed transcription-state analysis set; the remaining genes had no CRM in the final cross-stage map. Of these units, 11,115 lie within 10 kb of the linked TSS. This complete CRM census supplied the starting set for downstream analyses. Requiring the widest stage-specific called boundary to be at least 300 bp yielded 6,942 resolved units. Regulatory analyses additionally excluded annotated insulators and chromatin boundaries: a CRM was ineligible when its peak overlapped a chromatin-boundary interval defined by Batut et al. (*52*), or when it mapped to a curated functional element annotated as an insulator or boundary. This exclusion identified 5,449 temporal-analysis CRMs before the boundary-width requirement; requiring a boundary of at least 300 bp yielded 3,912 for the primary early ONT temporal comparison (Table S4; archived CRM catalog and temporal-class source data (*49*)).

CRM identities preserved distinct local supports rather than chaining overlapping broad model intervals. A genomic CRM could contain more than one subelement, and a CRM could link to more than one promoter. Cross-time correspondence preserved the original stage-specific identities and accepted direct recurrences or unambiguous shifted-shape pairs; ambiguous alternatives remained separate.

Comparison with REDfly

REDfly annotations were intersected with nonredundant CRMs in dm6 coordinates (fig. S8). The CRM universe contained 6,942 elements whose largest stage-specific called boundary was at least 300 bp. Canonical REDfly annotations longer than 2 kb were considered callable when at least 70% of 200-bp interval probes lay within 100 bp of the complete gene-specific model scan. Of 1,188 callable annotations, 1,042 overlapped at least one qualifying CRM; 754 of these recovered annotations contained two or more CRMs and 288 contained one. The remaining 146 annotations were not recovered. Uncertainty across annotation-length bins was estimated by resampling nonredundant physical loci.

For the broader evidence analysis, each CRM was assigned the strongest REDfly support available. Canonical support required overlap with an *in vivo* annotation linked to the same gene; remaining evidence was assigned hierarchically as STARR-seq, other reporter, inferred, predicted, or unsupported. This analysis asks whether literature fragments contain multiple physically resolved regulatory units and quantifies the internal regulatory structure of broad reporter-tested regions.

Orthogonal CRM-validation tracks were H3K4me1 from Brennan et al. (*6*; GSE218852), cycle-14a H3K27ac and H3K4me3 from Li et al. (*12*; GSE58935), CBP/Nejire from Koenecke et al. (*53*; GSE68983), early PRO-seq from Hunt et al. (*22*; GSM6454906), and the UCSC dm6 phastCons124way track (*54*). H3K4me1, H3K27ac, CBP/Nejire, H3K4me3 and phastCons were summarized over each interval; the two PRO-seq strands were combined after strand-specific extraction. Exact-width random intervals were sampled on the same chromosome.

sna CRM architecture and overlapping-register analysis

The reported proximal deletion spans both P2 and P3, whereas the distal deletion is centered on D2 (*31*, *34*) (fig. S13A; Table S5).

For an overlapping-position comparison, each molecule was assigned to one of two mutually exclusive opening configurations. A full nucleosome-to-nucleosome NFR had to reach the tested depth and its midpoint had to lie nearer one prespecified peak; ambiguous assignments were excluded. This identifies the predominant opening position but does not prove that the embedded alternative is closed. Among 939 overlapping pairs, 659 retained separable geometry and at least 50 bp unique to each side, and 73 had at least 20 actuated fibers at each position. Seven pairs met the family-wide false-discovery-rate and effect-size criteria, including the prespecified *sna* D1/D2 and P3/P2 comparisons; nine additional pairs met the exploratory pointwise 90% confidence criterion. The 300-, 400- and 500-bp screens were analyzed as separate sensitivity families (fig. S12).

sna DAF time course and autonomous-timer analysis

The *sna* time-course anchors were fixed before trajectory analysis. P1, P2 and P3 were centered at approximately −2,500, −1,700 and −1,350 bp relative to the canonical *sna* TSS. P2 and P3 were analyzed as alternative opening positions rather than by counting one NFR as two independently open CRMs. Coordinates are listed in Table S5.

DAF denominators were reconstructed from recalled BAMs using the alignment and span criteria above. WT trajectories used the six half-hour collections from 1.5–2 through 4–4.5 hours; the lower-coverage 1–1.5-hour collection was excluded. Dorsal-null trajectories used 1.5–2.5-, 2.5–3.5- and 3.5–4.5-hour collections. WT and *dorsal*-null trajectories were computed separately. Opening rates were reported at the 175-, 300- and 500-bp thresholds (Fig. 2 and fig. S13).

Regional summaries asked whether any P1–P3 position opened; compositional summaries assigned the predominant CRM or register, with promoter opening measured independently. WT total proximal and promoter opening changed modestly while P1/P2/P3 prevalence shifted across the sampled populations.

The autonomous-timer test applied the same anchors and chronology to *dorsal*-null embryos and asked whether P1 rose and P3 declined at ≥175 bp despite loss of broad opening and productive firing. Transcription required molecules spanning the complete *sna* window; total proximal and promoter opening were analyzed separately from P1/P2/P3 composition (Fig. 2D–F and fig. S13).

Genome-wide measurement of CRM opening through time

Temporal analyses used nonredundant CRMs with resolved physical boundaries. For each CRM and sample, the largest NFR contained within the boundary, allowing at most a 50-bp overhang, was measured on every callable molecule. Opening fractions were computed at 175, 300, and 500 bp. After requiring at least 30 callable fibers at the relevant ONT and PacBio endpoints and excluding annotated insulators and chromatin boundaries, 5,449 CRMs remained. The additional requirement that the widest called CRM boundary be at least 300 bp left 3,912 CRMs in the early ONT comparison in Fig. 3A; the matched PacBio, siGAF, and *zld*-heterozygous comparisons contained 3,910, 3,909, and 3,785 CRMs, respectively (fig. S14A–C). One siGAF CRM lacked a directly observed matched endpoint and was omitted.

The WT ONT 1.5–3-to-3–4.5-hour and PacBio 2–4-to-6–8-hour comparisons were analyzed as two internally calibrated transitions. Each assay contributes a within-platform temporal contrast; the assays were joined graphically only at their overlapping developmental windows. For each within-assay transition, the expected genome-wide change was estimated on the log-odds scale after leaving out all CRMs assigned to the focal gene. The temporal residual was the observed later opening percentage minus the percentage expected from the focal CRM's initial state and the leave-one-gene-out global shift.

<u>Definition of pulse CRMs</u>

Temporal classes were defined from WT opening alone. The trajectory metric averaged the 175- and 300-bp opening fractions. Early CRMs required top-quartile early opening, an ONT temporal residual of at most −5 percentage points and PacBio loss of at least 5 points. Candidate pulse CRMs required an ONT temporal residual of at least 2 points and PacBio loss of at least 5 points; elements that remained highly open at the later PacBio stage were excluded. Late CRMs were selected independently from the stable-high class by requiring a 6–8-hour/2–4-hour opening ratio of at least 1 and a nonnegative late temporal residual. Late-rising elements were excluded from this comparator. Sequence, binding, perturbation, gene identity and transcription were not used to define any class.

These 589 CRMs are stringent landmarks within a continuous regulatory landscape (Figs. 3 and 4; figs. S14 and S15; archived temporal-class source data (*49*)). Nearby gate choices preserved the same class cores (fig. S14I). In that panel, early-residual and ONT-rise rows vary the 1.5–3-to-3–4.5-hour stage-adjusted change gate, and PacBio-loss rows vary the required 2–4-to-6–8-hour decrease in percentage points. Endpoint rows vary the quantile of 6–8-hour opening used to identify persistently high CRMs, retention rows vary the 6–8-hour/2–4-hour opening ratio, and late-residual rows vary the stage-adjusted 2–4-to-6–8-hour change required for late membership. Coverage and molecule-half sensitivities are provided in the figure source data.

The descriptive embedding in Fig. 3D was fixed independently of these stringent classes. It combines measured embryonic opening states and transitions with paired sequence-predicted accessibility at 2–4 and 6–8 hours. Temporal classes, motifs, gene identity, transcription, and perturbation responses were excluded from the embedding fit. After the common CRM filters, broad WT-defined regimes contained 103 early-declining, 279 transient gastrulation-opening, and 596 late CRMs. These broad regimes describe structure within the continuum; the stringent classes above provide the fixed cohorts for class-wise inference.

<u>Independent sci-ATAC validation</u>

The 589 temporal CRMs were compared with the whole-embryo sci-ATAC-seq atlas of Calderon et al. (*55*; GEO GSE190149). Each CRM was assigned an overlapping external peak, choosing the peak whose center was closest to the CRM peak when more than one overlapped; 574 CRMs mapped to one of the 110,185 external peaks. External early accessibility was the 2–4-hour counts-per-million value, and external late accessibility was the mean of the 4–8- and 6–10-hour values. Late-to-early change was log2[(late CPM + 1)/(early CPM + 1)]. Class distributions used class-pure unique external peaks so that one broad peak was counted in only one temporal class. Cross-assay concordance was reduced to one median per linked gene and evaluated by Spearman rank correlation. A partial rank correlation additionally controlled both assays' initial states.

Zld and GAF motif and binding annotations

Motifs were scanned over the integrated physical CRM form in dm6 sequence using matched Trl/GAF and vfl/Zld position-weight matrices at MOODS $P \leq 10{-}3$. Reverse-complement and near-identical hits within 10 bp were collapsed. Motif frequency, best-site score above the matched calling threshold, longest exact GAGA platform, and inter-platform spacing were analyzed as continuous quantities. Best-site strength was summarized among CRMs with at least one called motif, whereas motif frequency supplied the unconditional occurrence measure. Motif profiles were taken from JASPAR 2022 (*56*).

Matched stage-5 GAF and Zld ChIP-NEXUS tracks from Brennan et al. (*6*; GSE218852) were summarized over each CRM, adjusted by equal-width flanks 1 kb from each edge, and percentile-ranked within factor. Binding balance was GAF minus Zld percentile. Sequence and binding annotations were added only after temporal classification; class contrasts are reported in Fig. 3C and fig. S14G. References 10 and 11 supplied occupancy support for the motif-site screen in fig. S14E.

Motif positioning relative to single-fiber NFR width was tested only within the 6,942-CRM parent census whose widest stage-specific boundary was at least 300 bp. Excluding 335 CRMs annotated as insulators or chromatin boundaries left 6,607 regulatory CRMs before motif- and depth-specific requirements. For each observed NFR, motif overlap was compared with eight same-width windows placed at random within the same CRM, preserving CRM width, motif burden, and the empirical NFR-width distribution. Observed-to-shuffled ratios were averaged equally across genes and confidence intervals were obtained by resampling genes. The exact-boundary GAF, Zld, and pooled Dl/Twi/Bcd analysis included 3,209 motif-containing CRMs. A second analysis restricted motifs to sites supported by published factor-occupancy data and included 1,495 qualifying CRMs. Narrow-NFR preference was defined as the observed-to-shuffled enrichment in NFRs <300 bp divided by that in NFRs ≥300 bp. Robust calls required both enrichment within narrow NFRs and narrow-versus-wide specificity at minimum-depth requirements of 5, 10, and 20 NFRs per gene. The 300-bp split in this analysis describes molecule-level NFR width and is distinct from the ≥300-bp total CRM-boundary eligibility rule (fig. S14D,E). Occupancy support used GAF GSE152770 (*10*), Zld GSE30757 (*11*), Grh GSE83305/GSM2199199 (*57*), and S2 Srp GSE304853 (*58*). Broad embryonic ENCODE support (*59*) used GATAe/grn ENCSR625TAA and ENCSR909QHH, Stat92E ENCSR290OJD, Pnt ENCSR997UIM, Hr78 ENCSR280XEB, and Mad ENCSR875FVD. These datasets supplied occupancy-supported motif sites across the relevant developmental interval.

Pioneer-perturbation response calls

The primary perturbation outcome in Fig. 3E was the empirical 3–4.5-hour opening deficit at each CRM: WT minus perturbation percentage of fibers with a contained NFR of at least 175 bp. Each comparison required at least 30 callable molecules in WT and the corresponding perturbation, used exact CRM boundaries with at most 50 bp of overhang, and retained only regulatory CRMs outside annotated insulators and boundaries. Positive values indicate less opening after perturbation.

After the common ≥300-bp filter, siGAF measurements were available for 104 early CRMs, 243 pulse CRMs and 239 late CRMs; maternal *zld*-heterozygous measurements were available for 103, 233 and 232, respectively. Equal-CRM means and chromosome-stratified, gene-unioned

1-Mb block-bootstrap intervals were calculated from the directly observed 3–4.5-h opening deficits (Table S8).

Timed CRM identity and transcriptional output

Temporal classes were compared within the same gene and stage in the WT PacBio datasets (Fig. 4A,B and fig. S16). For each CRM, focal fibers carried a boundary-anchored NFR at or above the tested opening threshold; the primary threshold was 500 bp. Productive firing and pausing without productive elongation were measured from the enhancer-masked gene labels. Each CRM effect was centered on the linked gene's overall state frequency. Pairwise comparisons used distinct temporal CRMs linked to the same promoter and reported the later-class-minus-earlier-class percentage-point difference. After the common ≥300-bp filter, the pair families contained 64 pulse-minus-early comparisons at 2–4 hours, 23 late-minus-early comparisons at 6–8 hours and 72 late-minus-pulse comparisons at 6–8 hours. A neutral interval of ±2.5 percentage points was used only to summarize direction; continuous effects were used for inference. Threshold sensitivities crossed 300-, 500- and 700-bp focal openings with productive-elongation-density cutoffs of 2, 2.857 and 4.5.

The fine DAF time courses measured full-gene productive firing, pausing without productive elongation, the probability that any timed CRM was open at least 175 bp, promoter opening, and temporal-program composition. CRM-union rates were measured molecule by molecule. The *ftz* program grouped distal UPS and Zebra as early, proximal UPS and R as pulse, and the neurogenic elements as late. The separate PacBio endpoint supplied late support outside the DAF-seq time series; the position-resolved DAF series is shown in fig. S18A. Additional DAF-seq analyses at *Kr* and *tll* used the same ≥175-bp opening and full-gene productive-firing definitions; the *Kr* collections were plotted in chronological source order (fig. S18B,C). Independent 2–4-hour PacBio CRM-state comparisons at *Dtg*, *gsb*, and *otk2* used the same 500-bp focal-opening and enhancer-masked transcription definitions, while the corresponding ONT panels measured CRM opening and gene-spanning transcription-associated fibers at 1.5–3 and 3–4.5 hours (fig. S17).

Genome-wide test of additivity among independently opened CRMs

Additivity was tested independently of the CNN's activating or repressive role labels. For each canonical or alternative TSS, all pairs of stage-specific CRMs with actuated-state boundaries were enumerated. Both CRMs were required to have a called boundary of at least 300 bp and to lie outside annotated insulators and chromatin boundaries. There was no CNN-role or hard-distance filter. The effective distance limit arose from joint molecule span and state coverage; canonical pairs extended to approximately 14.8 kb.

For each pair, molecules were required to cover both physical boundaries without read-edge censoring. The queried boundaries were derived from stage-specific NFRs that reached at least 500 bp. A molecule matched such a boundary when its NFR missed no more than 150 bp of the boundary, overlapped at least 300 bp of it, and was at least 350 bp at the primary tolerance. Thus, the pair states are boundary-matched opening configurations, not literal ≥500-bp bins on every molecule. Each molecule was assigned to state 00, 10, 01, or 11. A single NFR matching both elements was labeled shared and excluded from state 11, because it does not demonstrate two independently open CRMs. Each element required at least 10 open-state molecules. Formal productive-firing inference additionally required at least 50 molecules in state 00 and at least 10 in each of states 10, 01, and 11. Pairs overlapping a protein-coding promoter were summarized descriptively but excluded from family-wide inference restricted to distal regulatory pairs. Duplicate pair definitions yielding identical molecule-state partitions were tested once.

The primary interaction was defined on the productive-firing-probability scale as

Interaction = $P$(productive firing | 11) − $P$(productive firing | 10) − $P$(productive firing | 01) + $P$(productive firing | 00).

An interaction of zero is the additive expectation for the two individual increments above the 00 baseline. Positive and negative one-sided tests were performed from an HC1-robust linear probability model. The sensitivity model adjusted for the molecule's global accessibility fraction, its square, and the number of other independently open CRMs on that fiber. CNN role annotations were merged only after molecular measurement and never defined the tested pair universe. Benjamini–Hochberg correction was applied once per stage across the combined family of distal canonical- and alternative-promoter pairs, with $q \leq 0.10$ used for the discovery screen.

The analysis included 664 adequately sampled pairs at 2–4 hours and 882 at 6–8 hours. Interaction estimates were centered near zero; pointwise 90% intervals nominated subadditive, superadditive, and override-like candidates, while stage-wide Benjamini–Hochberg correction defined the stricter discovery subset. Counts, distributions, and named examples are reported in fig. S9 and Table S7.

Conditional elongating-Pol-II load was tested separately among productively firing molecules using the same covariates and minimum state counts of 20 in state 00 and five in states 10, 01, and 11. This secondary conditional estimate was reported separately because conditioning on productive firing can introduce selection effects.

Genome-wide and locus-resolved CRM co-opening

Genome-wide co-opening was measured separately in the WT 2–4-h and 6–8-h PacBio libraries. The analysis used nonredundant CRMs integrated across stages whose widest called boundary was at least 300 bp. CRMs overlapping annotated insulators or chromatin boundaries were excluded, as were physically overlapping CRM pairs because one nucleosome-free region could satisfy both elements by geometry. Pairs linked to the same analyzed TSS were included without a hard distance cutoff. A molecule was jointly callable only when it covered both complete CRM boundaries. For each CRM, the largest internal nucleosome-free region contained within its boundary, allowing at most 50 bp of overhang at either edge, was scored at the 175-bp pioneered, 300-bp merged and 500-bp actuated thresholds.

Each pair and threshold required at least 300 jointly callable molecules and 10 in each marginal open and closed state. The 00/10/01/11 table was tested by two-sided Fisher exact test and summarized by the Haldane-corrected log2 odds ratio; identical state partitions were counted once and Benjamini–Hochberg correction was applied within stage and threshold. Matched-scale comparisons used pairs measurable at all three thresholds, resampled genes for intervals, and used paired Wilcoxon tests on gene medians. Distance bins were 0–2, 2–4, 4–6, 6–8, and 8–12 kb.

To compare physical co-opening with transcriptional additivity, the same adequately sampled four-state CRM pairs used in fig. S9 were analyzed. A single nucleosome-free region spanning both CRM boundaries remained in the shared state. The physical statistic was the Haldane-corrected log2 odds ratio calculated from the 00, 10, 01, and 11 counts; the transcriptional statistic was the HC1-adjusted productive-firing interaction defined above. Spearman rank correlations were calculated separately at 2–4 and 6–8 hours, describing association within the adequately sampled pair family.

Position-resolved DAF-seq maps were constructed at *eve*, *ftz*, *sna* and *Kr*. The low-depth 1–1.5-h collection was omitted, and adjacent half-hour WT collections were pooled as 1.5–2.5,

2.5–3.5 and 3.5–4.5 h. Positions were sampled every 25 bp. On each molecule, a position was scored as pioneered when it lay within an internal nucleosome-free region of at least 175 bp; positions outside the aligned span were uncallable. Every matrix pixel is the pairwise-complete Pearson correlation between the two binary position states and required at least 150 molecules spanning both positions. Raw correlations were used for measurement; a one-bin, 25-bp Gaussian smoothing was applied only for display. Colored boxes mark CRM boundaries, with nonoverlapping arms used to distinguish the overlapping *Kr* CD2 elements.

Temporal-program analysis used five complete half-hour WT bins from 1.5–2 through 3.5–4 hours. A program was open when any member CRM was ≥175 bp and actuated when any member was ≥500 bp on the same jointly callable molecule. *eve* compared one early with one pulse CRM; *ftz* grouped distal UPS/Zebra versus proximal UPS/R; and *Kr* compared CD1 with CD2 proximal. Within dual-open molecules, coordinated expansion was tested against conditional independence by Jeffreys–Dirichlet intervals and stratified odds ratios; peak-versus-off-peak conversion used Fisher tests. A sensitivity removed fibers on which one NFR spanned both programs, and Benjamini–Hochberg correction was applied across loci.

<u>zen ventral repression element analysis</u>

The *zen* ventral repression element (VRE) analysis used the prespecified CRM at chr3R:6,755,493–6,755,818, centered at chr3R:6,755,650 (−1,457 bp from the *zen* TSS). The canonical REDfly interval RFRC:0000000543.006 spans chr3R:6,755,181–6,755,805. These coordinates were used as the physical CRM and literature annotation, respectively; neither was inferred from the DAF phenotype.

The CRM model independently identified the VRE as a strong active-output necessity peak at 2–4 hours, with a coincident but smaller autorepression-associated component. The focused empirical analysis then asked whether opening width at the VRE was associated with zen transcription on the same molecule and how Dorsal altered the opening spectrum.

The quantitative DAF genotype comparison was restricted to the first matched developmental window: WT 1.5–2- and 2–2.5-hour libraries and the *dorsal*-null 1.5–2.5-hour library. Position-resolved context maps additionally pooled the available collections within each genotype into 1.5–2.5-, 2.5–3.5-, and 3.5–4.5-hour windows (fig. S7A,B); later and aggregate libraries were excluded from the opening-spectrum and transcription comparisons. An opening was the gap between consecutive at-least-90-bp nucleosomes in the recall-refined FiberHMM track that contained the prespecified CRM anchor. Callable molecules spanned the anchor by 500 bp on each side and carried at least five footprint intervals. NFR-width bins were <175, 175–299, 300–499, and at least 500 bp. The opening-spectrum analysis contained 4,765 exact-width WT and 4,366 exact-width *dorsal*-null molecules; the genotype transcription comparison contained 4,900 WT and 4,380 *dorsal*-null gene-callable molecules.

The primary genotype outcome was the unconditional frequency of each opening state rather than conversion conditional on a smaller opening; opening and productive-firing estimates are reported in fig. S7C,D and its source data.

Opening-distribution standardization accounted for a substantial but incomplete component of the genotype difference in *zen* productive firing, supporting access-mediated restriction without claiming that access is sufficient.

An independent 2–4-hour PacBio analysis used 2,212 callable VRE-to-*zen* molecules. Exact-bounded opening width was associated with productive-firing entry, whereas conditional elongating-Pol-II load among firing molecules was not resolved. Together, the data indicate that Dorsal normally limits access to a productive VRE configuration (fig. S7C,D).

Dorsal motifs were called with the JASPAR2022 dl matrix at score at least 300. Fifteen overlapping raw hits were collapsed into five genomic clusters centered approximately −267, −198, −148, +41, and +126 bp from the prespecified CRM peak. The motif clusters support the known Dorsal VRE architecture but were not used to define the opening or transcription outcomes.

Genome-browser visualization

Representative genome-browser views and single-fiber snapshots were rendered from FiberHMM BAM and interval annotations with FiberBrowser 2.15.4. Exported vector graphics preserved the displayed genomic coordinates and annotation layers; panel assembly did not alter molecule or interval positions.

Statistical analysis

Unless otherwise stated, effect sizes are reported in absolute percentage points. Binomial intervals were Wilson intervals or molecule-bootstrap intervals as specified. Continuous molecule-level associations used Spearman rank correlation. Source- or movie-direction analyses first reduced each source to one estimate and then used signed-rank or exact sign tests. Gene-level paired effects used gene bootstrap intervals and Wilcoxon signed-rank tests. Categorical enrichment used Fisher exact tests with the stated one- or two-sided alternative. Multiple-testing correction used the Benjamini–Hochberg procedure within the complete prespecified family for a stage and outcome.

Molecule-bootstrap intervals resampled molecules within source and combined source estimates with equal weight where specified. Because each condition/time point is one pooled population library, intervals quantify molecule-, gene-, or genomic-block sampling within that population rather than between-library biological-replicate variation.

All class definitions, thresholds, source exclusions, and analysis roles were recorded in machine-readable manifests. Randomized analyses used fixed seeds recorded in those manifests. CRM catalogs, temporal and perturbation results, same-fiber pair measurements, and the panel-level figure-source index are supplied in the archived analysis release (*49*). Public software versions, release identifiers, repository commits, and archived output checksums are summarized in Table S9 and will accompany the code release.

Data and code availability

Processed PacBio, ONT, and DAF-seq data have been deposited in the Gene Expression Omnibus, and raw sequencing data in the Sequence Read Archive; accession numbers will be provided upon publication. Processed FiberHMM tracks, the CRM catalog, source-data tables, and figure-source manifests are supplied in the archived analysis release (*49*). Analysis outputs are checksum indexed in the figure-source manifests and Table S9.

Machine-readable provenance links each analysis to inputs, software, parameters, and checksums. Manuscript releases of FiberCNN (*46*), FiberHMM (*47*), FiberBrowser (*48*), and the analysis and figure-generation code and source data (*49*) are archived in Zenodo; live repositories are linked from the archive records.

### Supplementary Text

Validation of single-fiber transcription states

FiberHMM resolves protection geometry; promoter complexes and polymerases are annotated afterward from protection size, strand-oriented position and genomic context. Genes detected in more nuclear-cycle-14 nuclei and genes with greater early PRO-seq signal contain progressively more transcription-associated Fiber-seq molecules (fig. S1). This concordant population ordering supports the state definitions, while the sparse molecule-level states show why transcription is better modeled as a probability than as a deterministic outcome.

Elongating-Pol-II-sized protections form closely spaced convoys with an observed spacing peak near 49 bp (fig. S2). Gene-body nucleosome occupancy falls continuously as the number of Pol II protections increases, even after the bases occupied by Pol II and their flanks are excluded (fig. S3). Convoy membership does not explain additional nucleosome loss after matching the exact Pol II count. Convoys therefore describe the local organization of elongating polymerases, whereas gene-body chromatin loss primarily follows total transcriptional load.

Model validation and recovery of known regulatory structure

The model separates a shared local sequence–chromatin representation from promoter-specific regulatory integration because enhancer number and spacing vary substantially among genes. Matched promoter adapters outperform adapters trained for other promoters and molecule-shuffled controls on held-out fibers. Isotonic calibration also brings predicted probabilities into close agreement with observed held-out state frequencies (fig. S4). Representative maps at *ftz*, *eve*, *gt* and *slp1* show that the same procedure localizes both productive- and nonproductive-state associations (fig. S5).

The CRM map recovers known regulatory regions while resolving additional internal structure. Among 1,188 callable canonical REDfly annotations longer than 2 kb, 1,042 overlap at least one qualifying CRM and 754 of those recovered annotations contain at least two qualifying CRMs (fig. S8B,C). Subdivision increases with annotation length (fig. S8D). Thus, a broad validated reporter fragment often contains several physically resolved regulatory units.

Functional separation among model-predicted CRM classes is modest at 175–299 bp and widens as openings reach 300–499 and ≥500 bp: activating calls show the largest gains in productive firing and the greatest loss of pausing without productive firing, whereas repressive calls show smaller productive gains and retain more pausing (Fig. 1H). The ordered separation supports the model-defined classes. Additional pioneered-only openings (175–299 bp) do not measurably increase productive firing once the identity and width of the single actuated state are held constant. The broader pioneered-or-merged range (175–499 bp) has a small positive association (fig. S9A).

Distinct regulatory functions within and between CRMs

Alternative opening positions within overlapping sequence can have different transcriptional associations. Seven overlapping CRM pairs pass the family-wide comparison at the 500-bp threshold, including the *sna* P2/P3 and D1/D2 pairs (fig. S12). The proximal endogenous deletion contains both P2 and P3 and therefore does not isolate P2, whereas the distal deletion is centered on D2 (fig. S13A,B).

Distinct CRMs usually retain their individual contributions when open together. Across 664 2–4-hour and 882 6–8-hour adequately sampled pairs, productive-firing interactions were centered near zero. At directional FDR ≤0.10, 1/8 and 0/9 pairs were superadditive/subadditive at 2–4 and 6–8 hours, respectively. The broader pointwise-90%-interval tier contained 30/66 and 38/94 superadditive/subadditive pairs. The subadditive tier included 34 and 62 override-like configurations; 4 and 5 passed the nested directional-FDR override definition (fig. S9; Table S7).

CRM pairs co-opened above marginal expectation, with stronger coupling in the actuated than pioneered or merged states and positive association through 8–12 kb (fig. S10). DAF-seq resolved time-dependent off-diagonal co-opening blocks at *eve*, *ftz*, *sna*, and *Kr*. At *eve* and *ftz*, coordinated conversion from dual opening to dual actuation peaked around 2.5–3 hours and remained associated after conditioning on dual opening and excluding fibers spanned by one cross-program NFR (fig. S11). Thus, the gastrulation peak reflects both increased co-opening and coordinated expansion of opened programs.

Pair-specific co-opening was largely separable from transcriptional synergy: productive-firing interactions were centered near zero and only weakly related to co-opening strength (figs. S9 and S10). Together, the analyses support a model in which coupling changes the probability of reaching particular joint CRM configurations, while already open CRMs contribute predominantly additively. This effective opening-rate control is estimated from the observed population-state frequencies.

The *zen* VRE illustrates a different mechanism of repression. VRE opening is positively associated with entry into productive firing, yet Dorsal limits VRE opening in WT embryos. Loss of Dorsal increases VRE access and releases *zen* productive firing (fig. S7C,D). An element can therefore have positive transcriptional potential when open but function genetically as a spatial repressor because access to that productive state is restricted.

Additional consequences of timed CRM exchange

The stringent temporal classes mark extremes of a continuous regulatory landscape; the independent embedding in Fig. 3D provides a descriptive view of that continuum.

Temporal classes were associated with heterogeneous transcriptional changes: within-gene comparisons and additional *Kr*, *tll*, *Dtg*, *gsb*, and *otk2* time courses included decreases, little change, and increases as early and pulse CRMs exchanged prevalence (figs. S16 to S18).

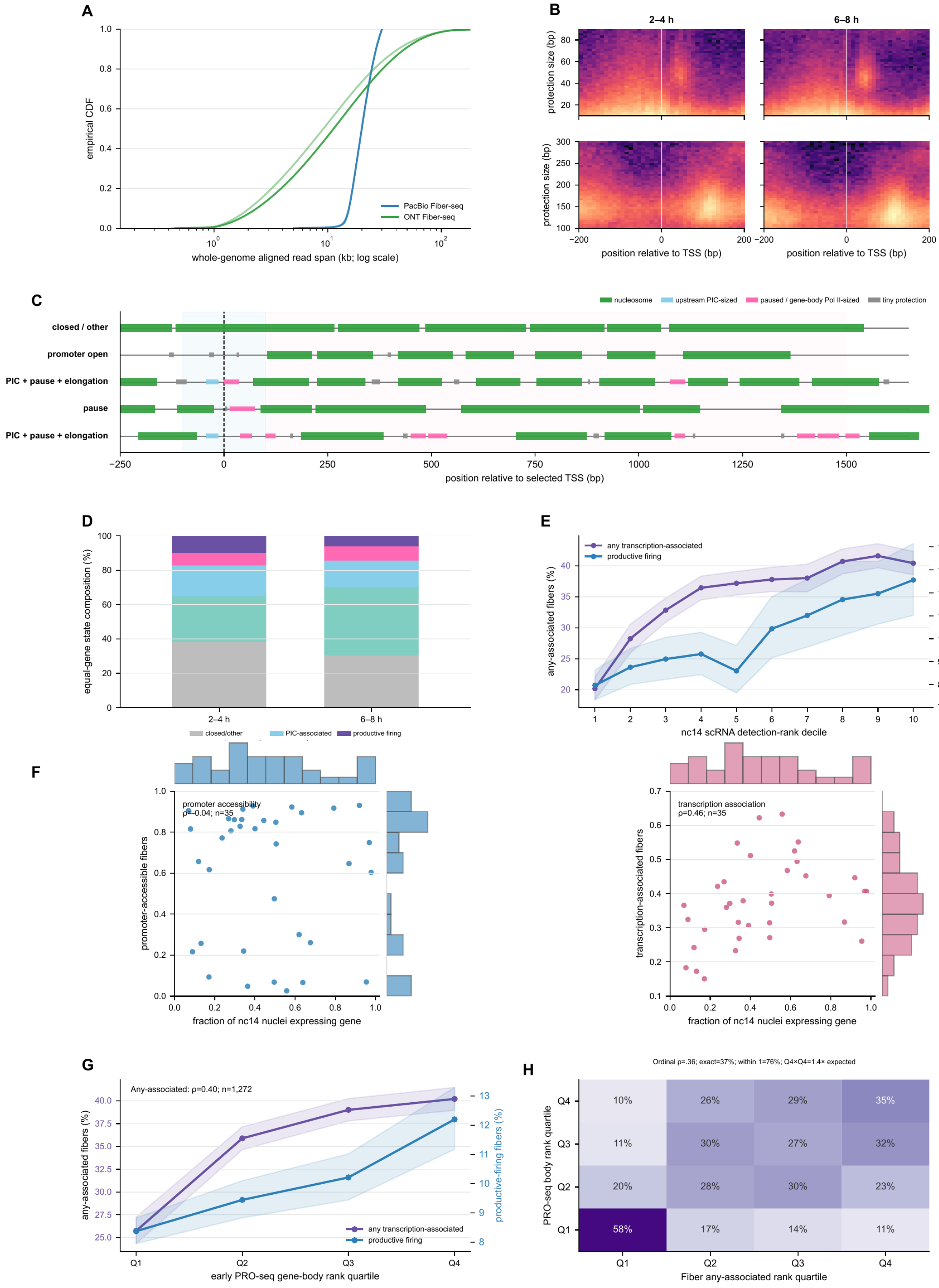


| | Q1 | Q2 | Q3 | Q4 |
|---|---|---|---|---|
| Q4 | 10% | 26% | 29% | 35% |
| Q3 | 11% | 30% | 27% | 32% |
| Q2 | 20% | 28% | 30% | 23% |
| Q1 | 58% | 17% | 14% | 11% |

**Fig. S1. Assay geometry and population ordering validate the transcription-associated Fiber-seq states.**

(**A**) Aligned-span distributions for the two whole-genome WT assays; provenance, coverage, and DAF amplicons are in Table S1. (**B**) TSS-aligned 10–90-bp protections and 100–300-bp nucleosome protections at 2–4 and 6–8 hours from deterministic samples of 100,000 QC-passing records per stage across all 1,301 genes with usable records in the full transcription-state analysis set. Maps span ±200 bp and share a color range within protection class. (**C**) Representative fibers from the operational state hierarchy in D. Green, nucleosomes; blue, upstream PIC-sized protection; pink, paused or gene-body Pol-II-sized protection; gray, tiny protection. Row labels enumerate all visible transcription-associated states and are therefore nonexclusive. (**D**) Equal-gene mutually exclusive state composition. (**E**) Mean any-associated and productive-firing fractions across nc14 scRNA-seq detection-rank deciles from Karaiskos et al. (*51*), with separate y axes and gene-bootstrap 95% intervals. (**F**) Matched comparisons for 35 developmental-patterning genes. The 2–4-hour Fiber-seq promoter-accessibility fraction from the same molecules is not ordered by the fraction of nc14 nuclei expressing the gene (Spearman $\rho = -0.04$), whereas the fraction with any transcription-associated footprint is positively ordered ($\rho = 0.46$). Marginal histograms show the distributions. (**G**) Corresponding Fiber-seq fractions across early PRO-seq gene-body quartiles from Hunt et al. (*22*; GSM6454906; n = 1,272 genes with finite PRO-seq body ranks). (**H**) Row-normalized concordance between PRO-seq and Fiber-seq any-associated quartiles. External RNA assays are population averages; Fiber-seq states are sparse single-molecule annotations.

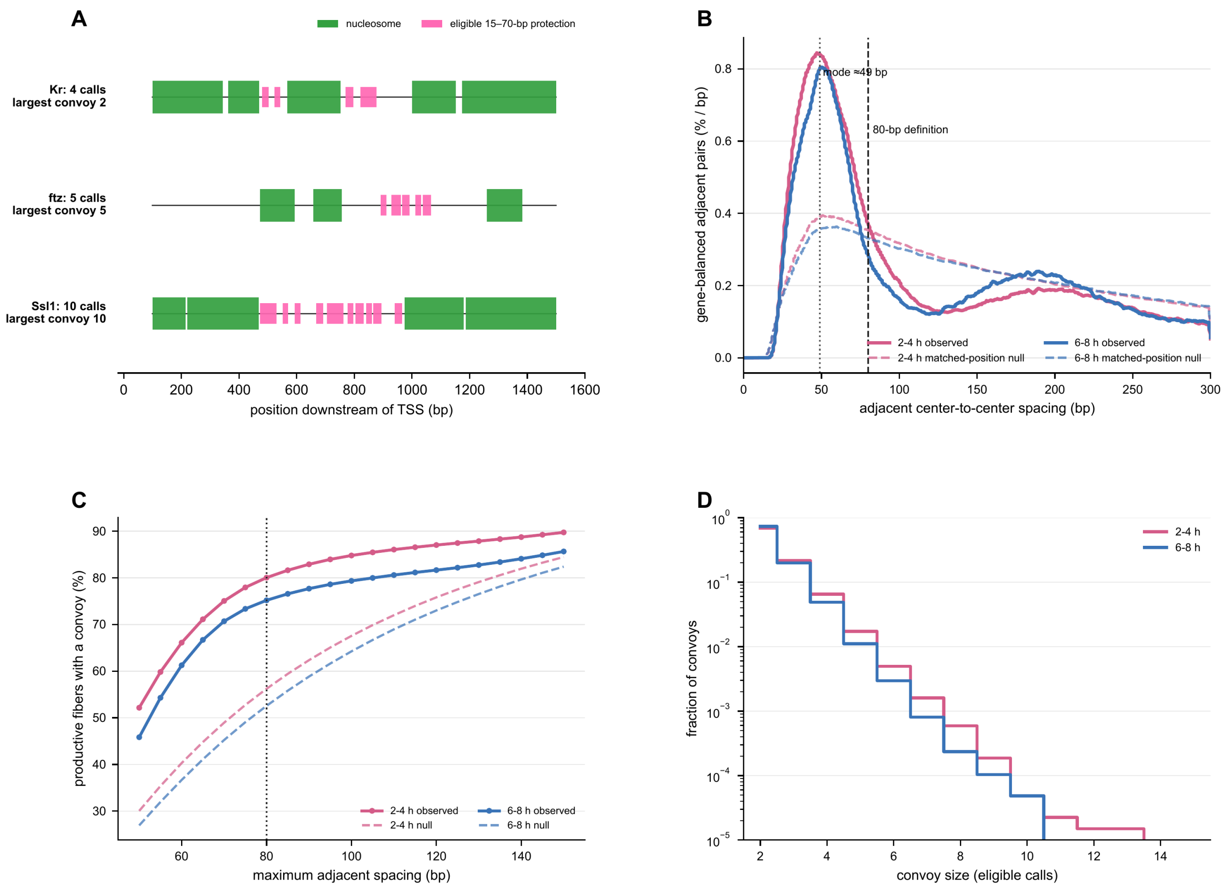
A
nucleosome
eligible 15–70-bp protection
Kr: 4 calls
largest convoy 2
ftz: 5 calls
largest convoy 5
Ssl1: 10 calls
largest convoy 10
position downstream of TSS (bp)
B
mode ≈49 bp
80-bp definition
gene-balanced adjacent pairs (% / bp)
2-4 h observed
2-4 h matched-position null
6-8 h observed
6-8 h matched-position null
adjacent center-to-center spacing (bp)
C
productive fibers with a convoy (%)
2-4 h observed
2-4 h null
6-8 h observed
6-8 h null
maximum adjacent spacing (bp)
D
fraction of convoys
2-4 h
6-8 h
convoy size (eligible calls)

**Fig. S2. Elongating-Pol-II-sized protections form nonrandom local convoys.**

(**A**) Exact 2–4-hour fibers containing maximum convoys of two, five, and ten eligible calls. Green, nucleosomes; pink, all operationally eligible 15–70-bp protections, including the shortest calls. (**B**) Gene-balanced observed and constrained-position-null distributions of adjacent center-to-center spacing at both stages. The observed mode is approximately 49 bp; the dashed vertical line marks the 80-bp operational definition. (**C**) Fraction of productive fibers containing at least one convoy across nearby maximum-spacing definitions. (**D**) Fraction of convoys by convoy size on a logarithmic scale. A convoy contains at least two eligible 15–70-bp gene-body protections connected by no more than 80 bp center spacing. The null preserves gene, call count, and call sizes while preventing overlap; the analysis reports spacing and convoy size.

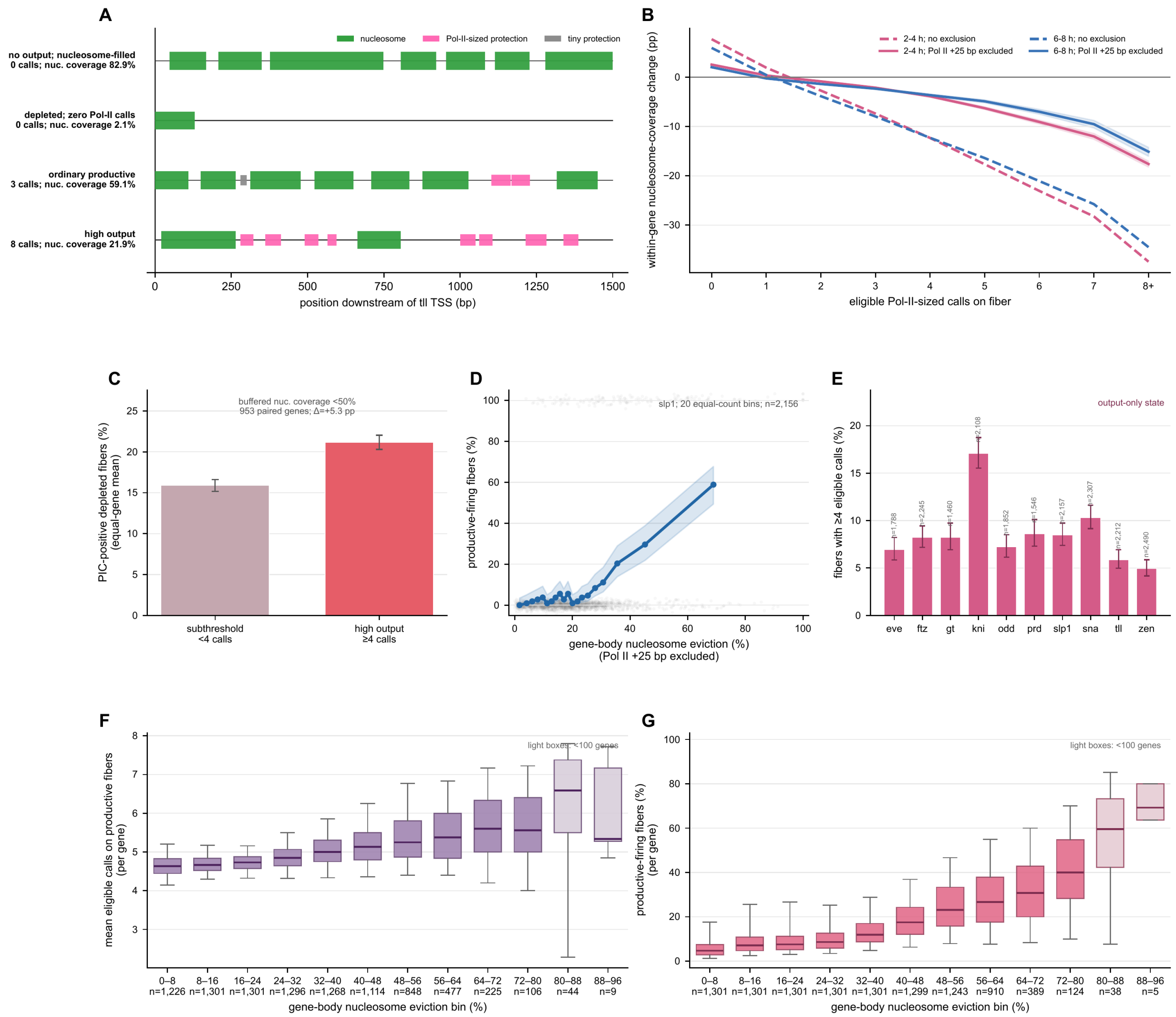

A
nucleosome
Pol-II-sized protection
tiny protection
no output; nucleosome-filled
0 calls; nuc. coverage 82.9%
depleted; zero Pol-II calls
0 calls; nuc. coverage 2.1%
ordinary productive
3 calls; nuc. coverage 59.1%
high output
8 calls; nuc. coverage 21.9%
position downstream of tll TSS (bp)
B
2-4 h; no exclusion
2-4 h; Pol II +25 bp excluded
6-8 h; no exclusion
6-8 h; Pol II +25 bp excluded
within-gene nucleosome-coverage change (pp)
eligible Pol-II-sized calls on fiber
C
buffered nuc. coverage <50%
953 paired genes; Δ=+5.3 pp
PIC-positive depleted fibers (%)
(equal-gene mean)
subthreshold
<4 calls
high output
≥4 calls
D
slp1; 20 equal-count bins; n=2,156
productive-firing fibers (%)
gene-body nucleosome eviction (%)
(Pol II +25 bp excluded)
E
output-only state
fibers with ≥4 eligible calls (%)
eve ftz gt kni odd prd slp1 sna tll zen
F
light boxes: <100 genes
mean eligible calls on productive fibers
(per gene)
0–8 8–16 16–24 24–32 32–40 40–48 48–56 56–64 64–72 72–80 80–88 88–96
n=1,226 n=1,301 n=1,301 n=1,296 n=1,268 n=1,114 n=848 n=477 n=225 n=106 n=44 n=9
gene-body nucleosome eviction bin (%)
G
light boxes: <100 genes
productive-firing fibers (%)
(per gene)
0–8 8–16 16–24 24–32 32–40 40–48 48–56 56–64 64–72 72–80 80–88 88–96
n=1,301 n=1,301 n=1,301 n=1,301 n=1,301 n=1,299 n=1,243 n=910 n=389 n=124 n=38 n=5
gene-body nucleosome eviction bin (%)

**Fig. S3. Gene-body nucleosome depletion tracks output but can persist without ongoing Pol-II footprints.**

(**A**) Representative 2–4-hour *tll* fibers spanning a nucleosome-filled no-output state, a depleted zero-call state, ordinary productive output, and high output. Green, nucleosomes; pink, eligible Pol-II-sized protections; gray, tiny protections. The vertical axis marks the TSS. (**B**) Equal-gene within-gene nucleosome-coverage change across the continuous eligible-call axis before interval exclusion (dashed) and after excluding each eligible Pol-II interval plus 25-bp flanks (solid); bands are gene-bootstrap 95% confidence intervals. (**C**) Equal-gene PIC-call frequency among chromatin-depleted fibers with either subthreshold output or high output. Both states require buffered nucleosome coverage below 50%; the 953 paired genes have at least 10 fibers in each state. Error bars are gene-bootstrap 95% confidence intervals. (**D**) 2–4-hour *slp1* productive-firing frequency across 20 equal-count bins of gene-body nucleosome eviction. The shaded band is a 95% Wilson molecule-sampling interval; faint points show individual binary fiber outcomes. (**E**) Fraction of 2–4-hour fibers with at least four eligible Pol-II-sized calls across the ten-gene set. This output-only state is defined independently of chromatin depletion; error bars are 95% Wilson molecule-sampling intervals. (**F**) Distribution across genes of mean eligible-call load conditional on productive firing in fixed 8%-wide nucleosome-eviction bins. Gene-bin cells require at least five productive fibers. (**G**) Distribution across genes of productive-firing frequency in the same fixed eviction bins. Gene-bin cells require at least 10 fibers. In F and G, boxes show the median and interquartile range, whiskers span the 5th–95th percentiles, and light boxes denote bins supported by fewer than 100 genes. Chromatin measurements in B–D, F, and G exclude eligible Pol-II intervals plus 25-bp flanks and require at least 400 residual base pairs.

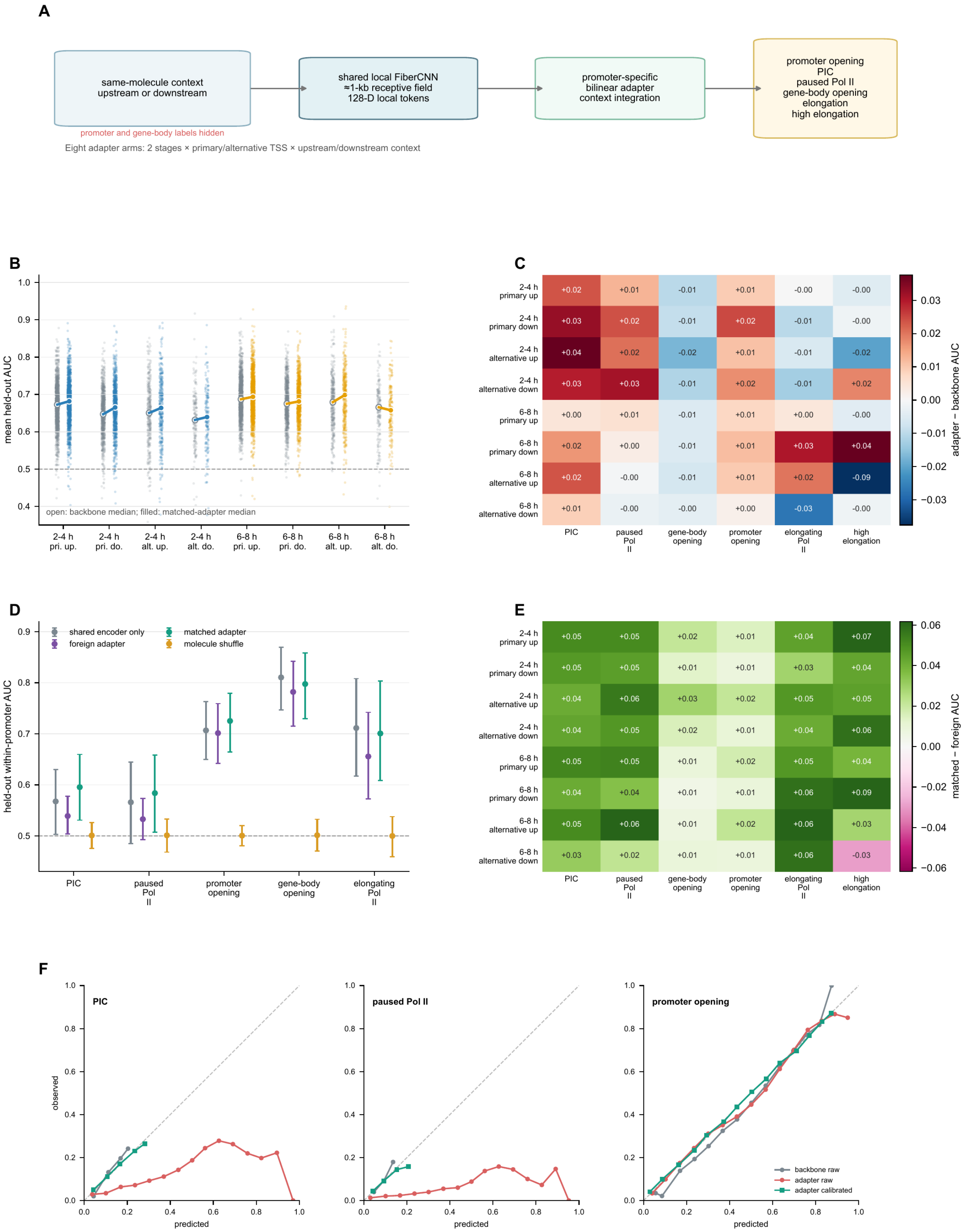
A
same-molecule context
upstream or downstream
promoter and gene-body labels hidden
shared local FiberCNN
≈1-kb receptive field
128-D local tokens
promoter-specific
bilinear adapter
context integration
promoter opening
PIC
paused Pol II
gene-body opening
elongation
high elongation
Eight adapter arms: 2 stages × primary/alternative TSS × upstream/downstream context
B
mean held-out AUC
open: backbone median; filled: matched-adapter median
2-4 h pri. up.
2-4 h pri. do.
2-4 h alt. up.
2-4 h alt. do.
6-8 h pri. up.
6-8 h pri. do.
6-8 h alt. up.
6-8 h alt. do.
C
PIC
paused Pol II
gene-body opening
promoter opening
elongating Pol II
high elongation
2-4 h primary up: +0.02 +0.01 -0.01 +0.01 -0.00 -0.00
2-4 h primary down: +0.03 +0.02 -0.01 +0.02 -0.01 -0.00
2-4 h alternative up: +0.04 +0.02 -0.02 +0.01 -0.01 -0.02
2-4 h alternative down: +0.03 +0.03 -0.01 +0.02 -0.01 +0.02
6-8 h primary up: +0.00 +0.01 -0.01 +0.01 +0.00 -0.00
6-8 h primary down: +0.02 +0.00 -0.01 +0.01 +0.03 +0.04
6-8 h alternative up: +0.02 -0.00 -0.01 +0.01 +0.02 -0.09
6-8 h alternative down: +0.01 -0.00 -0.00 +0.00 -0.03 -0.00
adapter − backbone AUC
D
shared encoder only
foreign adapter
matched adapter
molecule shuffle
held-out within-promoter AUC
PIC
paused Pol II
promoter opening
gene-body opening
elongating Pol II
E
PIC
paused Pol II
gene-body opening
promoter opening
elongating Pol II
high elongation
2-4 h primary up: +0.05 +0.05 +0.02 +0.01 +0.04 +0.07
2-4 h primary down: +0.05 +0.05 +0.01 +0.01 +0.03 +0.04
2-4 h alternative up: +0.04 +0.06 +0.03 +0.02 +0.05 +0.05
2-4 h alternative down: +0.04 +0.05 +0.02 +0.01 +0.04 +0.06
6-8 h primary up: +0.05 +0.05 +0.01 +0.02 +0.05 +0.04
6-8 h primary down: +0.04 +0.04 +0.01 +0.01 +0.06 +0.09
6-8 h alternative up: +0.05 +0.06 +0.01 +0.02 +0.06 +0.03
6-8 h alternative down: +0.03 +0.02 +0.01 +0.01 +0.06 -0.03
matched − foreign AUC
F
PIC
paused Pol II
promoter opening
observed
predicted
backbone raw
adapter raw
adapter calibrated

**Fig. S4. FiberCNN architecture, held-out performance and calibration.**

(**A**) A shared convolutional encoder learns local sequence and chromatin features after promoter and gene-body labels have been removed from the input. Separate promoter-specific adapters integrate those local features for each developmental stage, promoter class, and upstream or downstream regulatory interval. Neither component receives genomic position or distance to the TSS. (**B**) Mean held-out AUC across evaluable outputs for each gene. Linked open and filled medians compare the shared encoder alone with the matched promoter-specific adapter across the eight combinations of stage, promoter class, and direction. (**C**) Median change in AUC produced by the matched adapter for each output. (**D**) Held-out within-promoter AUC for the shared encoder alone, adapters trained for other promoters, the matched adapter, and molecule-shuffled controls. Points and intervals show medians and interquartile ranges across evaluable promoters. (**E**) Median matched-minus-foreign-adapter AUC for each output. (**F**) Reliability curves for initiation-complex occupancy, paused Pol II, and promoter opening in the representative 2–4-hour primary-upstream analysis before and after isotonic calibration. The diagonal denotes perfect calibration. All values are from held-out fibers.

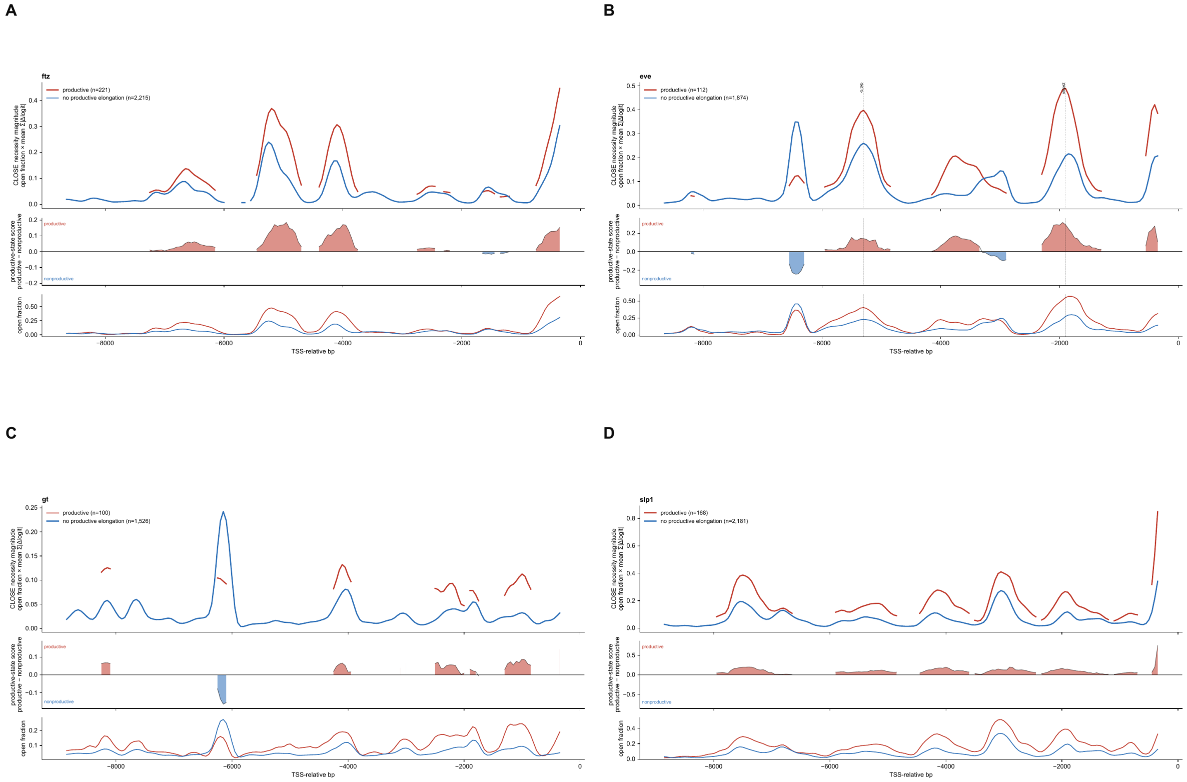
A
B
C
D
ftz
productive (n=221)
no productive elongation (n=2,215)
eve
productive (n=112)
no productive elongation (n=1,874)
gt
productive (n=100)
no productive elongation (n=1,526)
slp1
productive (n=168)
no productive elongation (n=2,181)
CLOSE necessity magnitude
open fraction × mean |Δlogit|
productive-state score
productive − nonproductive
productive
nonproductive
open fraction
TSS-relative bp

**Fig. S5. Representative promoter-specific regulatory maps.**

(A to D) Computational closing maps for *ftz*, *eve*, *gt*, and *slp1*. For each position, the open interval was replaced *in silico* by a locally plausible closed configuration and the change in predicted transcriptional state was measured. Upper traces show the response on productively firing fibers and on pooled fibers without productive firing. Lower traces show the weighted difference between those responses; positive values indicate preferential support for the productive state, whereas negative values indicate preferential support for a nonproductive state. The maps localize model-predicted regulatory requirements.

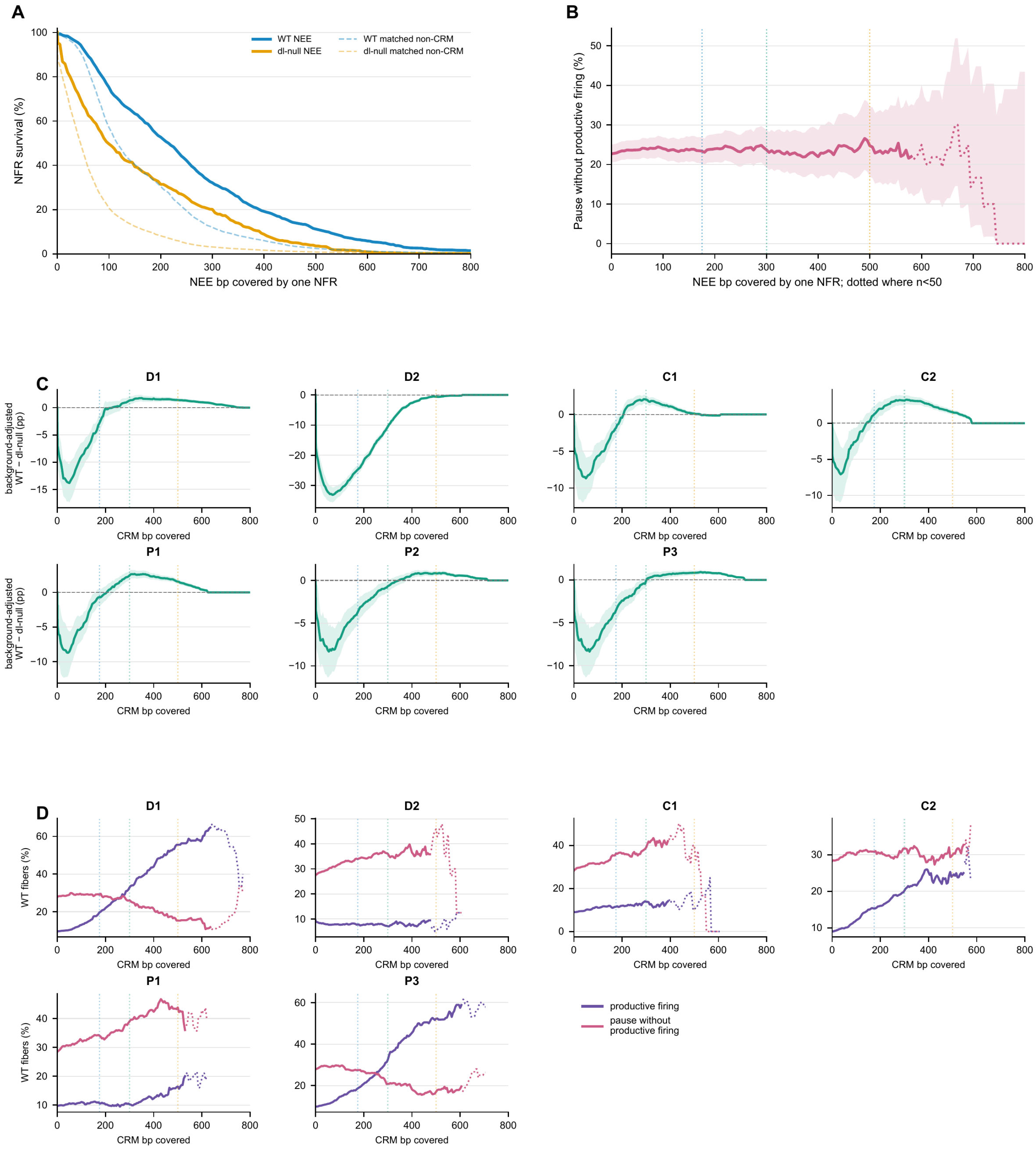
A
NFR survival (%)
NEE bp covered by one NFR
WT NEE
dl-null NEE
WT matched non-CRM
dl-null matched non-CRM
B
Pause without productive firing (%)
NEE bp covered by one NFR; dotted where n<50
C
D1
D2
C1
C2
P1
P2
P3
background-adjusted
WT − dl-null (pp)
CRM bp covered
D
WT fibers (%)
productive firing
pause without
productive firing

**Fig. S6. Pioneered, merged and actuated states at individual loci.**

(**A**) WT and dorsal-null DAF-seq opening-width survival curves at the complete rho NEE and at matched non-CRM intervals. (**B**) Pausing without productive elongation in independent WT 2–4-hour Fiber-seq molecules, grouped by the number of rho NEE bases covered by one NFR. Dotted portions contain fewer than 50 fibers. (**C**) Background-adjusted opening curves from the complete 1.5–4.5-hour WT and dorsal-null DAF-seq series for *sna* D1, D2, C1, C2, P1, P2, and P3. D1/D2 and P2/P3 overlap physically and therefore yield correlated measurements. (**D**) WT productive-firing and pausing-without-productive-elongation frequencies as a function of opening width at D1, D2, C1, C2, P1, and P3. P2 is omitted because its overlap with P3 prevents an isolated output estimate. The 175-, 300-, and 500-bp thresholds describe nested opening depths; transcriptional coupling is evaluated independently. Intervals quantify molecule sampling and, where applicable, sampling across matched background intervals from pooled collections.

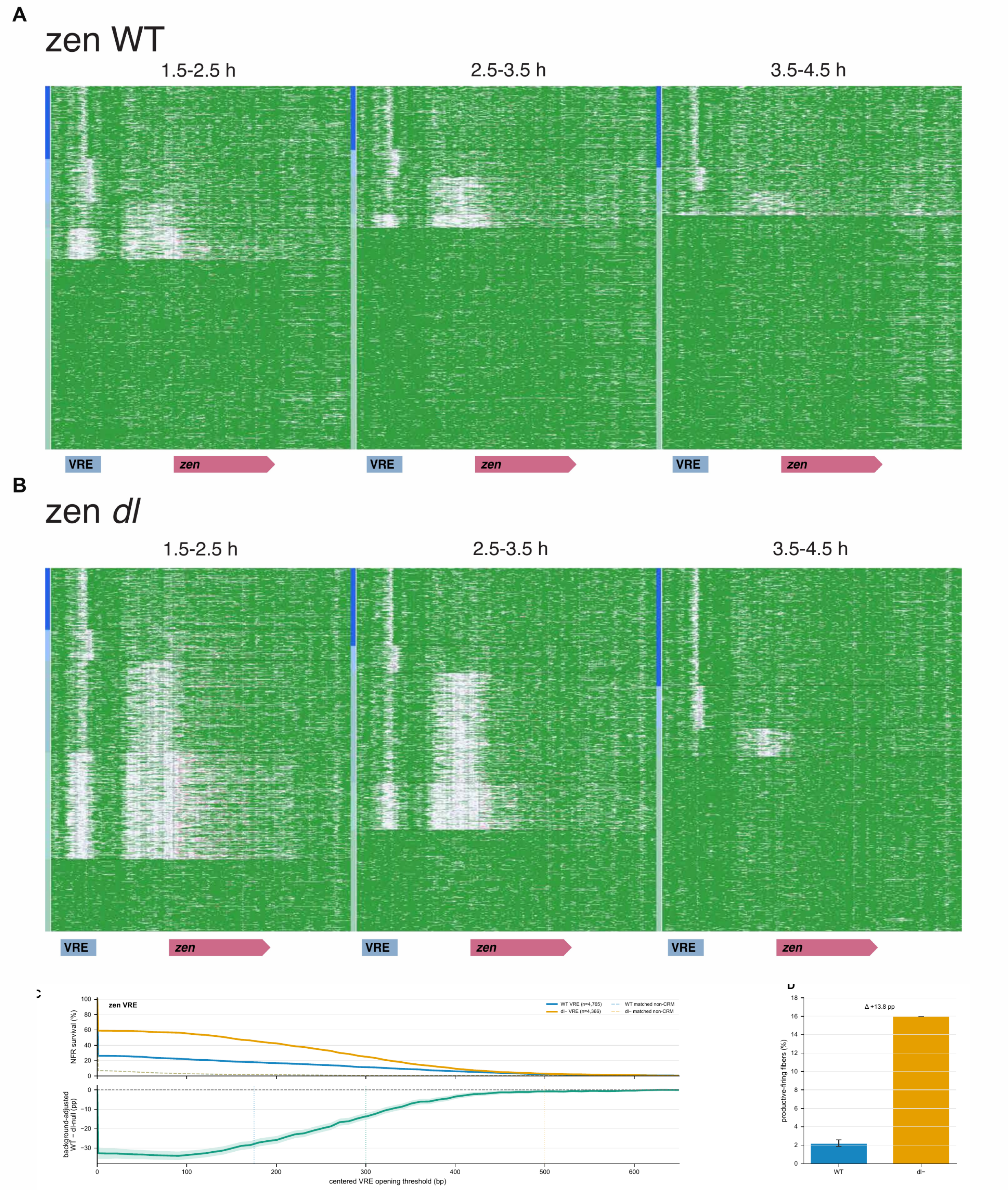
A
zen WT
1.5-2.5 h
2.5-3.5 h
3.5-4.5 h
VRE
zen
VRE
zen
VRE
zen
B
zen dl
1.5-2.5 h
2.5-3.5 h
3.5-4.5 h
VRE
zen
VRE
zen
VRE
zen
zen VRE
WT VRE (n=4,765)
dl− VRE (n=4,366)
WT matched non-CRM
dl− matched non-CRM
NFR survival (%)
background-adjusted
WT − dl-null (pp)
centered VRE opening threshold (bp)
Δ +13.8 pp
productive-firing fibers (%)
WT
dl−

**Fig. S7. Dorsal restricts a productive opening state at the zen VRE.**

(A and B) Position-resolved WT and maternal *dorsal*-null DAF-seq fibers across pooled 1.5–2.5-, 2.5–3.5-, and 3.5–4.5-hour windows at the *zen* locus. Green intervals are nucleosome-sized protections, white intervals are NFRs, the blue interval marks the VRE, and the magenta arrow marks *zen*. (**C**) Opening-width survival at the exact VRE boundary in the first matched window (WT, n = 4,765; *dorsal*-null, n = 4,366) and at exact-width matched non-CRM intervals. The lower trace shows the matched-background-adjusted WT-minus-*dorsal*-null difference across nested opening thresholds. (**D**) Productive-firing frequency across *zen* in the same first developmental window (WT, n = 4,900; *dorsal*-null, n = 4,380); the bars differ by 13.8 percentage points. Productive firing requires at least 2.857 eligible Pol-II-associated footprints per kilobase of the internally CRM-masked gene-body window. Intervals quantify molecule and matched-background sampling within pooled collections.

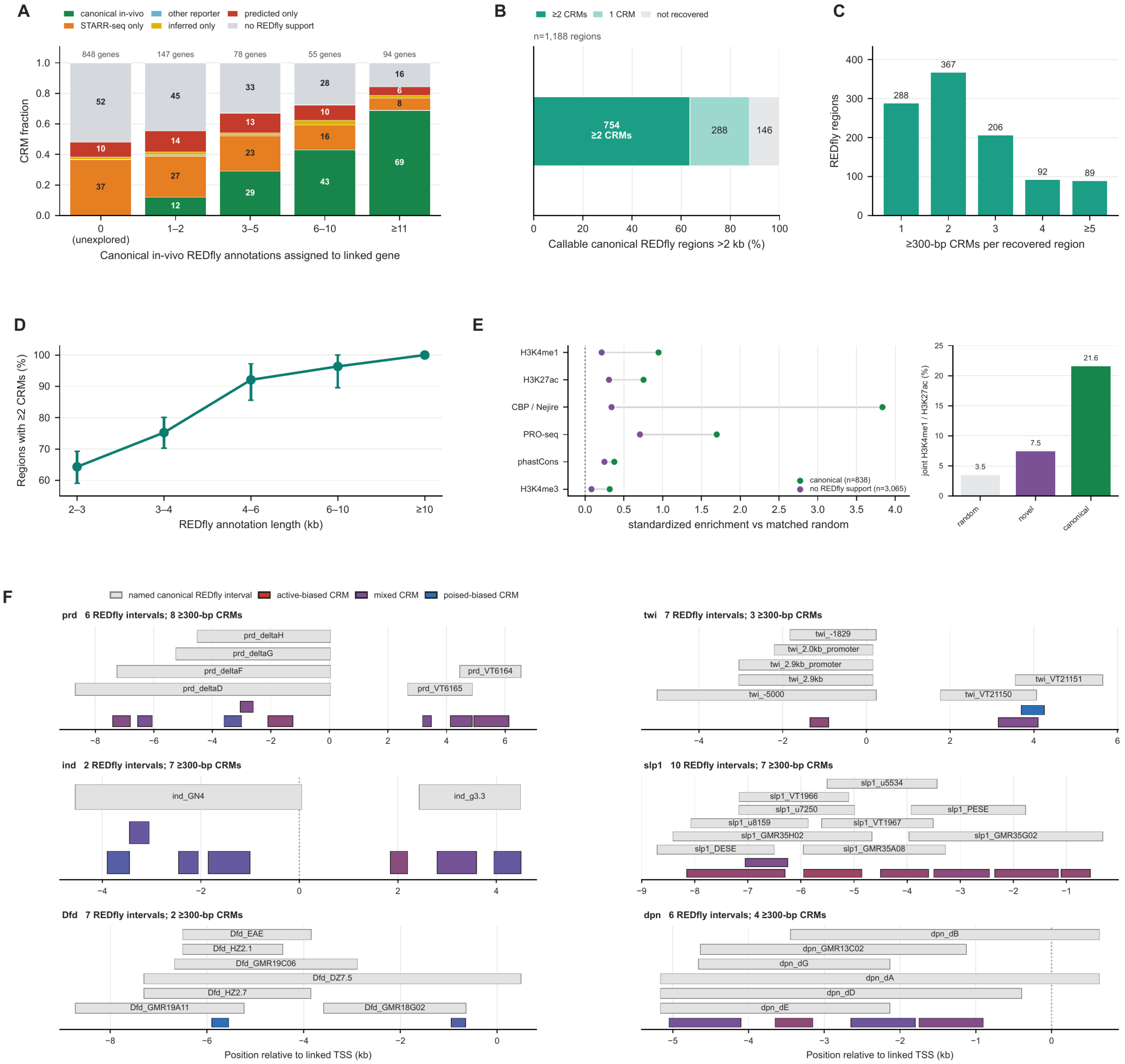
A
canonical in-vivo
other reporter
predicted only
STARR-seq only
inferred only
no REDfly support
848 genes
147 genes
78 genes
55 genes
94 genes
CRM fraction
0
(unexplored)
1–2
3–5
6–10
≥11
Canonical in-vivo REDfly annotations assigned to linked gene
B
≥2 CRMs
1 CRM
not recovered
n=1,188 regions
754
≥2 CRMs
288
146
Callable canonical REDfly regions >2 kb (%)
C
REDfly regions
288
367
206
92
89
≥300-bp CRMs per recovered region
D
Regions with ≥2 CRMs (%)
REDfly annotation length (kb)
2–3
3–4
4–6
6–10
≥10
E
H3K4me1
H3K27ac
CBP / Nejire
PRO-seq
phastCons
H3K4me3
canonical (n=838)
no REDfly support (n=3,065)
standardized enrichment vs matched random
joint H3K4me1 / H3K27ac (%)
3.5
7.5
21.6
random
novel
canonical
F
named canonical REDfly interval
active-biased CRM
mixed CRM
poised-biased CRM
prd 6 REDfly intervals; 8 ≥300-bp CRMs
prd_deltaH
prd_deltaG
prd_deltaF
prd_deltaD
prd_VT6164
prd_VT6165
twi 7 REDfly intervals; 3 ≥300-bp CRMs
twi_-1829
twi_2.0kb_promoter
twi_2.9kb_promoter
twi_2.9kb
twi_-5000
twi_VT21151
twi_VT21150
ind 2 REDfly intervals; 7 ≥300-bp CRMs
ind_GN4
ind_g3.3
slp1 10 REDfly intervals; 7 ≥300-bp CRMs
slp1_u5534
slp1_VT1966
slp1_u7250
slp1_PESE
slp1_u8159
slp1_VT1967
slp1_GMR35H02
slp1_GMR35G02
slp1_DESE
slp1_GMR35A08
Dfd 7 REDfly intervals; 2 ≥300-bp CRMs
Dfd_EAE
Dfd_HZ2.1
Dfd_GMR19C06
Dfd_DZ7.5
Dfd_HZ2.7
Dfd_GMR19A11
Dfd_GMR18G02
dpn 6 REDfly intervals; 4 ≥300-bp CRMs
dpn_dB
dpn_GMR13C02
dpn_dG
dpn_dA
dpn_dD
dpn_dE
Position relative to linked TSS (kb)
Position relative to linked TSS (kb)

**Fig. S8. Broad REDfly enhancers commonly contain multiple physically resolved CRMs.** All panels use the 6,942 nonredundant CRMs whose largest stage-specific boundary is at least 300 bp. (**A**) Best REDfly support for each CRM, stratified by the number of canonical *in vivo* REDfly annotations previously assigned to the linked gene. Canonical support requires overlap with an *in vivo* annotation assigned to the same gene; remaining support is classified hierarchically as STARR-seq, other reporter evidence, inferred, predicted, or unsupported. Numbers above bars give genes. (**B**) Recovery of 1,188 callable canonical REDfly annotations longer than 2 kb. Of these, 1,042 overlap at least one qualifying CRM: 754 contain at least two CRMs and 288 contain one; 146 are not recovered. (**C**) Number of ≥300-bp CRMs within each of the 1,042 recovered annotations. (**D**) Fraction of recovered annotations containing at least two CRMs across annotation-length bins. Intervals were obtained by bootstrapping nonredundant physical loci. (**E**) Orthogonal biochemical and evolutionary validation of the regulatory CRM census. Points compare the 838 CRMs with canonical *in vivo* REDfly support and the 3,065 CRMs without REDfly support with exact-width, same-chromosome random intervals. Values are mean-signal differences standardized by the random-window standard deviation for H3K4me1 (Brennan et al. (*6*); GSE218852), H3K27ac and H3K4me3 (Li et al. (*12*); GSE58935), CBP/Nejire (Koenecke et al. (*53*); GSE68983), PRO-seq (Hunt et al. (*22*); GSM6454906), and dm6 phastCons124way. The adjacent bars give the fraction exceeding the random-window 90th percentile for both H3K4me1 and H3K27ac: 3.5% of random intervals, 7.5% of REDfly-unsupported CRMs, and 21.6% of canonical CRMs. Pioneer-factor tracks were excluded from the orthogonal evidence panel. This panel begins with the 6,607 regulatory CRMs after exclusion of annotated insulators and chromatin boundaries. (**F**) REDfly annotations and called CRMs at *prd*, *twi*, *ind*, *slp1*, *Dfd*, and *dpn*. Gray bars show named canonical REDfly intervals; colored rectangles show CRMs with a peak inside the displayed span. Each CRM is shown once using its widest called boundary across stages. Color denotes the balance between model support for productive and nonproductive transcriptional states. Dashed lines mark linked TSSs. The overlaps reveal substructure within reporter-tested regions.

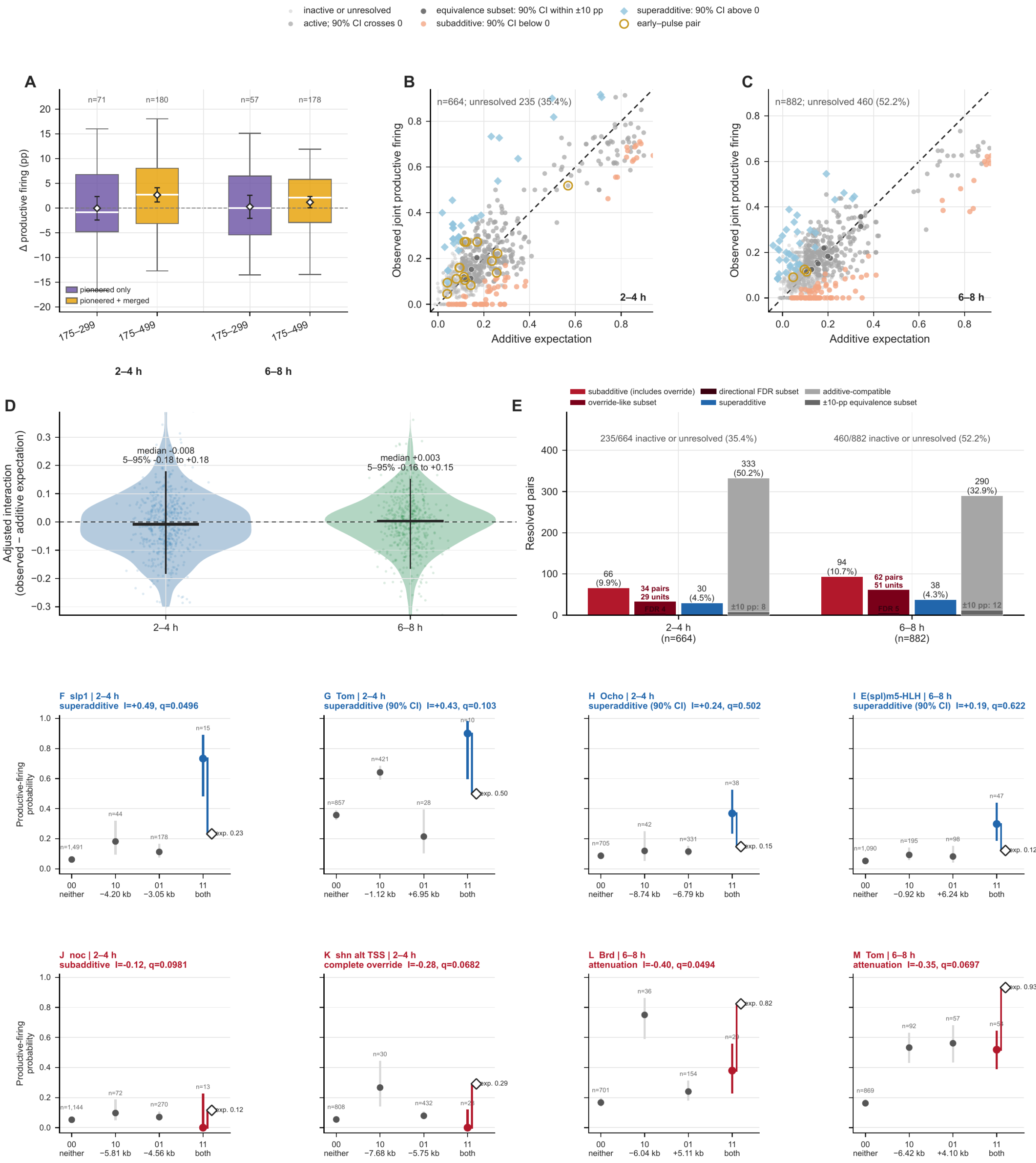

inactive or unresolved
active; 90% CI crosses 0
equivalence subset: 90% CI within ±10 pp
subadditive: 90% CI below 0
superadditive: 90% CI above 0
early–pulse pair
A
Δ productive firing (pp)
n=71
n=180
n=57
n=178
pioneered only
pioneered + merged
175–299
175–499
2–4 h
6–8 h
B
n=664; unresolved 235 (35.4%)
Observed joint productive firing
Additive expectation
2–4 h
C
n=882; unresolved 460 (52.2%)
Observed joint productive firing
Additive expectation
6–8 h
D
Adjusted interaction
(observed − additive expectation)
median -0.008
5–95% -0.18 to +0.18
median +0.003
5–95% -0.16 to +0.15
2–4 h
6–8 h
E
subadditive (includes override)
override-like subset
directional FDR subset
superadditive
additive-compatible
±10-pp equivalence subset
Resolved pairs
235/664 inactive or unresolved (35.4%)
460/882 inactive or unresolved (52.2%)
66
(9.9%)
34 pairs
29 units
30
(4.5%)
333
(50.2%)
±10 pp: 8
94
(10.7%)
62 pairs
51 units
38
(4.3%)
290
(32.9%)
±10 pp: 12
2–4 h
(n=664)
6–8 h
(n=882)
F slp1 | 2–4 h
superadditive I=+0.49, q=0.0496
G Tom | 2–4 h
superadditive (90% CI) I=+0.43, q=0.103
H Ocho | 2–4 h
superadditive (90% CI) I=+0.24, q=0.502
I E(spl)m5-HLH | 6–8 h
superadditive (90% CI) I=+0.19, q=0.622
Productive-firing
probability
n=1,491
n=44
n=178
n=15
exp. 0.23
n=857
n=421
n=28
n=10
exp. 0.50
n=705
n=42
n=331
n=38
exp. 0.15
n=1,090
n=195
n=98
n=47
exp. 0.12
00
neither
10
−4.20 kb
01
−3.05 kb
11
both
10
−1.12 kb
01
+6.95 kb
10
−8.74 kb
01
−6.79 kb
10
−0.92 kb
01
+6.24 kb
J noc | 2–4 h
subadditive I=-0.12, q=0.0981
K shn alt TSS | 2–4 h
complete override I=-0.28, q=0.0682
L Brd | 6–8 h
attenuation I=-0.40, q=0.0494
M Tom | 6–8 h
attenuation I=-0.35, q=0.0697
n=1,144
n=72
n=270
n=13
exp. 0.12
n=808
n=30
n=432
exp. 0.29
n=701
n=36
n=154
exp. 0.82
n=869
n=92
n=57
exp. 0.93
10
−5.81 kb
01
−4.56 kb
10
−7.68 kb
01
−5.75 kb
10
−6.04 kb
01
+5.11 kb
10
−6.42 kb
01
+4.10 kb

**Fig. S9. Additional shallow opening and productive-output interactions among CRM states.** The analysis uses nonredundant regulatory CRMs whose widest called boundary is ≥300 bp. (**A**) Contribution of additional small openings while holding one observed actuated CRM identity and width constant within a 9-kb upstream field. The 50–100-bp/kb group contains 450–900 bp of additional opening; pioneered-only (175–299 bp) and pioneered-or-merged (175–499 bp) doses were tested separately. **(B and C)** Observed joint productive-firing probability versus the additive expectation from the 00, 10, and 01 states for 664 adequately sampled pairs at 2–4 hours and 882 at 6–8 hours. One NFR spanning both CRMs was not counted as independent dual opening. Adjusted pointwise 90% intervals classify subadditive and superadditive candidates; active pairs whose intervals cross zero are consistent with additivity, with a nested equivalence subset requiring the interval to lie within ±10 percentage points. **(D and E)** Interaction distributions and class counts; a ±15-point sensitivity is also shown. **(F to M)** Four superadditive and four subadditive or overriding examples. Circles show 00/10/01/11 firing probabilities with binomial 95% intervals; the open diamond is the unadjusted additive expectation and the colored segment its displacement from state 11. Model estimates and $q$ values use the HC1-adjusted linear-probability model. Red examples and *slp1* pass directional FDR $q \leq 0.10$; other blue examples are exploratory pointwise candidates. These configurations illustrate the observed interaction classes.

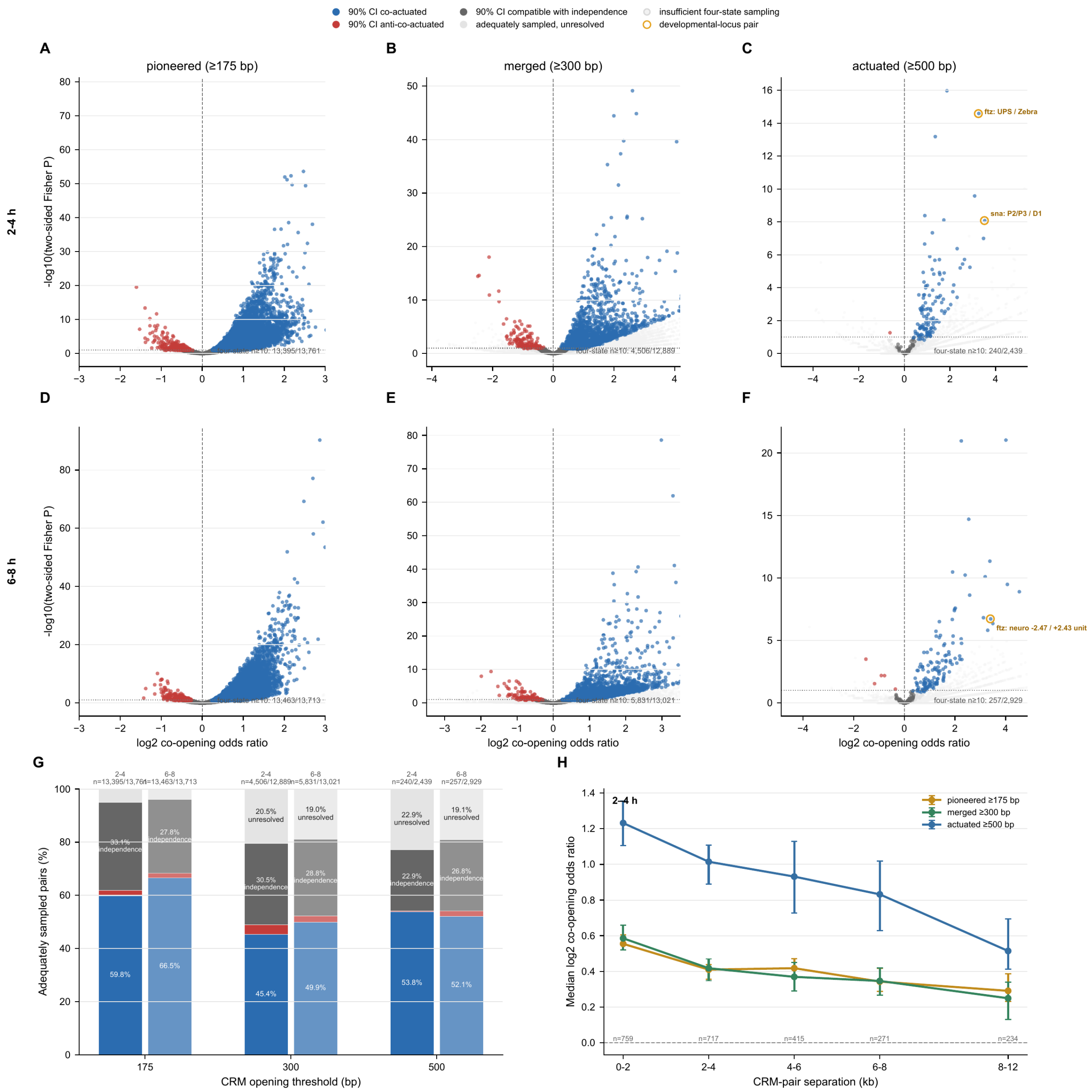


I

| CRM pair | programs | both actuated (n) | pioneered percentile | log2 OR | OR (90% CI) |
|---|---|---|---|---|---|
| ftz: UPS / Zebra | early / early | 30 | 93 | 3.24 | 9.47 (6.24–14.39) |
| sna: P2/P3 / D1 | early / early | 14 | 84 | 3.51 | 11.38 (6.19–20.92) |

**Fig. S10. CRM-pair coordination is detectable across nested opening states but often unresolved for full actuation.**

The analysis uses nonoverlapping regulatory CRM pairs with widest called boundaries ≥300 bp and molecules spanning both complete boundaries. **(A to C)** Co-opening effect and Fisher *P* value at the 175-bp pioneered, 300-bp merged, and 500-bp actuated thresholds at 2–4 hours. **(D to F)** Corresponding 6–8-hour analyses. Pointwise 90% odds-ratio intervals classify co-actuated, anti-co-actuated, independence-compatible (entirely within 0.5–2 and including 1), or unresolved pairs after requiring at least 10 molecules in each four-state cell; less-supported pairs are shown separately. (**G**) Class prevalence among adequately sampled pairs. (**H**) Median 2–4-hour co-opening odds ratio by CRM separation through 12 kb, with gene-resampling 95% intervals. (**I**) Named co-actuated pairs; counts report dual actuation and the percentile ranks the same pair's pioneered-state effect. Classifications are exploratory same-molecule associations from pooled libraries.

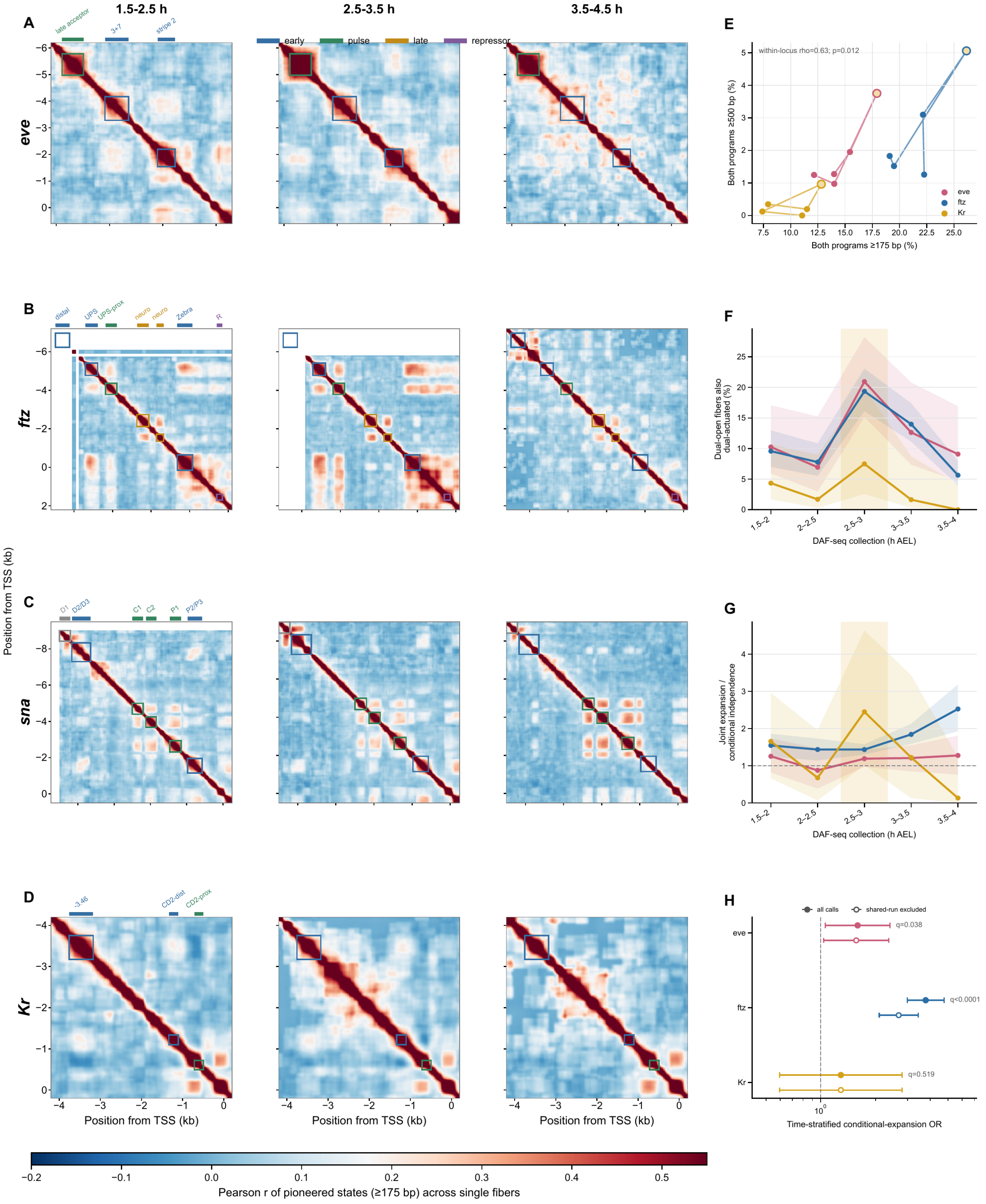

1.5-2.5 h
2.5-3.5 h
3.5-4.5 h
A
B
C
D
E
F
G
H
eve
ftz
sna
Kr
early
pulse
late
repressor
late acceptor
3+7
stripe 2
distal
UPS
UPS-prox
neuro
Zebra
R
D1
D2/D3
C1
C2
P1
P2/P3
-3.46
CD2-dist
CD2-prox
Position from TSS (kb)
Pearson r of pioneered states (≥175 bp) across single fibers
within-locus rho=0.63; p=0.012
Both programs ≥500 bp (%)
Both programs ≥175 bp (%)
Dual-open fibers also dual-actuated (%)
DAF-seq collection (h AEL)
Joint expansion / conditional independence
all calls
shared-run excluded
q=0.038
q<0.0001
q=0.519
Time-stratified conditional-expansion OR

**Fig. S11. DAF-seq resolves locus-level co-opening and a temporally concentrated transition to coordinated actuation.**

**(A to D)** Position-resolved WT DAF-seq opening correlations at *eve*, *ftz*, *sna*, and *Kr* in pooled 1.5–2.5-, 2.5–3.5-, and 3.5–4.5-hour windows. Each 25-bp position was open when it lay in an NFR ≥175 bp; pixels are Pearson correlations from ≥150 molecules spanning both positions. One-bin smoothing was used only for display, and nonoverlapping arms distinguish the *Kr* CD2 elements. (**E**) Dual actuation versus dual opening across five half-hour bins; gold marks 2.5–3 hours. (**F**) Fraction of dual-open molecules that were dual-actuated, which peaked at 2.5–3 hours at *eve* and *ftz*. (**G**) Dual expansion relative to conditional independence within dual-open molecules. (**H**) Time-stratified conditional-expansion odds ratios, with open symbols excluding one-NFR cross-program fibers. Expansion remained coordinated at *eve* and *ftz* but was unresolved at *Kr*; full estimates and intervals are printed in the panels and source data. Together, the panels resolve the temporal concentration of pairwise association on individual molecules.

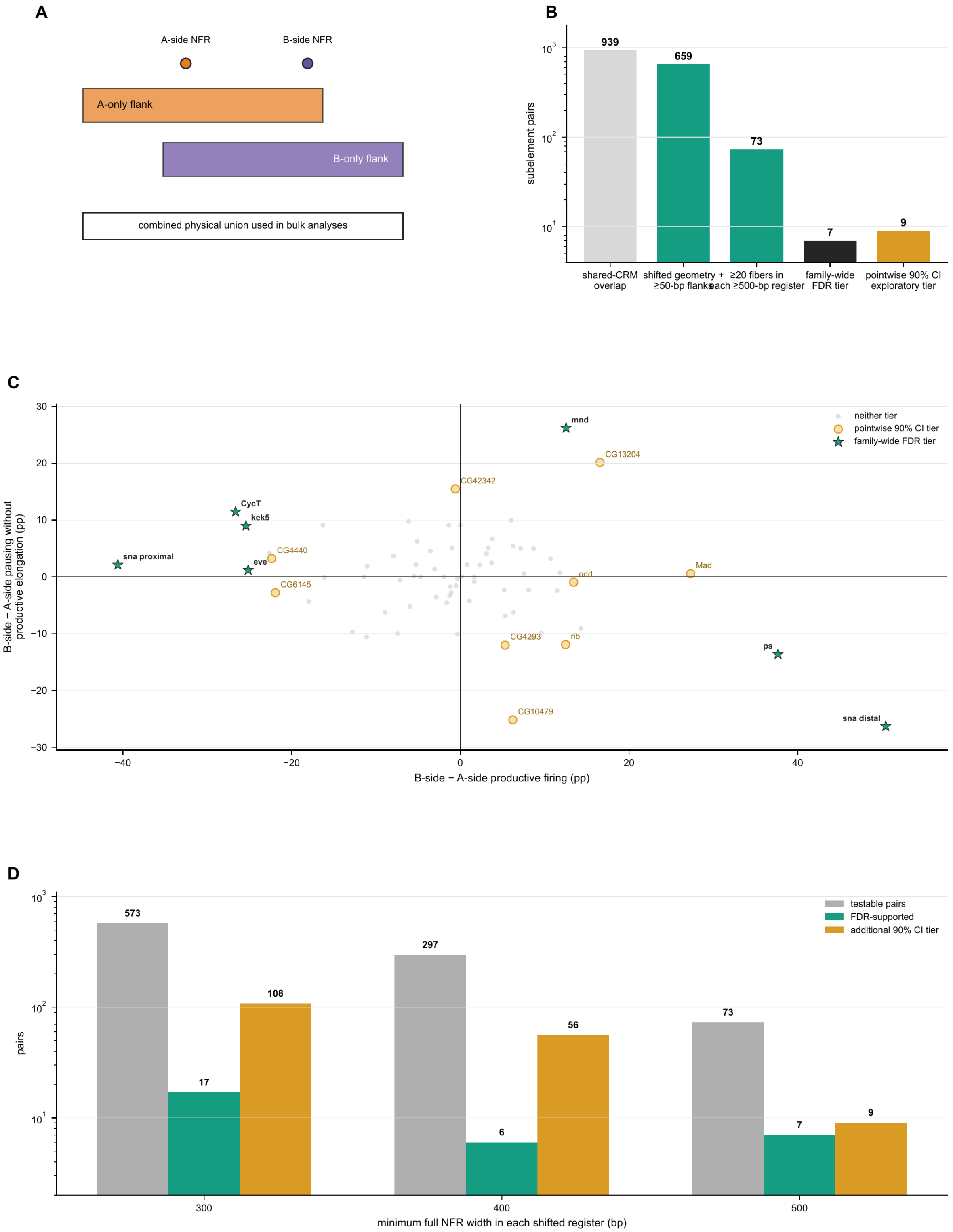
A
A-side NFR
B-side NFR
A-only flank
B-only flank
combined physical union used in bulk analyses
B
subelement pairs
939
659
73
7
9
shared-CRM overlap
shifted geometry + ≥50-bp flanks
≥20 fibers in each ≥500-bp register
family-wide FDR tier
pointwise 90% CI exploratory tier
C
neither tier
pointwise 90% CI tier
family-wide FDR tier
mnd
CG13204
CG42342
CycT
kek5
sna proximal
CG4440
eve
CG6145
odd
Mad
CG4293
rib
ps
CG10479
sna distal
B-side − A-side pausing without productive elongation (pp)
B-side − A-side productive firing (pp)
D
testable pairs
FDR-supported
additional 90% CI tier
573
17
108
297
6
56
73
7
9
pairs
minimum full NFR width in each shifted register (bp)

**Fig. S12. Shifted opening registers nominate a small set of enhancer subelements with distinct transcriptional associations.**

(**A**) Bulk analyses use a combined physical union, whereas the register test assigns a boundary-anchored NFR to the nearer prespecified subelement peak using bases unique to each side; mixed assignments are excluded. (**B**) At the prespecified 500-bp threshold, 939 shared-CRM overlapping pairs yield 659 shifted geometries, 73 pairs with at least 20 assigned fibers per register, seven pairs supported after family-wide FDR correction, and nine additional pairs supported by a deliberately exploratory pointwise 90% confidence interval plus an absolute-effect floor of 10 percentage points. The exploratory tier is used only to nominate examples. (**C**) Productive-firing and pausing-without-productive-elongation differences among the 73 identifiable 500-bp pairs. Stars denote the family-wide tier; gold points denote the additional exploratory tier. (**D**) Recovery after rerunning the same assignment and evidence rules at 300-, 400-, and 500-bp minimum full-NFR widths. Testable pairs number 573, 297, and 73; FDR-supported pairs number 17, 6, and 7; additional exploratory pairs number 108, 56, and 9. Transcriptional states were computed after masking the queried CRM from the gene-body measurement interval.

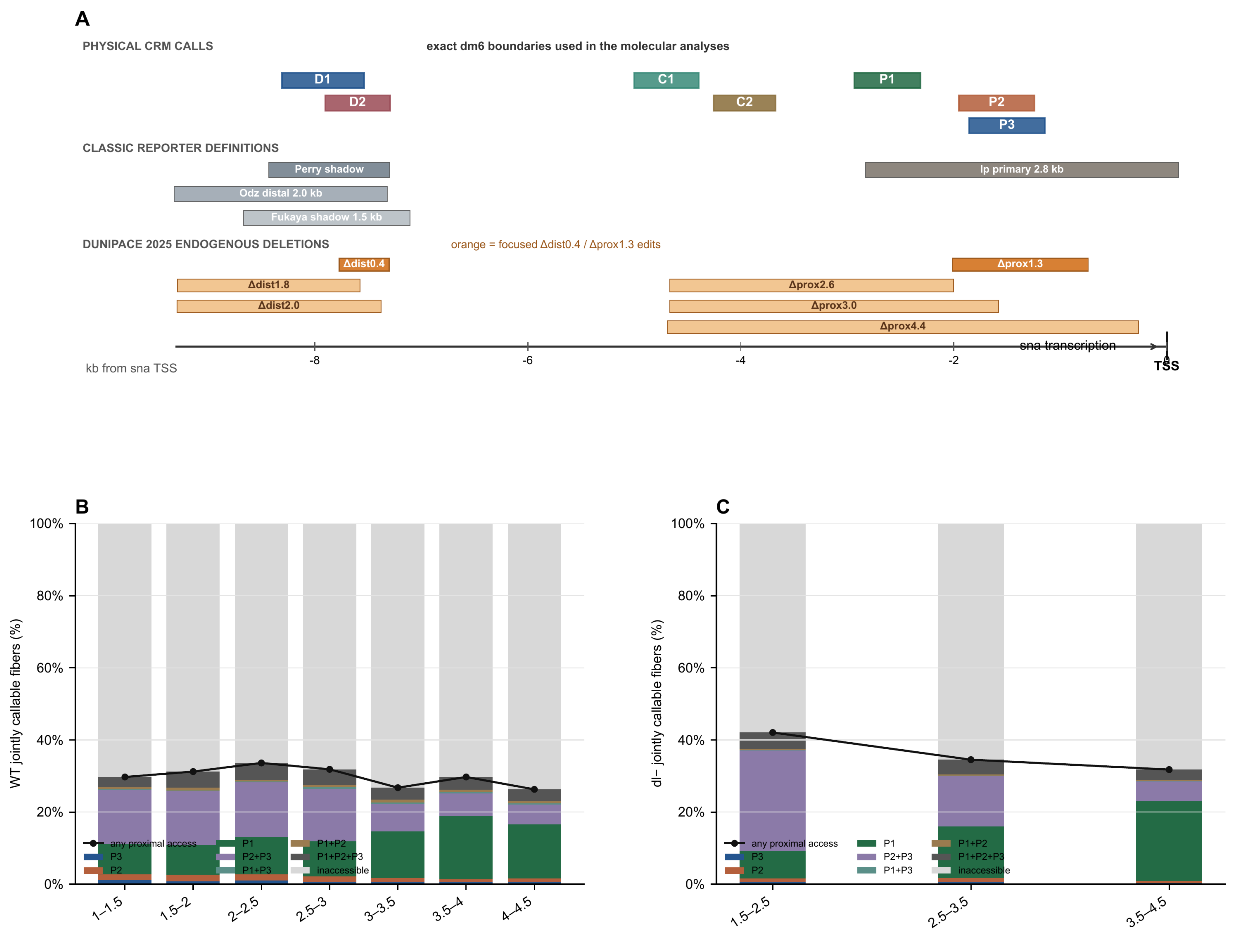
A
PHYSICAL CRM CALLS
exact dm6 boundaries used in the molecular analyses
D1
D2
C1
C2
P1
P2
P3
CLASSIC REPORTER DEFINITIONS
Perry shadow
Odz distal 2.0 kb
Fukaya shadow 1.5 kb
Ip primary 2.8 kb
DUNIPACE 2025 ENDOGENOUS DELETIONS
orange = focused Δdist0.4 / Δprox1.3 edits
Δdist0.4
Δdist1.8
Δdist2.0
Δprox1.3
Δprox2.6
Δprox3.0
Δprox4.4
sna transcription
-8
-6
-4
-2
0
TSS
kb from sna TSS
B
WT jointly callable fibers (%)
100%
80%
60%
40%
20%
0%
any proximal access
P3
P2
P1
P2+P3
P1+P3
P1+P2
P1+P2+P3
inaccessible
1–1.5
1.5–2
2–2.5
2.5–3
3–3.5
3.5–4
4–4.5
C
dl– jointly callable fibers (%)
100%
80%
60%
40%
20%
0%
any proximal access
P3
P2
P1
P2+P3
P1+P3
P1+P2
P1+P2+P3
inaccessible
1.5–2.5
2.5–3.5
3.5–4.5

**Fig. S13. The autonomous *sna* timer redistributes access within a partially accessible proximal locus.**

Every displayed element maps to a nonredundant CRM integrated across stages whose widest called boundary is at least 300 bp. CRMs overlapping annotated insulators or chromatin boundaries were excluded: an element was ineligible when its peak overlapped a chromatin-boundary interval defined by Batut et al. (*52*) or it mapped to a curated functional element labeled as an insulator or boundary. All seven displayed *sna* elements pass this rule. (**A**) D1, D2, C1, C2, P1, P2, and P3 physical CRM calls in dm6, classic *sna* reporter definitions, and the endogenous distal- and proximal-region deletions reported by Dunipace et al. Exact interval coordinates and CRM overlaps are listed in Table S5. The proximal deletion spans both P2 and P3. (**B**) Absolute WT composition of jointly callable fibers across P1/P2/P3 configurations using the 175-bp opening threshold. Gray denotes fibers with no proximal opening. (**C**) Corresponding absolute composition in *dorsal*-null embryos. The black lines in B and C give total proximal accessibility; including inaccessible fibers shows that the total accessible fraction changes modestly while the balance among accessible configurations is redistributed.

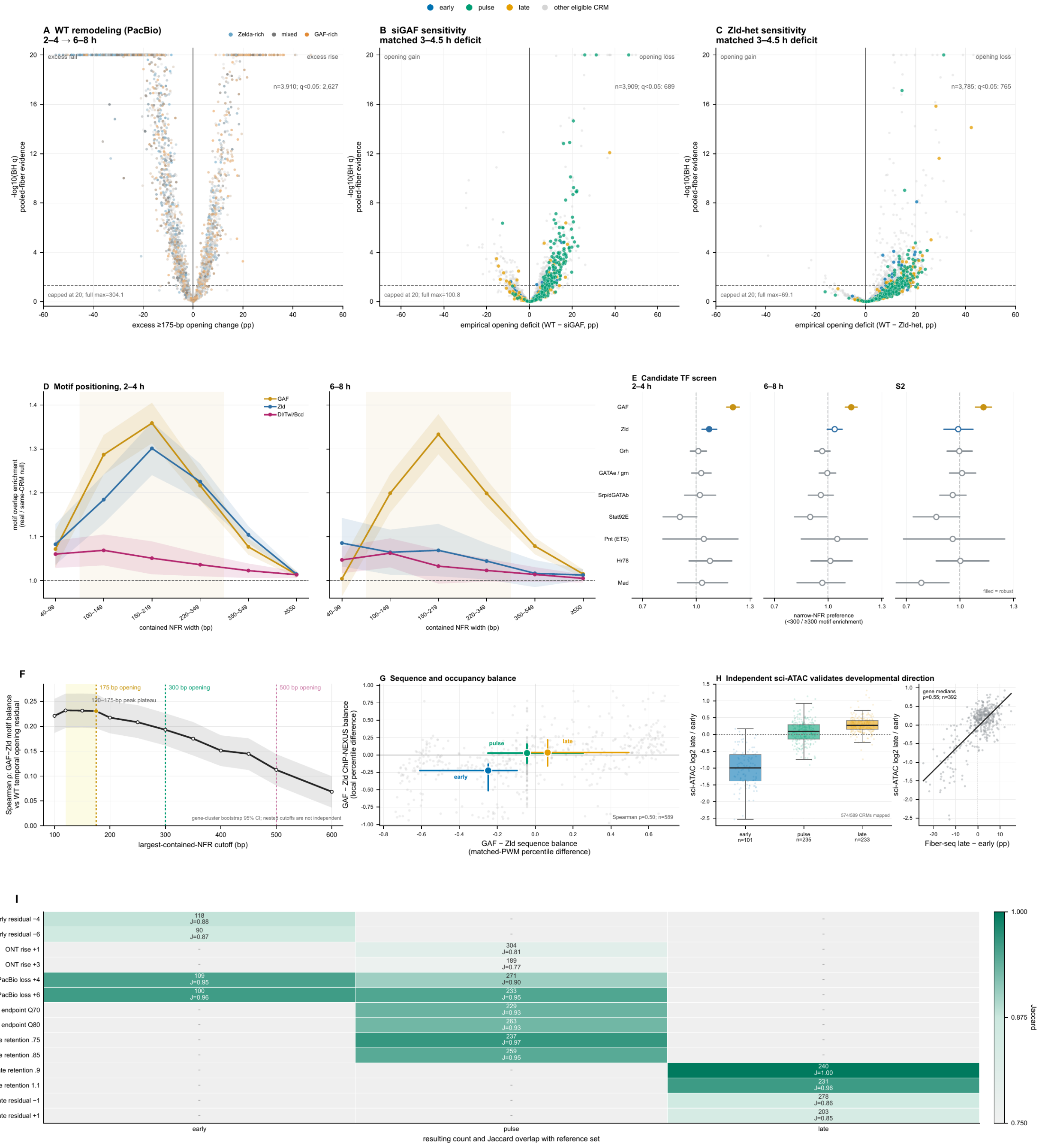

early
pulse
late
other eligible CRM
A WT remodeling (PacBio) 2–4 → 6–8 h
Zelda-rich
mixed
GAF-rich
excess fall
excess rise
n=3,910; q<0.05: 2,627
capped at 20; full max=304.1
-log10(BH q) pooled-fiber evidence
excess ≥175-bp opening change (pp)
B siGAF sensitivity matched 3–4.5 h deficit
opening gain
opening loss
n=3,909; q<0.05: 689
capped at 20; full max=100.8
empirical opening deficit (WT − siGAF, pp)
C Zld-het sensitivity matched 3–4.5 h deficit
n=3,785; q<0.05: 765
capped at 20; full max=69.1
empirical opening deficit (WT − Zld-het, pp)
D Motif positioning, 2–4 h
6–8 h
GAF
Zld
Dl/Twi/Bcd
motif overlap enrichment (real / same-CRM null)
contained NFR width (bp)
E Candidate TF screen 2–4 h
6–8 h
S2
GAF
Zld
Grh
GATAe / grn
Srp/dGATAb
Stat92E
Pnt (ETS)
Hr78
Mad
filled = robust
narrow-NFR preference (<300 / ≥300 motif enrichment)
F
175 bp opening
300 bp opening
500 bp opening
120–175-bp peak plateau
Spearman ρ: GAF−Zld motif balance vs WT temporal opening residual
gene-cluster bootstrap 95% CI; nested cutoffs are not independent
largest-contained-NFR cutoff (bp)
G Sequence and occupancy balance
GAF − Zld ChIP-NEXUS balance (local percentile difference)
Spearman ρ=0.50; n=589
GAF − Zld sequence balance (matched-PWM percentile difference)
H Independent sci-ATAC validates developmental direction
sci-ATAC log2 late / early
574/589 CRMs mapped
early n=101
pulse n=235
late n=233
gene medians ρ=0.55; n=392
Fiber-seq late − early (pp)
I
early residual −4
early residual −6
ONT rise +1
ONT rise +3
PacBio loss +4
PacBio loss +6
pulse endpoint Q70
pulse endpoint Q80
pulse retention .75
pulse retention .85
late retention .9
late retention 1.1
late residual −1
late residual +1
118 J=0.88
90 J=0.87
304 J=0.81
189 J=0.77
109 J=0.95
271 J=0.90
100 J=0.96
233 J=0.95
229 J=0.93
263 J=0.93
237 J=0.97
259 J=0.95
240 J=1.00
231 J=0.96
278 J=0.86
203 J=0.85
resulting count and Jaccard overlap with reference set
Jaccard

**Fig. S14. Developmental remodeling and pioneer-factor dependence across the CRM continuum.**

(**A**) WT change in ≥175-bp opening from 2–4 to 6–8 hours. (**B**) Direct 3–4.5-hour siGAF deficits; colored points denote 104 early CRMs, 243 pulse CRMs, and 239 late CRMs. (**C**) Direct 3–4.5-hour maternal *zld*-heterozygote deficits; colored points denote 103 early CRMs, 233 pulse CRMs, and 232 late CRMs. (**D**) Gene-balanced enrichment of GAF, Zld, and pooled Dl/Twi/Bcd motifs in exact single-fiber NFRs relative to eight same-width within-CRM shuffles. (**E**) Narrow-versus-wide NFR specificity for occupancy-supported candidate-factor motifs; filled points pass enrichment and specificity criteria at 5, 10, and 20 NFRs per gene. (**F**) GAF-minus-Zld motif balance versus WT temporal opening across candidate NFR cutoffs. (**G**) Sequence versus stage-5 ChIP-NEXUS binding balance across temporal CRMs. (**H**) Calderon et al. sci-ATAC validation of developmental direction across temporal classes. (**I**) Temporal-class sensitivity to nearby gates. Residual rows vary stage-adjusted temporal-change gates, endpoint rows vary the late-stage opening quantile, and retention rows vary the 6–8-hour/2–4-hour opening ratio; each row label gives the tested cutoff. Analyses begin with the 6,942-CRM ≥300-bp census and exclude 335 insulator/boundary CRMs; panel-specific motif and depth filters are shown. In D and E, <300 bp describes a molecule-level NFR, distinct from CRM eligibility. Intervals quantify the sampling unit indicated in each panel.

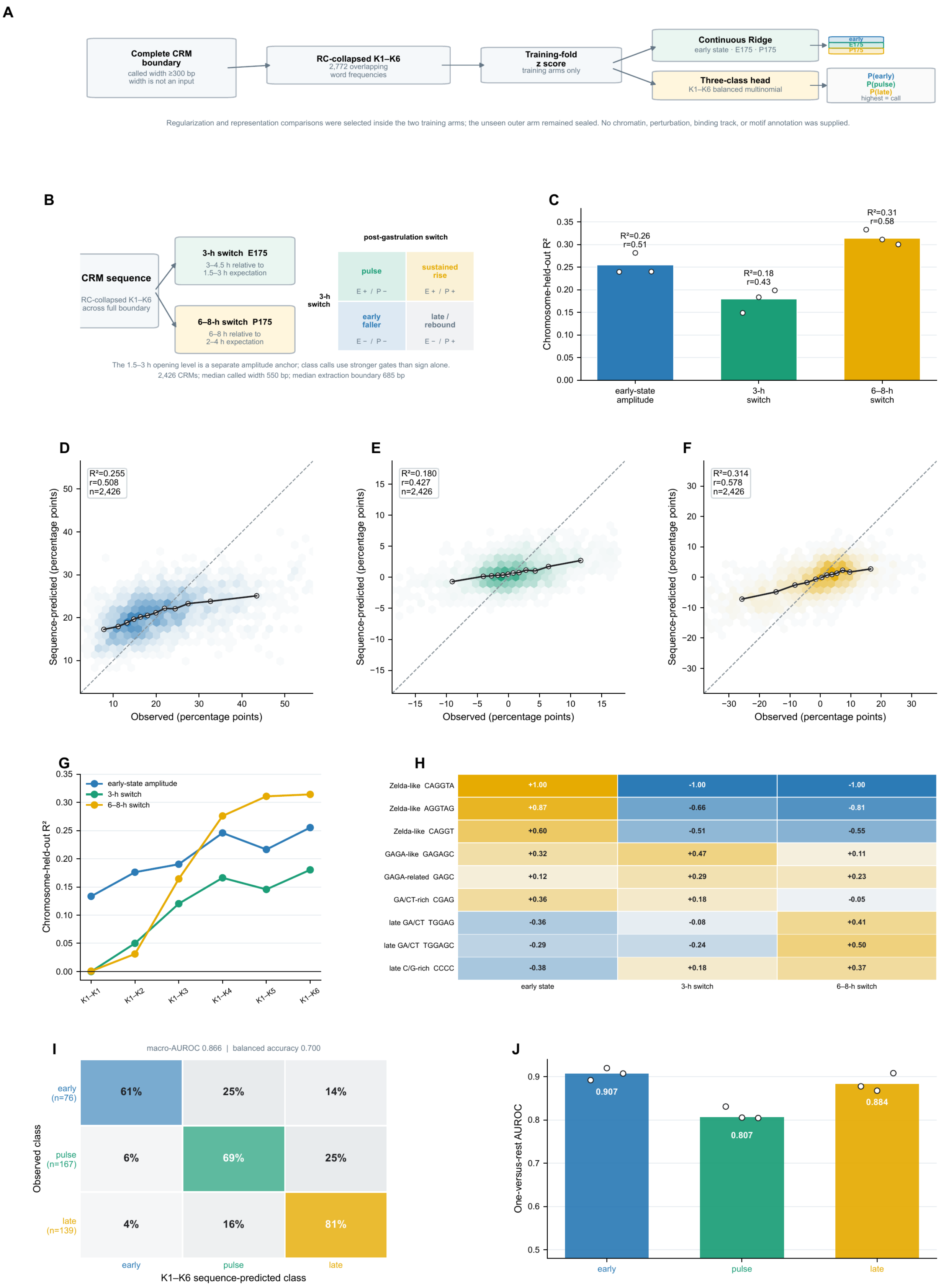
A
Complete CRM boundary
called width ≥300 bp
width is not an input
RC-collapsed K1–K6
2,772 overlapping word frequencies
Training-fold z score
training arms only
Continuous Ridge
early state · E175 · P175
Three-class head
K1–K6 balanced multinomial
P(early)
P(pulse)
P(late)
highest = call
Regularization and representation comparisons were selected inside the two training arms; the unseen outer arm remained sealed. No chromatin, perturbation, binding track, or motif annotation was supplied.
B
CRM sequence
RC-collapsed K1–K6 across full boundary
3-h switch E175
3–4.5 h relative to 1.5–3 h expectation
6–8-h switch P175
6–8 h relative to 2–4 h expectation
post-gastrulation switch
3-h switch
pulse
E + / P −
sustained rise
E + / P +
early faller
E − / P −
late / rebound
E − / P +
The 1.5–3 h opening level is a separate amplitude anchor; class calls use stronger gates than sign alone.
2,426 CRMs; median called width 550 bp; median extraction boundary 685 bp
C
Chromosome-held-out R²
R²=0.26
r=0.51
R²=0.18
r=0.43
R²=0.31
r=0.58
early-state amplitude
3-h switch
6–8-h switch
D
R²=0.255
r=0.508
n=2,426
E
R²=0.180
r=0.427
n=2,426
F
R²=0.314
r=0.578
n=2,426
Sequence-predicted (percentage points)
Observed (percentage points)
G
Chromosome-held-out R²
early-state amplitude
3-h switch
6–8-h switch
K1–K1
K1–K2
K1–K3
K1–K4
K1–K5
K1–K6
H
Zelda-like CAGGTA +1.00 -1.00 -1.00
Zelda-like AGGTAG +0.87 -0.66 -0.81
Zelda-like CAGGT +0.60 -0.51 -0.55
GAGA-like GAGAGC +0.32 +0.47 +0.11
GAGA-related GAGC +0.12 +0.29 +0.23
GA/CT-rich CGAG +0.36 +0.18 -0.05
late GA/CT TGGAG -0.36 -0.08 +0.41
late GA/CT TGGAGC -0.29 -0.24 +0.50
late C/G-rich CCCC -0.38 +0.18 +0.37
early state
3-h switch
6–8-h switch
I
macro-AUROC 0.866 | balanced accuracy 0.700
early (n=76) 61% 25% 14%
pulse (n=167) 6% 69% 25%
late (n=139) 4% 16% 81%
Observed class
early
pulse
late
K1–K6 sequence-predicted class
J
One-versus-rest AUROC
0.907
0.807
0.884
early
pulse
late

**Fig. S15. Sequence-only prediction of CRM opening and timing.**

(**A**) Sequence-only architecture using normalized reverse-complement-collapsed 1–6-mer frequencies from complete CRM boundaries; CRM width was excluded. Models and regularization were selected within the two outer-training chromosome arms before evaluation on the third. (**B**) Early-state amplitude plus two signed residual switches: E175 for 1.5–3 to 3–4.5 hours and P175 for 2–4 to 6–8 hours. Their signs define conceptual pulse, sustained-rise, early-fall, and late/rebound sectors; paper classes are stringent subsets. (**C**) Chromosome-held-out prediction of amplitude, E175, and P175. **(D to F)** Observed versus out-of-fold predictions; connected bins illustrate range compression. (**G**) Performance across maximum word lengths. (**H**) Stable predictive word-family coefficients. **(I and J)** Held-chromosome temporal-class confusion and one-versus-rest AUROC. The continuous analysis used 2,426 architecture-clean CRMs on chr2L, chr3L, and chr3R; 382 of 589 class exemplars lay on these modeled arms. Inputs comprised CRM-boundary sequence alone.

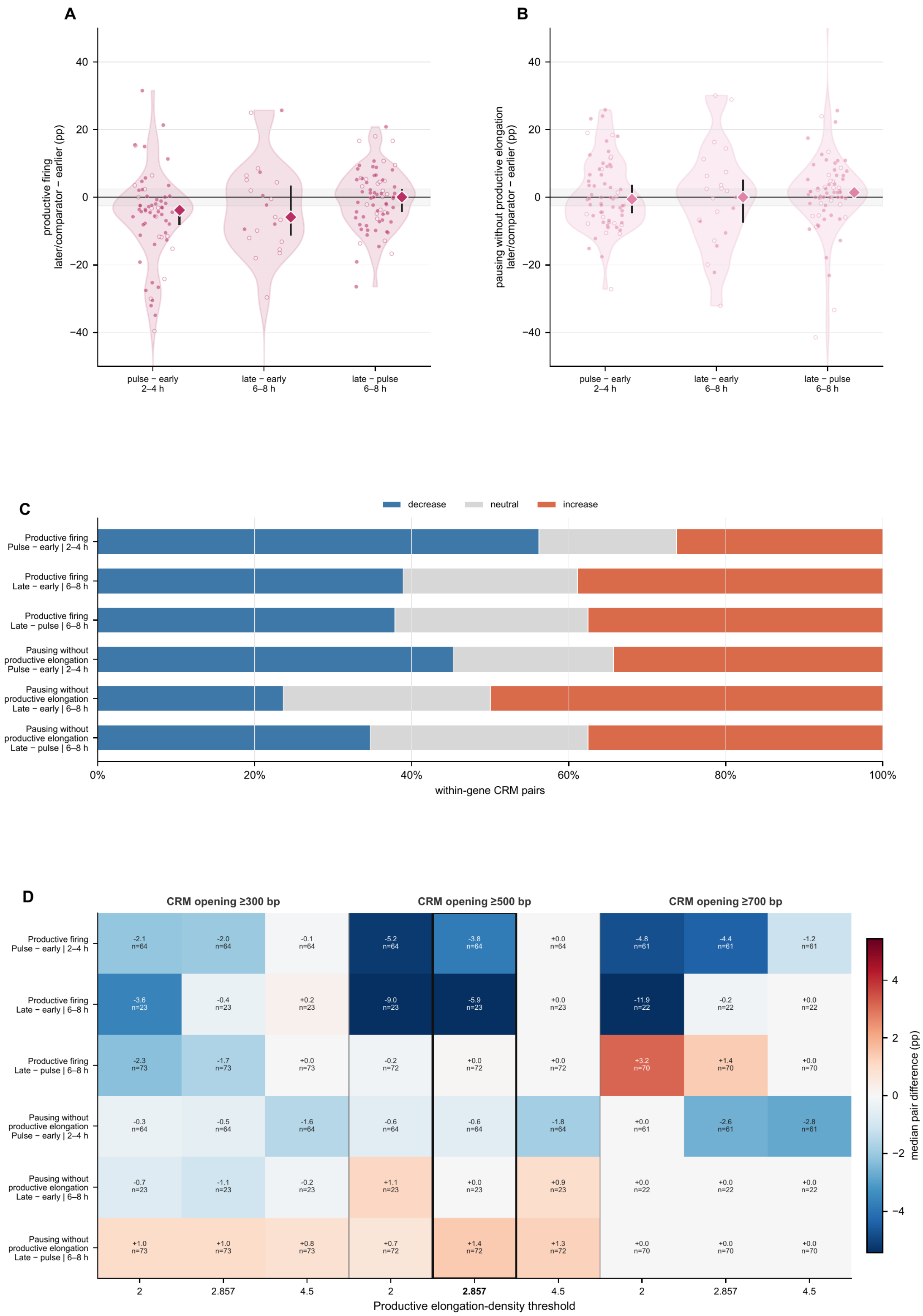

A
productive firing
later/comparator − earlier (pp)
pulse − early
2–4 h
late − early
6–8 h
late − pulse
6–8 h
B
pausing without productive elongation
later/comparator − earlier (pp)
C
decrease
neutral
increase
Productive firing
Pulse − early | 2–4 h
Productive firing
Late − early | 6–8 h
Productive firing
Late − pulse | 6–8 h
Pausing without
productive elongation
Pulse − early | 2–4 h
Pausing without
productive elongation
Late − early | 6–8 h
Pausing without
productive elongation
Late − pulse | 6–8 h
within-gene CRM pairs
D
CRM opening ≥300 bp
CRM opening ≥500 bp
CRM opening ≥700 bp
-2.1 n=64, -2.0 n=64, -0.1 n=64, -5.2 n=64, -3.8 n=64, +0.0 n=64, -4.8 n=61, -4.4 n=61, -1.2 n=61
-3.6 n=23, -0.4 n=23, +0.2 n=23, -9.0 n=23, -5.9 n=23, +0.0 n=23, -11.9 n=22, -0.2 n=22, +0.0 n=22
-2.3 n=73, -1.7 n=73, +0.0 n=73, -0.2 n=72, +0.0 n=72, +0.0 n=72, +3.2 n=70, +1.4 n=70, +0.0 n=70
-0.3 n=64, -0.5 n=64, -1.6 n=64, -0.6 n=64, -0.6 n=64, -1.8 n=64, +0.0 n=61, -2.6 n=61, -2.8 n=61
-0.7 n=23, -1.1 n=23, -0.2 n=23, +1.1 n=23, +0.0 n=23, +0.9 n=23, +0.0 n=22, +0.0 n=22, +0.0 n=22
+1.0 n=73, +1.0 n=73, +0.8 n=73, +0.7 n=72, +1.4 n=72, +1.3 n=72, +0.0 n=70, +0.0 n=70, +0.0 n=70
Productive elongation-density threshold
median pair difference (pp)

**Fig. S16. Timed CRM exchange and transcription-associated output.**

(**A**) Later-minus-earlier within-gene CRM-pair differences in productive firing for 64 pulse-versus-early pairs at 2–4 hours, 23 late-versus-early pairs at 6–8 hours, and 72 late-versus-pulse pairs at 6–8 hours. All constituent units satisfy regulatory eligibility, exclude annotated insulators and boundaries, and have a widest member CRM boundary of at least 300 bp. Points are CRM pairs; filled points have at least 20 fibers in each mutually exclusive opening arm and open points have 1–19. Diamonds show pair medians and intervals that resample genes as clusters. (**B**) Corresponding differences in pausing without productive elongation. (**C**) Decrease, neutral, and increase fractions for the same pair effects; neutral is ±2.5 percentage points. (**D**) Sensitivity of median pair differences to 300-, 500-, and 700-bp focal CRM-opening thresholds and productive-elongation-density thresholds of 2, 2.857, and 4.5 eligible Pol-II-associated footprints per kilobase. Columns are grouped by the molecular CRM-opening threshold, distinct from the ≥300-bp CRM-boundary eligibility requirement; the black outline marks the primary 500-bp/2.857 definition. Timed classes show heterogeneous transcriptional associations.

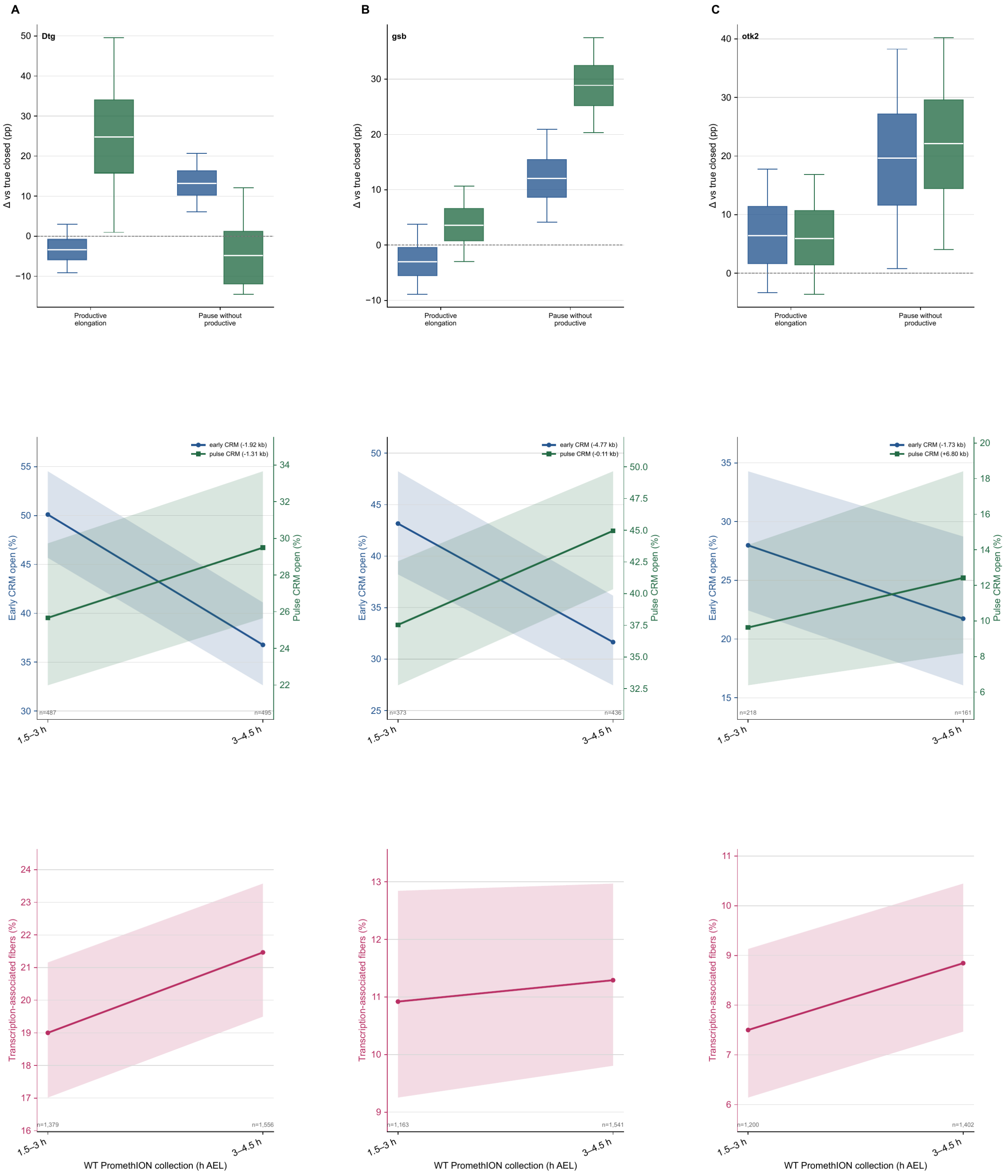
A
B
C
Dtg
gsb
otk2
Δ vs true closed (pp)
Productive elongation
Pause without productive
early CRM (-1.92 kb)
pulse CRM (-1.31 kb)
early CRM (-4.77 kb)
pulse CRM (-0.11 kb)
early CRM (-1.73 kb)
pulse CRM (+6.80 kb)
Early CRM open (%)
Pulse CRM open (%)
n=487
n=495
n=373
n=436
n=218
n=161
1.5–3 h
3–4.5 h
Transcription-associated fibers (%)
n=1,379
n=1,556
n=1,163
n=1,541
n=1,200
n=1,402
WT PromethION collection (h AEL)

**Fig. S17. Additional pulse CRM examples.**

(A to C) *Dtg*, *gsb*, and *otk2*. Top, independent 2–4-hour PacBio Fiber-seq comparisons of productive elongation and pausing without productive elongation on fibers carrying the early or pulse CRM relative to fibers on which the focal CRM is closed. Boxes show the bootstrap median and interquartile range; whiskers span the 5th–95th percentiles. Middle, ONT Fiber-seq opening frequencies of the early and pulse CRMs at 1.5–3 and 3–4.5 hours. Bottom, fraction of ONT gene-spanning fibers carrying at least one initiation-complex-, paused-Pol-II-, or elongating-Pol-II-sized protection in the corresponding transcription window. Blue denotes the early CRM, green denotes the pulse CRM, and magenta denotes transcription-associated fibers. Intervals quantify sampling from pooled molecular collections.

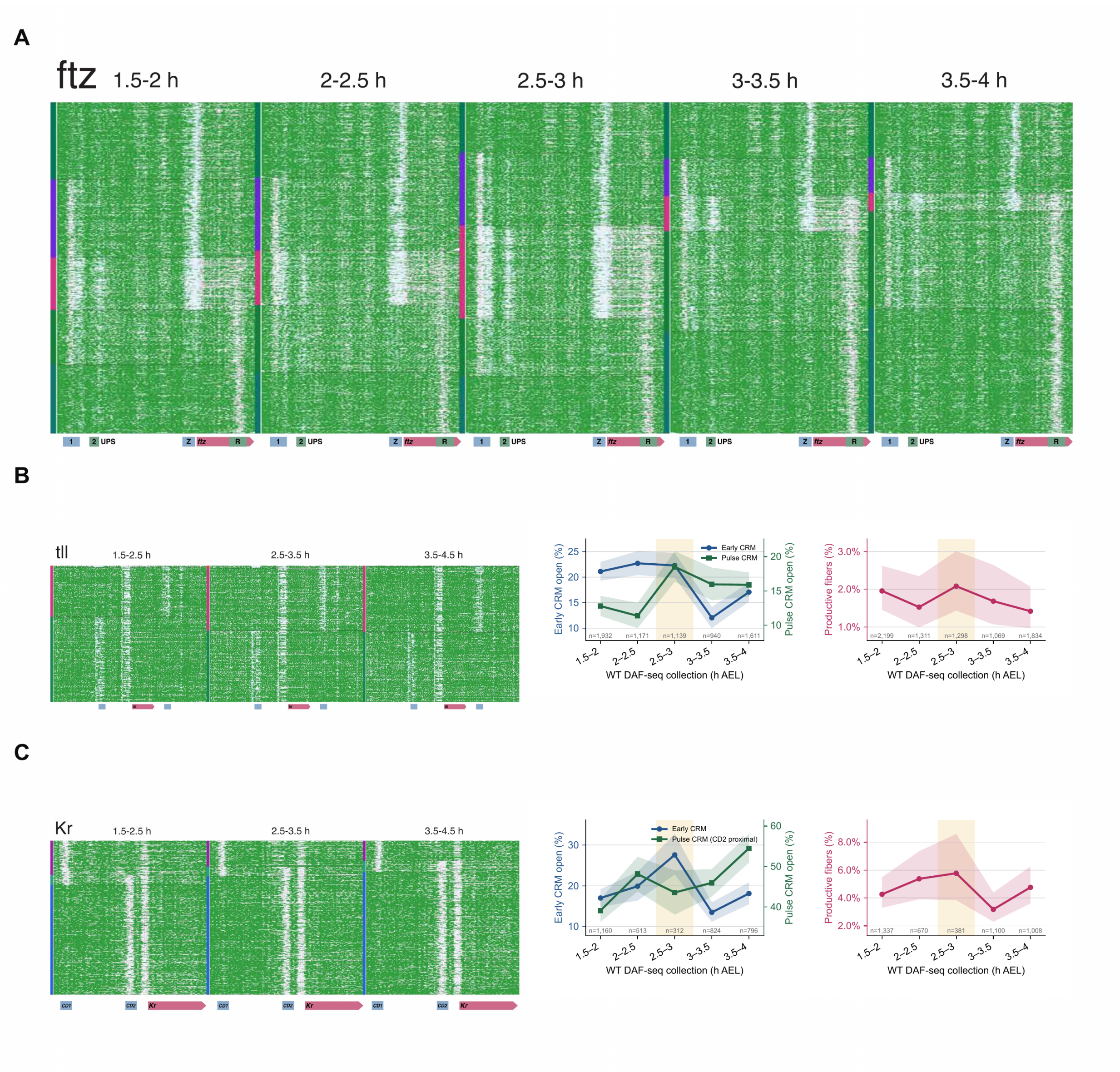
A
ftz
1.5-2 h
2-2.5 h
2.5-3 h
3-3.5 h
3.5-4 h
1
2 UPS
Z ftz R
B
tll
1.5-2.5 h
2.5-3.5 h
3.5-4.5 h
Early CRM
Pulse CRM
Early CRM open (%)
Pulse CRM open (%)
Productive fibers (%)
WT DAF-seq collection (h AEL)
C
Kr
CD1
CD2
Kr
Pulse CRM (CD2 proximal)

**Fig. S18. DAF-seq time courses at *ftz*, *tll*, and *Kr*.**

(**A**) Position-resolved WT DAF-seq fibers at *ftz* across five half-hour collections from 1.5 to 4 hours. Element and gene annotations are shown below each collection; quantitative *ftz* opening and productive-output trajectories are shown in Fig. 4. (B and C) Position-resolved WT DAF-seq fibers at *tll* and *Kr* in pooled 1.5–2.5-, 2.5–3.5-, and 3.5–4.5-hour windows (left), early- and pulse-program opening across five half-hour collections (middle), and full-gene productive-firing frequency (right). Blue denotes early-program opening, green denotes pulse-program opening, and magenta denotes productive firing. *Kr* compares CD1 with the CD2-proximal pulse element and is shown in chronological source-collection order. CRM opening is the fraction of jointly callable fibers carrying an NFR of at least 175 bp at the indicated program; productive firing requires at least 2.857 eligible Pol-II-associated footprints per kilobase among gene-callable fibers. Ribbons are molecule-sampling 95% Wilson intervals, and the shaded 2.5–3-hour window marks the corresponding collection. The position-resolved panels display only fibers carrying at least one active regulatory element in the plotted locus; quantitative opening and transcriptional rates use the full jointly callable fiber set.

| Assay | Genotype or perturbation | Developmental window | Platform | Retained nuclear call-track coverage | Aligned span p50/p95 (kb) |
|---|---|---|---|---|---|
| Fiber-seq | WT | 2–4 h | PacBio | 2,375.5× (16,383,316 records) | 19.9 / 28.1 |
| Fiber-seq | WT | 6–8 h | PacBio | 2,414.1× (16,587,728 records) | 19.9 / 28.1 |
| Fiber-seq | WT | 1.5–3 h | ONT | 491.7× (6,067,058 records) | 9.2–11.3 / 60.3–65.7 |
| Fiber-seq | WT | 3–4.5 h | ONT | 462.5× (6,444,175 records) | 9.2–11.3 / 60.3–65.7 |
| Fiber-seq | GAF depleted | 3–4.5 h | ONT | 195.2× (3,107,984 records) | 9.3 / 46.3 |
| Fiber-seq | Maternal *zld* heterozygote | 3–4.5 h | ONT | 124.4× (2,234,721 records) | 5.5 / 40.4 |
| DAF-seq | WT | 1–4.5 h; seven 0.5-h bins, plus pooled 2–4 h | ONT | targeted | 1.8–12.3 / 6.0–16.9; pooled rho, 6.0 / 6.0 |
| DAF-seq | dorsal-null | three 1-h bins, plus pooled 2–4 h | ONT | targeted | 6.0–12.3 / 6.0–13.6; pooled rho, 6.0 / 6.0 |

**Table S1.**
Samples, sequencing sources, retained call sets, and aligned spans.

| Assay | Genotype or perturbation | Developmental window | Analysis role | Call-track and wet-lab metadata |
|---|---|---|---|---|
| Fiber-seq | WT | 2–4 h | Core chromatin/transcription map; CRM model | Pooled independently collected embryo/Hia5 reactions (T.W.T.; pool structure in Study Design); FiberHMM 2.16.7; Hia5/PacBio model; nucleosome and factor-footprint recall; 11 technical movies; duplicated movie m84046_250513_034312_s1 excluded |
| Fiber-seq | WT | 6–8 h | Long temporal axis | Pooled independently collected embryo/Hia5 reactions (T.W.T.; pool structure in Study Design); FiberHMM 2.16.7; Hia5/PacBio model; nucleosome and factor-footprint recall; 11 technical movies; duplicated movie m84046_250513_034312_s1 excluded |
| Fiber-seq | WT | 1.5–3 h | Pre-gastrulation short-axis state | Pooled embryo/Hia5 reactions generated over two days (C.C. and T.W.T.; pool structure in Study Design); FiberHMM 2.16.7; Hia5/Nanopore model; topology-constrained nucleosome recall and factor-footprint recall |
| Fiber-seq | WT | 3–4.5 h | Gastrulation short-axis state | Pooled embryo/Hia5 reactions generated over two days (C.C. and T.W.T.; pool structure in Study Design); FiberHMM 2.16.7; Hia5/Nanopore model; topology-constrained nucleosome recall and factor-footprint recall |
| Fiber-seq | GAF depleted | 3–4.5 h | Same-window pioneer perturbation | Pooled embryo/Hia5 reactions generated over two days (C.C. and T.W.T.; pool structure in Study Design); FiberHMM 2.16.7; Hia5/Nanopore model; topology-constrained nucleosome recall and factor-footprint recall |
| Fiber-seq | Maternal *zld* heterozygote | 3–4.5 h | Same-window pioneer perturbation | Pooled embryo/Hia5 reactions generated over two days (C.C. and T.W.T.; pool structure in Study Design); FiberHMM 2.16.7; Hia5/Nanopore model; topology-constrained nucleosome recall and factor-footprint recall |
| DAF-seq | WT | 1–4.5 h; seven 0.5-h bins, plus pooled 2–4 h | Locus time course and pooled rho comparison | Pooled embryo/DAF reactions generated over two days (C.C. and T.W.T.; pool structure in Study Design); FiberHMM 2.16.7; DddB/Nanopore DAF model; nucleosome and factor-footprint recall; deamination-fingerprint deduplication at Jaccard ≥0.95 |
| DAF-seq | dorsal-null | three 1-h bins, plus pooled 2–4 h | Spatial-input autonomy | Pooled embryo/DAF reactions generated over two days (C.C. and T.W.T.; pool structure in Study Design); FiberHMM 2.16.7; DddB/Nanopore DAF model; nucleosome and factor-footprint recall; deamination-fingerprint deduplication at Jaccard ≥0.95 |

**Table S1 continued.**
Analysis roles and call-track metadata.

| Target | Amplicon | dm6 interval | Modal span (kb) | Aligned span p50/p95 (kb) | Collections represented |
|---|---|---|---|---|---|
| eve | short | chr2R:9,972,169-9,978,245 | 6.1 | 6.1 / 6.1 | WT |
| eve | long | chr2R:9,972,459-9,989,315 | 16.9 | 1.8 / 16.9 | WT |
| ftz | default | chr3R:6,858,106-6,867,158 | 9.1 | 8.7 / 9.1 | WT |
| Kr | default | chr2R:25,221,309-25,231,948 | 10.6 | 3.4 / 10.6 | WT |
| rho | default | chr3L:1,459,955-1,465,925 | 6.0 | 6.0 / 6.0 | WT; dorsal-null |
| sna | short | chr2L:15,474,794-15,484,544 | 9.8 | 9.7 / 9.8 | WT |
| sna | long | chr2L:15,474,793-15,488,379 | 13.6 | 12.3 / 13.6 | WT; dorsal-null |
| tll | default | chr3R:30,845,691-30,859,086 | 13.4 | 5.0 / 13.4 | WT |
| zen | default | chr3R:6,749,971-6,760,141 | 10.2 | 10.2 / 10.2 | WT; dorsal-null |

**Table S1 continued.**
DAF-seq amplicons analyzed.

| Target | Amplicon | Forward primer | Forward sequence (5′→3′) | Reverse primer | Reverse sequence (5′→3′) |
|---|---|---|---|---|---|
| eve | short | eve_short_fwd | AAGCACTGCTTAACACCTTCAACAC | eve_short_rev | CCCCTAATCCCTTCGACATCATTAT |
| eve | long | eve_long_fwd | CTTTGGCCTAATACCCAAGCAACTG | eve_long_rev | CAGTTCACTTACCCCTTGGAATCTG |
| ftz | default | ftz_fwd | AACTGACCTAATTTACGCCACACAC | ftz_rev | CCACCAATTCGCACCTAACACTAAA |
| Kr | default | Kr_fwd | GCCGAACCCCTCATCATATCTAAAA | Kr_rev | CGCCCAATCCATCCGTTTACTTTTA |
| rho | default | rho_fwd | CCACACCCTGCTTGCCCCA | rho_rev | CGCTTTCCGCCCCCTATATCATTTT |
| sna | short | sna_short_fwd | TCCTCTTCTGTCCCTCGTATTTC | sna_short_rev | CCCCCACCAGACACCCATT |
| sna | long | sna_long_fwd | ATCCTCTTCTGTCCCTCGTATTTCT | sna_long_rev | CACCTTCAACTAGGTCAATACTCCA |
| tll | default | tll_fwd | CATCGACCTCGAAATTCCCCC | tll_rev | GTTCAACTCGCTAACTACCCCTTTT |
| zen | default | zen_fwd | CCCAGAAACCAATCTATCACCCTAT | zen_rev | AACTCGATCTCCACCATCCTTATTC |

**Table S2.**
DAF-seq primer sequences.

| Term or outcome | Operational definition | Interpretation limit |
|---|---|---|
| Pioneered | centered NFR ≥175 bp | Operational pioneered state |
| Merged | centered NFR ≥300 bp | Operational merged state |
| Actuated | centered NFR ≥500 bp | Operational actuated state |
| Productive firing | elongation density ≥2.857 eligible Pol-II-associated footprints per kilobase | Binary productive-elongation outcome |
| Conditional elongating-Pol-II load | elongation density among productively firing molecules | Secondary conditional outcome; subject to selection |
| Independent 11 state | two separately boundary-matched NFRs | One NFR spanning both CRMs is shared, not 11 |

**Table S3.**
Operational state definitions.

| Resolution or analysis set | Count | Meaning |
|---|---|---|
| Cross-stage CRM catalog | 12,121 | CRM census across 1,276 genes |
| CRMs within 10 kb of linked TSS | 11,115 | CRM catalog and promoter links in the archived source data (*49*) |
| Nonredundant CRMs with called boundary ≥300 bp | 6,942 | Common CRM universe used for REDfly and selected regulatory analyses |
| Temporal-analysis CRM set | 5,449 | Adequately covered regulatory CRMs outside annotated insulators and chromatin boundaries |
| Main ONT temporal-comparison set | 3,912 | Temporal-analysis set after the common ≥300-bp boundary filter |
| Early CRMs | 104 | Stringent WT-defined temporal landmarks after the common boundary filter |
| Pulse CRMs | 245 | Rise during gastrulation followed by later decline |
| Late CRMs | 240 | Nondeclining late comparator selected from WT measurements |
| siGAF-supported early/pulse/late | 104 / 243 / 239 | Direct 3–4.5-hour empirical-deficit comparison |
| *zld*-heterozygote-supported early/pulse/late | 103 / 233 / 232 | Direct 3–4.5-hour empirical-deficit comparison |

**Table S4.**
CRM count cascade.

| Gene | Element label | TSS-relative anchor(s) | Genomic anchor or form (dm6) | Use in this study and coordinate source |
|---|---|---|---|---|
| *sna* | D1 | −7,898 | chr2L:15,485,790–15,486,562; peak 15,486,150 | Productive distal opening position |
| *sna* | D2 | −7,498 | chr2L:15,485,546–15,486,154; peak 15,485,750 | Low-output distal position; Δdist0.4 is centered on this element |
| *sna* | C1 | −448 | peak chr2L:15,482,900 | Intervening CRM |
| *sna* | C2 | −1,148 | peak chr2L:15,482,200 | Intervening CRM |
| *sna* | P1 | −2,548 | chr2L:15,480,564–15,481,187; peak 15,480,800 | Later/pulse component of the proximal enhancer |
| *sna* | P2 | −1,698 | chr2L:15,479,493–15,480,205; peak 15,479,950 | Overlapping low-output, Snail-bound autoregulatory position; Δprox1.3 also contains P3 |
| *sna* | P3 | −1,498 | chr2L:15,479,399–15,480,108; peak 15,479,750 | Overlapping productive early position |
| *sna* | Δprox1.3 | −2,014 to −740 | chr2L:15,478,992–15,480,266 (1-based inclusive) | Dunipace et al. (*34*) mutant-junction flanks; P2 712/712 bp; P3 709/709 bp |
| *sna* | Δprox2.6 | −4,666 to −2,003 | chr2L:15,480,255–15,482,918 (1-based inclusive) | Dunipace et al. (*34*) mutant-junction flanks; P1 623/623 bp |
| *sna* | Δprox3.0 | −4,666 to −1,583 | chr2L:15,479,835–15,482,918 (1-based inclusive) | Dunipace et al. (*34*) mutant-junction flanks; P1 623/623 bp; P2 371/712 bp; P3 274/709 bp |
| *sna* | Δprox4.4 | −4,691 to −267 | chr2L:15,478,519–15,482,943 (1-based inclusive) | Dunipace et al. (*34*) mutant-junction flanks; P1 623/623 bp; P2 712/712 bp; P3 709/709 bp |
| *sna* | Δdist0.4 | −7,771 to −7,304 | chr2L:15,485,556–15,486,023 (1-based inclusive) | Dunipace et al. (*34*) mutant-junction flanks; D1 233/772 bp; D2 468/608 bp |
| *sna* | Δdist1.8 | −9,288 to −7,576 | chr2L:15,485,828–15,487,540 (1-based inclusive) | Dunipace et al. (*34*) mutant-junction flanks; D1 735/772 bp; D2 327/608 bp |
| *sna* | Δdist2.0 | −9,294 to −7,380 | chr2L:15,485,632–15,487,546 (1-based inclusive) | Dunipace et al. (*34*) mutant-junction flanks; D1 772/772 bp; D2 523/608 bp |
| *rho* | NEE | −1,400 and −1,050 | chr3L:1,461,970–1,462,785; anchors 1,462,400 and 1,462,550 | Exact 2–4-hour CRM union; larger centered opening used |
| *zen* | VRE | −1,457 | chr3R:6,755,493–6,755,818; peak 6,755,650 | Physical CRM boundary; REDfly interval chr3R:6,755,181–6,755,805 |
| *eve* | late CRM | −5,300 | chr2R:9,974,018 | Prespecified DAF anchor; corresponding cross-stage CRM recorded in the archived CRM catalog (*49*) |
| *eve* | stripe 2 CRM | −1,900 | chr2R:9,977,418 | Prespecified DAF anchor; corresponding cross-stage CRM recorded in the archived CRM catalog (*49*) |
| *ftz* | distal UPS | −5,248 | chr3R:6,858,898–6,859,548; peak 6,859,075 | Early program; CRM boundary used for analysis |
| *ftz* | proximal UPS | −4,098 | chr3R:6,859,998–6,860,423; peak 6,860,225 | Pulse program; CRM boundary used for analysis |
| *ftz* | distal neurogenic CRM | −2,473 | chr3R:6,861,648–6,862,148; peak 6,861,850 | Late program; CRM boundary used for analysis |
| *ftz* | proximal neurogenic CRM | −1,523 | chr3R:6,862,623–6,862,998; peak 6,862,800 | Late program; CRM boundary used for analysis |
| *ftz* | Zebra | −448 | chr3R:6,863,623–6,864,000; composite peak 6,863,875 | Early program; CRM boundary used for analysis |
| *ftz* | R | +1,400 to +1,700 | chr3R:6,865,723–6,866,023 | Gastrulation/intragenic program; prespecified DAF interval |

**Table S5.**
Prespecified locus coordinates, deletions, and analysis roles.

| Component or metric | Specification or result |
|---|---|
| Architectural factorization | Shared, position-agnostic local sequence/chromatin encoder plus promoter-specific long-range regulatory adapters; the complete predictor is intentionally promoter specific |
| One-base-resolution model input | 15 channels: definite and ambiguous factor-sized protection, nucleosome occupancy, two learnable soft NFR-width channels, eight chromatin-conditioned A/C/G/T channels, a valid-data mask, and a nucleosome-boundary-ambiguity channel |
| Chromatin-conditioned sequence input | A/C/G/T represented separately beneath definite or ambiguous factor-sized protections and throughout nucleosome-bounded regulatory gaps of at least 150 bp; overlap allowed at protected positions; sequence outside these contexts set to zero |
| Positional inputs | No genomic-coordinate, absolute-position, relative-position, distance-to-TSS, or adapter positional input |
| Shared backbone | 21-bp input projection to 128 features; 8 residual blocks with two 21-bp convolutions each; dropout 0.1; 1,021-bp receptive field; 200-bp tokens; masked mean-plus-maximum global pooling |
| Residual-block dilations | 1, 2, 2, 4, 4, 4, 4, 4 |
| NFR/sequence routing | NFR reconstruction floor 150 bp; initial soft boundary 500 bp and temperature 50 bp; separate A/C/G/T channels under factor-sized protections and across regulatory NFRs |
| Label protection | Preprocessing physically removes TSS −200 through gene end +200 before input |
| Backbone training split | 2–4 h: 1,617,028 training fibers from 793 genes; 100,000 validation fibers from 236 gene-held-out genes; zero gene overlap |
| Backbone optimization | AdamW, learning rate $5 \times 10^{-4}$, weight decay 0.01, 3,000-step warmup plus cosine decay, 6,000/4,000 train/validation token budgets, maximum 12 epochs, patience 3, seed 42 |
| Selected 2–4 h checkpoint | Epoch 2; validation mean AUC 0.64753; SHA-256 aa135d80e315087ea7f91c03da3f63e4ba060e1c6f9f240490336b71c70de3e5 |
| Adapter input geometry | Upstream: TSS −9,000 to −200 after excision. Downstream: up to 6 kb after gene end, capped at TSS +12 kb. Complete retained-window span required |
| Promoter-model input | Backbone weights held fixed during promoter-model training; 128-dimensional 200-bp tokens; tied 128→64 Q/K and 128→128 V projections; row-softmax bilinear attention; mean attended plus mean raw-token pooling; six 256→128→1 heads |
| Adapter size and optimization | 222,918 trainable parameters per promoter; AdamW, learning rate $10^{-3}$, weight decay 0.01, batch 16, maximum 15 epochs, patience 4, and gradient clipping at 1.0 |
| Stage, TSS and direction combinations | Eight combinations: primary or alternative TSS × upstream or downstream × 2–4 or 6–8 h; 5,243 promoter-specific models with at least two evaluable outputs (2–4 h: 1,273 primary upstream, 815 primary downstream, 344 alternative upstream and 167 alternative downstream; 6–8 h: 1,283, 827, 360 and 174, respectively) |
| Held-out 2–4 h primary-upstream performance | 1,273 promoters; median mean AUC 0.682, interquartile range 0.643–0.712 |
| Selected genes | *sna* 0.686; *eve* 0.700; *ftz* 0.785 |
| Adapter necessity | Matched promoter-specific minus foreign-promoter model: +0.0320 AUC; matched minus shared-backbone-only model: +0.0090; matched minus sequence-shuffled control: +0.1774 (5,313 tests) |
| Calibration | Validation-fitted isotonic calibration; test ECE 0.260 to 0.006 and Brier score 0.209 to 0.118 |
| Reproducibility records | Training and evaluation code, per-promoter model specifications, and archived output checksums are documented in the public code and source-data archives |

**Table S6.**
Model architecture and performance.

| **Stage** | **Adequately sampled CRM pairs** | **Median interaction** | **5th–95th percentile** | **Directional pair FDR super/sub** | **Pointwise 90% super/sub** | **Active additivity-compatible (equivalence subset)** |
|---|---|---|---|---|---|---|
| 2–4 h | 664 | −0.008 | −0.181 to +0.177 | 1 / 8 | 30 / 66 | 333 (*8*) |
| 6–8 h | 882 | +0.003 | −0.163 to +0.151 | 0 / 9 | 38 / 94 | 290 (*12*) |

**Table S7. Additivity-screen summary.**
The directional pair-FDR column counts all superadditive and subadditive pair calls. The ‘FDR 4’ and ‘FDR 5’ annotations in fig. S9E identify the nested complete- or attenuating-override subset within the broader subadditive class.

| Perturbation, 3–4.5 h | Pulse CRM mean deficit | Late CRM mean deficit | Pulse-minus-late difference |
| --- | --- | --- | --- |
| siGAF | 9.55 pp (n = 243) | 1.83 pp (*n* = 239) | +7.72 pp |
| Maternal *zld* heterozygote | 9.77 pp (n = 233) | 8.02 pp (*n* = 232) | +1.75 pp |

**Table S8. Opening responses of pulse and late CRMs to pioneer-factor perturbation.** Positive values indicate lower ≥175-bp opening in the perturbation. The two perturbations are not dose matched.

| Resource | Specification | Release or provenance note |
|---|---|---|
| Reference assembly and gene annotation | dm6 / BDGP Release 6; FlyBase gene models in Ensembl release 100 (BDGP6.28) | Nuclear analysis span, 137,547,960 bp. The header-stripped GTF is an exact content match to Drosophila_melanogaster.BDGP6.28.100.gtf (SHA-256 3b5316decbac4e4836cefcf59c2ef754e12ec50b0d20d694f183a23365c54b11); the chr-prefixed copy changes sequence-name prefixes only. |
| Read alignment | pbmm2 1.8.0; minimap2 2.28-r1209 and 2.30-r1287; samtools 1.19.2 and 1.21 | PacBio Hia5: pbmm2 CCS preset with coordinate sorting. ONT Hia5: minimap2 map-ont. DddB DAF-seq: minimap2 map-ont with MD tags. Exact commands are retained in BAM program headers where available. |
| FiberHMM footprint calls | FiberHMM 2.16.7 (tag v2.16.7; commit d4607e47e33fddc7933a5103560f34517caabe01) | Hia5/PacBio, Hia5/Nanopore and DddB/Nanopore DAF models were selected explicitly. Nucleosome and factor-sized-footprint recall were applied to every call set analyzed here; duplicated PacBio movie m84046_250513_034312_s1 was excluded. The manuscript FiberHMM source snapshot is archived in Zenodo (*47*), with the live repository linked there. |
| FiberBrowser | FiberBrowser 2.15.4 | Genome-browser snapshots were rendered from FiberHMM annotations. The manuscript FiberBrowser source snapshot is archived in Zenodo (*48*). |
| DAF-seq molecule processing | Deamination-fingerprint deduplication and alignment-based callable-molecule reconstruction | Targeted DAF-seq reads were deduplicated at Jaccard ≥0.95 before quantitative extraction. Callable molecules were reconstructed from alignments; absence from a feature-only BigBed was not treated as closure. |
| CRM catalog | Cross-stage nonredundant catalog | 12,121 total cross-stage units (SHA-256 3f81f556a4113937ac43398fabeeff8bc6b15fdc7fb717dd3faa1de44b0002db); 6,942 have a widest stage-specific boundary of at least 300 bp and are eligible for resolved-unit regulatory analyses. |
| REDfly comparison | Canonical in vivo and broader evidence hierarchy | The exact database export date and comparison table are included in the archived source data (*49*). |
| Motif resource | JASPAR 2022 | dm6 motif track; matched Trl/GAF and vfl/Zld MOODS scans on both strands at $P \leq 10{-}3$; dl calls used the stated score threshold. |
| Statistical environment | Python 3.10.13; NumPy 1.26.4; pandas 2.3.3; SciPy 1.15.3; Matplotlib 3.10.0; scikit-learn 1.6.1; statsmodels 0.14.5 | Versions report the environment used to rebuild this supplement on 18 August 2026. |
| FiberCNN checkpoint | 2–4-h primary-upstream selected checkpoint | Epoch 2; validation mean AUC 0.64753; SHA-256 aa135d80e315087ea7f91c03da3f63e4ba060e1c6f9f240490336b71c70de3e5. The checkpoint and FiberCNN source are archived in Zenodo (*46*). |
| Code and data | GEO and SRA (accessions provided upon publication); Zenodo records (*46–49*) | FiberCNN, FiberHMM, FiberBrowser, model checkpoints, and analysis and figure-generation code and source data are archived in Zenodo (*46–49*). |
| Embryonic sci-ATAC atlas | Calderon et al. (*55*); GEO GSE190149 | Whole-embryo CPM values over 110,185 peaks. The 2–4-h window was used as the external early state and the mean of 4–8- and 6–10-h windows as the external late state. |
| nc14 single-cell RNA matrix | Karaiskos et al. (*51*); dge_normalized.txt, 1,297 cells | Fraction of stage-6/nc14 cells with normalized expression above zero was used to rank genes. |
| Early embryonic PRO-seq | Hunt et al. (*22*); GSM6454906 / GSE211180 | WT naive-embryo forward- and reverse-strand tracks were summarized in the matched gene-body window. |
| CRM-validation histone tracks | Brennan et al. (*6*), GSE218852; Li et al. (*12*), GSE58935 | Brennan H3K4me1 and Li cycle-14a H3K27ac/H3K4me3 tracks were used in fig. S8E. |
| CRM-validation CBP/Nejire track | Koenecke et al. (*53*); GSE68983 | Embryonic CBP/Nejire ChIP-seq track used in fig. S8E. |
| Evolutionary conservation | UCSC dm6 phastCons124way | Basewise conservation track used in fig. S8E; exact source-track identifier and checksum accompany the source data. |
| Candidate-TF occupancy support | GSE152770; GSE30757; GSE83305; GSE304853; ENCODE accessions listed at right | Stage- or cell-matched support: GAF GSE152770, Zld GSE30757, Grh GSE83305/GSM2199199 (*57*), and S2 Srp GSE304853 (*58*). Broad embryonic support: GATAe/grn ENCSR625TAA and ENCSR909QHH, Stat92E ENCSR290OJD, Pnt ENCSR997UIM, Hr78 ENCSR280XEB, and Mad ENCSR875FVD. |
| CAGE TSS refinement | Schor et al. (*50*); E-MTAB-4787; strand-specific nc14 CAGE | The strongest local CAGE peak within ±100 bp of annotated transcript starts was used when summed signal was at least 1; the annotated TSS was retained otherwise. |
| Pioneer-factor ChIP-NEXUS | Brennan et al. (*6*); GSE218852; stage-5 GAF and Zld | For each factor, absolute signal over the CRM was adjusted by equal-width flanks placed 1 kb from each edge and percentile ranked across eligible CRMs. |

**Table S9.**
Software, reference resources, and reproducibility identifiers.
